# Experienced Meditators Show Multifaceted Attention-Related Differences in Neural Activity

**DOI:** 10.1101/2023.02.10.527999

**Authors:** Neil W Bailey, Oliver Baell, Jake Elijah Payne, Gregory Humble, Harry Geddes, Isabella Cahill, Aron T Hill, Sung Wook Chung, Melanie Emonson, Oscar W Murphy, Paul B Fitzgerald

## Abstract

**Objectives:** Mindfulness meditation (MM) is suggested to improve attention. Research has explored this using the ‘attentional-blink’ (AB) task, where stimuli are rapidly presented, and a second target stimulus (T2) is often missed if presented ∼300ms after an initial target stimulus (T1). This research showed improved task-accuracy and altered neural activity after an intensive 3-month MM retreat. We tested whether these results replicated in a community sample of typical meditators.

**Methods:** Thirty-one mindfulness meditators and 30 non-meditators completed an AB task while electroencephalography (EEG) was recorded. Between-group comparisons were made for task-accuracy, event-related potential activity (posterior-N2 and P3b), theta and alpha oscillatory phase synchronisation to stimuli presentation, and alpha-power. Primary aims examined effects within time windows reported by previous research. Additional exploratory aims assessed effects across broader time windows.

**Results:** No differences were detected in task-accuracy or neural activity within our primary hypotheses. However, exploratory analyses showed posterior-N2 and theta phase synchronisation effects indicating meditators prioritised attending to T2 stimuli (p < 0.01). Meditators also showed more alpha-phase synchronisation, and lower alpha-power when processing T2 stimuli (p < 0.025).

**Conclusions:** Our results showed multiple differences in neural activity that suggested enhanced attention in meditators. The neural activity patterns in meditators aligned with theoretical perspectives on activity associated with enhanced cognitive performance. These include enhanced alpha ‘gating’ mechanisms, increased oscillatory synchronisation to stimuli, and more equal allocation of neural activity across stimuli. However, meditators did not show higher task-accuracy, nor did effects align with our primary hypotheses or previous research.

**Preregistration:** This study was not preregistered.

## Introduction

Mindfulness meditation (MM) is a broad term to describe meditation practices that attend to aspects of the present moment without judgment (e.g., the breath, bodily sensations, awareness) (Crane et al., 2017; Van Dam et al., 2018). Over recent decades, MM has been incorporated into mindfulness-based interventions (MBIs) to alleviate symptoms of depression, pain, and addiction (Hayes, 2012; Kuyken et al., 2008). Our understanding of the mechanisms of MM is rapidly improving, with studies replicating mechanistic relationships between mindful attention, emotional regulation and well-being outcomes with moderate consistency (Britton et al., 2018; Chambers et al., 2009; Kiken et al., 2015). However, there are an array of theoretical perspectives regarding the neurophysiological mechanisms that underpin the effects of MM, and not enough empirical evidence to draw strong, comprehensive, or specific conclusions about the accuracy of the proposed mechanisms (Hölzel et al., 2011; Tang et al., 2015; Van Dam et al., 2018). A better mechanistic understanding of MM is therefore required. Specifically, there is a need to elucidate the neurophysiological changes that underlie the benefits of the practice to well-being. This might allow the design of MM interventions with enhanced efficacy by specifically targeting the effective mechanism.

One promising psychological mechanism that may underlie the effects of MM is improved attentional function (Kiken et al., 2015; Tang et al., 2007). MM’s ability to improve attention is supported by controlled behavioural studies showing MM practice increased sustained and executive attention (Jha et al., 2007; Lutz et al., 2009; Slagter et al., 2009; Tang et al., 2007) and improved performance on various attentional tasks (Atchley et al., 2016; Bailey et al., 2019; Bailey et al., 2022; Van Dam et al., 2018). One sophisticated approach to measure MM-related changes in attention is to examine the limited temporal capacity of attention using the attentional blink (AB) phenomenon (Martens & Wyble, 2010; Shapiro et al., 1997). In a typical AB task, individuals are presented with a rapid stream of ∼20 distractor stimuli. Within that rapid stream of stimuli, two targets (T1 and T2) are presented in close temporal succession, with T1 typically appearing randomly after 2-8 stimuli have already been presented and the T2 stimulus appearing 200 to 700ms after T1 (Ward et al., 1996). The AB phenomenon refers to a reduction in accuracy at recalling the T2 stimulus when it is presented within 200-500ms after T1, with AB trials presenting T2 stimuli at this brief delay often referred to as a “short interval” attention blink trials (Shapiro et al., 1997). A number of cognitive models have been proposed to explain the AB phenomenon (for a review, see Martens and Wyble (2010)). Capacity-based models suggest competition between stimuli for attentional resources, so T1 induces a drain on limited attentional resources and insufficient attentional resources are available to successfully process T2 (Potter et al., 1998; Shapiro et al., 1997). In contrast, selection-based models emphasise the role of attentional control, where the magnitude of an individual’s AB is affected by the extent to which distracting information is suppressed (Di Lollo et al., 2005; Olivers & Meeter, 2008). However, it is worth noting that thus far, evidence supporting one analytical model of the AB phenomenon does not necessarily negate the mechanisms and functional processes proposed by another (further discussion of this point is available in the supplementary materials section 1).

The neurophysiological mechanisms of AB phenomena have been explored using EEG (Slagter et al., 2007; Vogel et al., 1998). This research has focused on an ERP known as the P3b, which is a positive voltage occurring maximally in parietal electrodes around 350 to 600ms following stimulus presentation, and which has been associated with voluntary attentional focus (Falkenstein et al., 1993; Falkenstein et al., 1991). Research has found the P3b time-locked to the second target stimuli to be entirely suppressed in trials in which the second target is ‘blinked’ (not consciously perceived) and ultimately not recalled (Dell’Acqua et al., 2015). A reduced AB effect (i.e., increased ability to detect T2 stimuli) has also been associated with an earlier onset and smaller amplitude of the T1-induced P3b, suggesting that when less neural activity is devoted to the T1 stimulus, more neural resources are available to detect and encode the T2 stimulus (Sergent et al., 2005; Slagter et al., 2007). In addition to the P3b AB effect, research has also suggested that short interval AB trials reduce the amplitude of the visual processing related posterior-N2, an ERP peaking approximately 200ms after stimuli presentation, with posterior-maximal negative voltages (Zivony et al., 2018). This is thought to reflect the lack of engagement of attention processes time-locked to T2 stimuli (Zivony et al., 2018). In addition to the ERP AB findings, research has suggested that theta oscillations (rhythmic EEG activity occurring between 4-8Hz) are related to a range of cognitive processes, including attention (Mizuhara & Yamaguchi, 2007). A positive relationship between the successful detection of AB targets and theta phase synchronisation (TPS) to the onset of AB target presentation has also been identified (Slagter et al., 2009). Finally, decreased synchronisation of alpha (8-13Hz) oscillations to the onset of the distractor stimuli presentation (which are presented prior to T1) and increased alpha-power just prior to T1 stimuli presentation has also been associated with improved performance in the AB task (Slagter et al., 2009). As alpha oscillations have been linked with functional inhibition of brain regions (Klimesch, 2012), it is possible that desynchronisation of alpha oscillations around the stimulus and increased alpha-power just prior to the target stimulus onset inhibits processing of the distractors, then releases any inhibitory processes ongoing on brain regions responsible for processing the target AB stimuli, resulting in better AB performance.

Perhaps unsurprisingly, MM training and experience have been shown to reduce the AB phenomenon (Slagter et al., 2007; van Leeuwen et al., 2009; Wang et al., 2021). However, to date, only Slagter et al. (2007) have measured neural activity while meditators perform the AB task. They compared EEG activity from non-meditator controls and experienced mindfulness meditators (with an average of 2,967 hours of meditation experience) before and after the experienced meditators underwent an intensive 3-month meditation retreat, and the non-meditators practised MM for 20 minutes per day for one week. Following the retreat, the experienced meditators were better at identifying the T2 AB stimuli compared to the controls (demonstrating a reduced AB effect) (Slagter et al., 2007; Slagter et al., 2009). The improved T2 detection was correlated with a reduced P3b following T1 stimuli, as well as increased T2-locked TPS (Slagter et al., 2007; Slagter et al., 2009). Slagter et al. (2007) suggested that the reduction in T1-elicited P3b in meditators may reflect a reduced propensity to mentally ‘cling’ to a target, whereas the elevated TPS may reflect an increased capacity to process experience from moment to moment. They also found a reduction in alpha phase synchronisation (APS) to the distractor stimuli (prior to the onset of T1) in meditators, potentially implicating the release of alpha inhibiting the processing of distractor stimuli before T1 presentation (Slagter et al., 2009). Notably, these findings were only after an intensive 3-month retreat and it is unclear if more typical daily MM practice will produce similar effects. Exploring a community sample of MM may provide findings that are more generalisable to a typical (and increasingly popular) MM practice (Cramer et al., 2016). Additionally, while Slagter et al. (2007) have been cited over 1000 times, no replications of their study have been attempted.

Given this background, the primary aim of the study was to compare brain activity related to the AB phenomenon (P3b, TPS, APS and alpha-power) between a cross-sectional sample of experienced community meditators and healthy control non-meditators. The present study also utilised advanced EEG analysis methods, which can separately detect differences in overall neural response strength and differences in the distribution of brain activity. Following the research by Slagter et al. (2007, 2009), our primary hypotheses were that: PH1) compared to non-meditator controls, meditators would show a smaller allocation of attention-related neural resources to T1 as indexed by a lower amplitude T1-elicited P3b during short interval trials; PH2) meditators would show more consistency in the timing of theta oscillatory neural activity (higher TPS) in response to T2 during short interval trials but not long interval trials, indexed by higher T2-locked TPS values; PH3) the meditators would show greater alpha-power around stimuli presentation in short and long interval T1 trials compared to controls. Finally, the AB task presented stimuli every 100ms (at 10Hz), which is within the alpha frequency. This is likely to produce alpha synchronisation to the task stimuli, an effect that may be modified in the meditation group, which has undergone considerable training in an attention-based practice. Slagter et al. (2009) reported a reduction in APS during the presentation of the distractor stimuli prior to T1 presentation after the meditation retreat (in contrast to the increased alpha-power). As such, we had one further primary hypothesis: PH4) APS would be reduced in the meditation group during the presentation of the distractor stimuli prior to T1 stimuli. Additionally, while we tested these primary hypotheses within the time windows reported by Slagter et al. (2007, 2009), to ensure we did not miss significant effects that appeared outside these specific windows, we conducted additional exploratory analyses for the ERP, TPS, APS, and alpha-power variables which included all time points in the EEG epochs for each of these measures (exploratory hypotheses are explained below), while employing data-driven multiple comparison controls.

Additionally, since behavioural research using a cross-sectional design has previously shown that meditators show a reduced AB effect compared to non-meditator controls, we had a non-primary replication hypothesis, RH1) that our meditation group would show a reduced AB effect as indicated by meditators showing higher accuracy than controls in short interval T2 trials. Further, while Slagter et al. (2007) focused on the P3b in response to T1 only, our view is that it is sensible to hypothesise that EH1) ERPs to T2 would be increased in meditators, or EH2) the relationship between ERP amplitude to T1 and T2 is different in meditators, perhaps reflecting an increased ability to attend to the T2 stimulus as a result of a reduced focus on the T1 stimuli. Additionally, since previous research has not examined potential differences in the topographical distribution of neural activity in meditators during the AB task, four non-directional exploratory hypotheses were that: EH3) meditators would show differences in the scalp distribution of ERPs, EH4) meditators would show differences in the scalp distribution of TPS, EH5) meditators would show differences in the scalp distribution of alpha-power, and EH6) meditators would show differences in the scalp distribution of APS.

## Method

### Participants

A sample of 39 experienced community meditators and 36 healthy control non-meditators were recruited after responding via phone call or email to community advertising at universities, meditation organisations, and on social media. To meet the eligibility criteria for classification as an experienced meditator, participants were required to have had at least two years of meditation experience and have practised meditation for a minimum of two hours per week over the last three months. Meditation was defined by Kabat-Zinn’s definition: “paying attention in a particular way: on purpose, in the present moment, and nonjudgmentally” (Kabat-Zinn, 1994). This definition included participants who practice both open monitoring meditation, which involves simple awareness without a specific focus besides awareness itself and focused attention meditation, which involves deliberate attention on a specific object, such as the breath (Cahn & Polich, 2009; Lutz et al., 2008). Trained MM researchers (BLINDED FOR REVIEW) interviewed and screened participants to ensure the participants’ practices fit the criteria, and screening uncertainties were resolved through discussion and consensus between the principal investigator (BLINDED FOR REVIEW) and one other researcher. Eligibility as a non-meditator control required participants to have less than two hours of lifetime meditation experience.

Participants were considered ineligible to participate if they were currently taking psychoactive medication; had experienced brain injury; had previously been diagnosed with a psychiatric or neurological condition; or met the criteria for any drug, alcohol or neuropsychiatric disorders as measured by the Mini International Neuropsychiatric Interview (MINI) (Sheehan et al., 1998). Participants who scored above the moderate range (greater than 25) in the Beck Anxiety Inventory (BAI) (Beck et al., 1988) or the mild range (greater than 19) in the Beck Depression Inventory-II (BDI-II) (Beck et al., 1961) were also excluded.

Ethical approval of the study was provided by the (BLINDED FOR REVIEW). All participants provided written informed consent prior to participation in the study. Before participants underwent EEG recording, participants provided their gender, age, years of education, and meditation experience (total years of practice, frequency of practice, and the usual length of a meditation session). Participants also completed the Five Facet Mindfulness Questionnaire (FFMQ) (Baer et al., 2006), BAI, and BDI-II. Two controls were excluded from the study due to scoring above the moderate anxiety range on the BAI. Two controls and one meditator were excluded after scoring in the mild depression range on the BDI-II. Another control was excluded after revealing a history of meditation. Two meditators were excluded due to a previous history of seizures, substance abuse or mental illness, and another three were excluded from the analysis due to not completing the AB task. Lastly, two meditators and one control were excluded from the study as their performance of the AB task was near chance.

The final sample included 31 meditators aged between 20 and 64 years and 30 healthy controls aged between 20 and 60. The two groups did not differ in any demographic or self-report measure except for the FFMQ score (all p > 0.05, except for the FFMQ, where p < 0.001). Table 1 summarises all measures (note that one participant did not complete the BAI, and another did not complete the FFMQ, so their data were excluded from those measures). The final sample of meditators had a mean of 6.44 (SD = 4.25) years of meditation experience, 7.65 (SD = 2.21) hours of current practice per week and a mean of 55.65 (SD = 44.90) minutes of meditating per session.

**Table 1.**
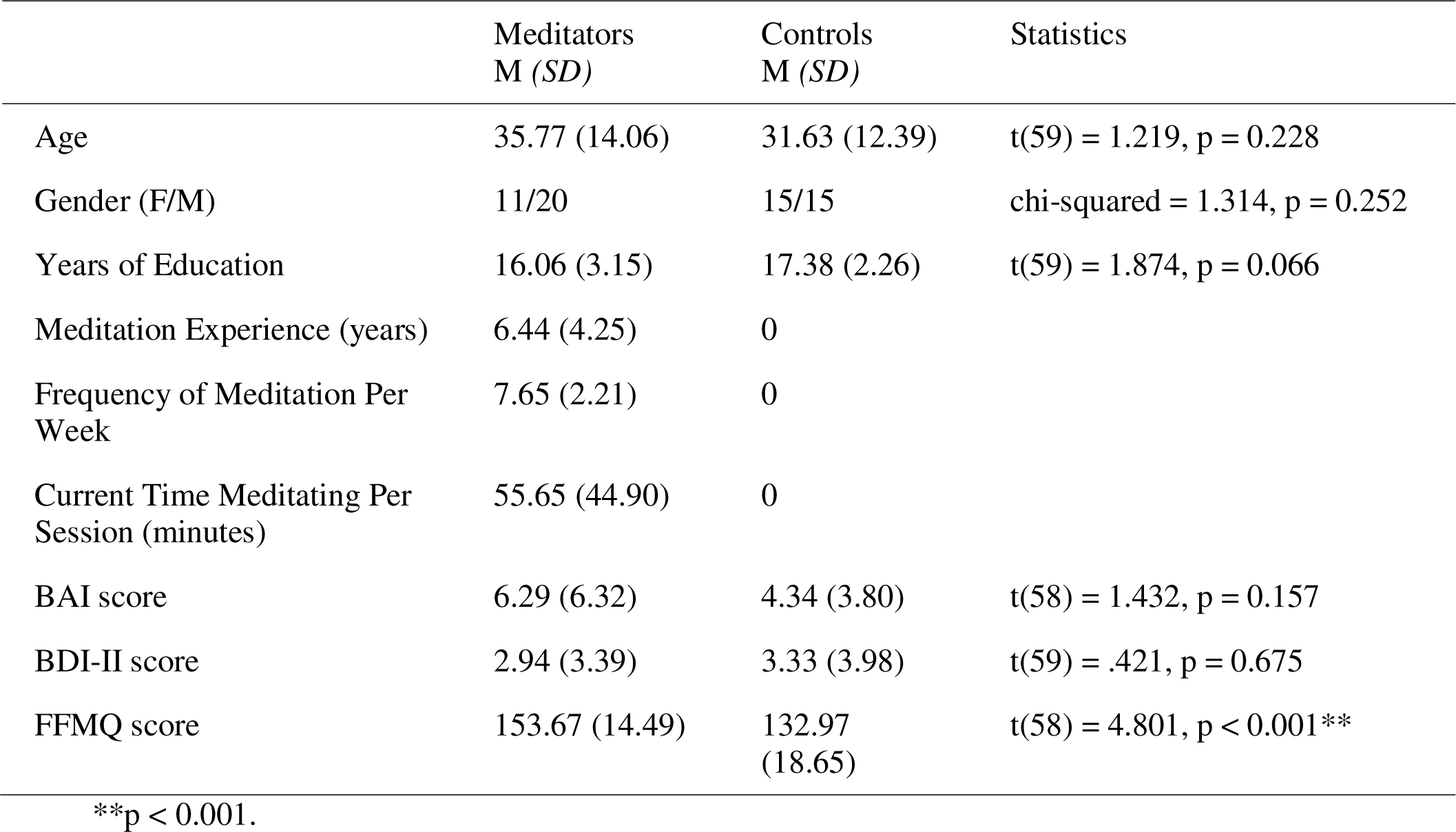
Demographic and self-report means (M), standard deviations (SD), and statistics. BAI = Beck Anxiety Inventory, BDI-II = Beck Depression Inventory II, FFMQ = Five Facet Mindfulness Questionnaire.

### Procedure

Participants first performed a Go/Nogo task (data not yet published) and an auditory oddball task [BLINDED FOR REVIEW], followed by the AB task. The AB task was a replication of the task used by Slagter et al. (2007). The task involved 12 practice trials followed by four blocks of 90 trials where the participants viewed a stream of 19 stimuli (letters and numbers) presented for 66ms, with a 33ms blank screen between each stimulus. Before the task began, participants were instructed that there could be one or two numbers in each trial and were to enter the number/s they observed on a number pad once each trial ended. Each new trial began after the participant pressed the Enter key to continue, and participants were offered the option of a short break between each of the four blocks. T1 occurred at a random position from 3 to 9 in the stream (after 2-8 distractor stimuli had already been presented). In trials with two numbers, T2 could occur either 300ms (short interval) or 700ms (long interval) after T1. Each block contained 54 short interval trials, 18 long interval trials, and 18 T1-only trials (where no T2 stimulus was presented). The order of the trials within each block was randomised. The number of correct trials (both T1 and T2 correct), trials where T1 was incorrect, and trials where T2 was incorrect were recorded for each participant. The total task time was approximately 45 minutes. After the AB task, participants were administered transcranial magnetic stimulation concurrent with EEG (data not yet published).

### Measures

#### Electrophysiological Recording and Pre-Processing

64-channel EEG data were recorded continuously during the tasks using a Quick-Cap containing Ag/AgCl electrodes and SynAmps 2 amplifier (Compumedics, Melbourne, Australia). Data were recorded by Neuroscan Aquire software, with samples obtained at 1000Hz and an online bandpass filter from 0.05 to 200Hz (24dB/octave roll-off). Each electrode was connected to a reference electrode positioned between CPz and Cz. Prior to the start of the recording, all electrode impedances were reduced to <5kΩ.

EEG recordings were pre-processed offline in MATLAB R2018b (The Mathworks, Inc.) using the RELAX EEG cleaning pipeline (Bailey et al., 2022a; Bailey et al., 2022b), which calls EEGLAB (Delorme & Makeig, 2004) and fieldtrip functions (Oostenveld et al., 2011). Within the RELAX pipeline, data were first bandpass filtered with a fourth-order Butterworth filter from 0.25 to 80Hz and bandstop filtered from 47 to 53Hz to reduce the line noise. Next, multiple default RELAX approaches were used to reject extreme outlying channels (Bailey et al., 2022a; Bailey et al., 2022b; Bigdely-Shamlo et al., 2015), followed by the marking of extreme outlying EEG periods for exclusion from the Multiple Wiener Filter cleaning and deletion before independent component analysis (see Bailey et al. 2022a for details). Three sequential Multiple Wiener Filters were used to reduce 1) muscle activity (Fitzgibbon et al., 2016), 2) eye blinks, then 3) horizontal eye movement and electrode drift (Somers et al., 2018). Finally, data were re-referenced to the robust average reference (Bigdely-Shamlo et al., 2015), and the remaining artifacts were cleaned using wavelet-enhanced independent component analysis (ICA) (Castellanos & Makarov, 2006) to reduce artifactual components identified by ICLabel (Pion-Tonachini et al., 2019) after ICA decomposition using cudaICA (Raimondo et al., 2012). Full details of the pre-processing pipeline are available in Bailey et al. (2022a) and Bailey et al. (2022b).

After cleaning, EEG activity was epoched to the onset of the AB task stimuli from -200 to 1000ms surrounding the T1 or T2 stimuli for ERP analysis and from -2000 to 2000ms for oscillation analyses. The fieldtrip ‘ft_freqanalysis’ function was used with Morlet Wavelet analysis settings and a cycle width of 5 to compute frequency power.

ERP data were baseline corrected using the baseline subtraction method to the average activity in the -200 to 0ms period prior to target stimulus onset, as per the methods of Slagter et al. (2007). To test our hypothesis PH1 for the P3b ERP, we averaged data within the 350 to 600ms time window following the stimuli.

TPS and APS were quantified through the calculation of a phase-locking factor (PLF) value within the theta range (4 to 8.5Hz) and alpha range (8.5 to 15Hz, in replication of Slagter et al. 2009) (Lachaux et al., 1999; Ueno et al., 2009). PLF values range from 0 to 1, where 1 represents perfectly correlated phase differences between trials, and 0 represents completely uncorrelated phase differences (Ueno et al., 2009; Varela et al., 2001). The methods for this computation are described in more detail in the supplementary materials (section 2b). To test hypothesis PH2, TPS data were averaged within the 121 to 501ms window after T2 stimuli. To test hypothesis PH4, APS data were averaged within the -414 to -214ms window prior to T1 stimuli.

For alpha-power frequency power analyses, trials were baseline corrected to oscillatory power across the entire epoch (in replication of Slagter et al., 2009). While this means a potential signal reduction in potential “active” periods (as the data from those periods is contained within the baseline subtraction), this approach prevents spurious conclusions about differences in active periods being, in fact, driven by an arbitrarily selected baseline period. As such, significant differences at any time point in the epoch reflect an increase or decrease of oscillatory power at those time points relative to the ongoing oscillatory power across the entire epoch. Baseline correction of frequency power data was performed using the relative method ([data – mean baseline activity] / mean baseline activity). To test hypothesis PH3, alpha power was averaged within the -31 to 160ms time window following T1. Only epochs from correctly responded to target stimuli were used in the EEG analysis (for epochs locked to T1, this meant trials where T1 was responded to correctly, while for T2 locked epochs, this meant trials where participants correctly identified both T1 and T2 stimuli). Each condition (short vs long interval and T1 vs T2 stimuli) was averaged separately within each participant for ERP and oscillation analyses.

### Data Analysis

#### EEG Comparisons

EEG data comparisons of ERPs, TPS, alpha-power, and APS, between meditators and non-meditators, were performed using the Randomised Graphical User Interface (RAGU) method (Koenig et al., 2011). RAGU compares scalp field differences over all epoch time points and electrodes using rank order randomisation statistics with no preliminary assumptions about time windows and electrodes to analyse (Koenig et al., 2011). Prior to conducting primary tests, a Topographical Consistency Test (TCT) was conducted to confirm the consistent distribution of scalp activity within each group and condition. A significant TCT result suggests that potential between-group differences in the Global Field Power (GFP) and Topographic Analysis of Variance (TANOVA) tests (described shortly) are due to real group differences instead of variation within one of the groups (Koenig & Melie-garcía, 2010). RAGU allows for comparisons of global neural response strength (independent of the distribution of activity) with the GFP test. The GFP is an index of the total voltage differences across all channels, regardless of the specific locations of the activity; it is equivalent to the standard deviation across all channels at each time point (Habermann et al., 2018). The GFP test compares differences between groups or conditions from the real data against randomised permutation data to identify specific time periods following a stimuli where groups or conditions significantly differed in neural response strength. RAGU also allows for comparisons of the distribution of neural activity with the TANOVA (with the recommended L2 normalisation of the amplitude of neural activity which transforms data for such that the overall GFP = 1 within each individual, providing distribution comparisons that are independent of differences in global amplitude). Note that there are currently no Bayesian statistical approaches analogous to the TANOVA.

TPS, alpha-power, and APS values were compared with Root Mean Square (RMS) and TANOVA tests (to separately compare overall neural response strength and distribution of neural activity, respectively). The RMS is computed in the same manner as the GFP, but without implementing an average re-referencing across the data prior to its computation. This is the recommended approach when oscillatory power or phase synchronisation comparisons are computed with RAGU, as the average reference was computed prior to the oscillation measurement transforms. As such, the RMS test is a comparison of the RMS between groups rather than the GFP, a measure which is a valid indicator of neural response strength in the power or phase synchronisation domain (Habermann et al., 2018). In other respects, the statistic used to compare RMS between groups is identical to the GFP test described in the previous paragraph.

RAGU controls for multiple comparisons in space by using only a single value representing all electrodes for the GFP/RMS and TANOVA tests (the GFP/RMS value for the GFP/RMS test and the global dissimilarity value for the TANOVA). RAGU also controls for multiple comparisons across time points in the epoch using global duration statistics which calculate the periods of significant effects within the epoch that are longer than 95% of significant effects in the randomised data with the alpha level at 0.05 (Koenig et al., 2011). However, because the computation of measures of oscillatory power or phase consistency elicits a dependence in values across neighbouring timepoints, RAGU’s global duration control method is only appropriate for ERP analyses. For our oscillatory power and phase measures we implemented the same duration controls as Slagter et al. (2009). Because our primary hypotheses were obtained from Slagter et al. (2007, 2009), we averaged data within specific windows of interest for our primary analyses. However, to explore potential effects outside of these windows, we also used RAGU for whole epoch analyses (from -100 to 800ms for ERPs and from -500 to 1500ms for oscillatory analyses), with multiple comparison controls implemented using the global duration statistics. The recommended 5000 randomisation permutations were conducted with an alpha of *p* = 0.05. For more in-depth information about RAGU and its analyses, please refer to Koenig et al. (2011), Koenig and Melie-garcía (2010) and Habermann et al. (2018). The p-values from our primary hypotheses (with data averaged within a priori hypothesised time windows of interest) were submitted to False Discovery Rate (FDR) multiple comparison controls (Benjamini & Hochberg, 2000) to control for experiment-wise multiple comparisons (referred to as FDR-p). For the sake of brevity, only main effects and interactions involving group are reported in the manuscript, while other results of interest are reported in the supplementary materials (section 3). For brevity, the full details of all statistical analyses are reported in the supplementary materials (section 2). However, we note here that some time windows of interest occurred prior to the presentation of T1 stimuli, in line with Slagter et al. (2009). These time windows were analysed as the results from Slagter et al. (2009) suggested differences in the meditation group in the synchronisation of neural activity to the distractor stimuli that were presented prior to T1, perhaps suggesting less reactivity to those stimuli in preparation for processing the target.

To test our hypotheses for ERPs (PH1, EH1, EH2, and EH3), global field power (GFP) and topographical analysis of variance (TANOVA) tests were averaged between 350 to 600ms (P3b period) (Polich, 2007) after T1 onset to make direct comparisons with Slagter et al. (2007). For this averaged activity, GFP and TANOVA tests were used to conduct repeated measures ANOVA design statistics, examining 2 groups (meditators vs controls) x 2 conditions (short and long interval). To test our exploratory hypotheses that differences might be present outside of this specific time window or might be present following T2 (EH1, EH2, and E3), GFP and TANOVA tests were used to conduct the repeated measures ANOVA design statistics, examining 2 groups (meditators vs controls) x 2 conditions (short and long interval) x 2 targets (T1 and T2) for event-related potential (ERP) data across the entire -100 to 800ms interval after T1 onset.

To test our hypotheses for TPS (PH2 and EH4), we compared TPS between the groups, root mean squared (RMS) and TANOVA tests were used to conduct repeated measures ANOVA design, examining 2 group (meditators vs controls) x 2 condition (short and long interval) comparisons for TPS data surrounding T2 onset. To make comparisons with Slagter et al. (2009), RMS and TANOVA tests were averaged within the 121 to 501ms window (where Slagter et al. 2009 detected an effect that was maximal at electrodes FC6 and Fz) and the 309 to 558ms window (where Slagter et al. 2009 detected an effect that was maximal at electrode T8) after the T2 stimuli. An additional exploratory analysis was performed, including T1 stimuli in a repeated measures ANOVA design examining 2 groups (meditators vs controls) x 2 conditions (short and long interval) x 2 conditions (T1 and T2) for TPS data from -500 to 1500ms around the stimuli to determine if any effects were missed by the analysis focused only on T2.

To test our hypotheses related to alpha-power and APS (PH3, PH4, EH5 and EH6) RMS and TANOVA tests were used to conduct repeated measures ANOVA design comparisons of alpha-power and APS (separately), examining 2 group (meditators vs controls) x 2 condition (short and long interval) comparisons for data averaged within a -31 to 160ms period for alpha-power and averaged within a -414 to 214ms period for APS. Similar to the ERPs and TPS tests, we also performed a whole epoch analysis from -500 to 1500ms surrounding T1 onset to test for effects outside those reported by Slagter et al. (2009).

#### Single Electrode Replication Comparisons

In addition to the RAGU analysis, traditional single-electrode comparisons were conducted for comparison with previous research, using time windows and electrodes that showed significant results in comparisons by Slagter et al. (2007, 2009). Methods and results for these comparisons are reported in the supplementary materials (sections 2 and 3 respectively).

#### Behavioural and Demographic Comparisons

Between-group comparisons of the demographic and behavioural data were performed using SPSS v23 or the robust statistics WRS2 package from R where parametric assumptions were not met (Field and Wilcox, 2017). Independent samples t-tests compared age, BAI, BDI-II, FFMQ, and years of education. A three-way repeated measures ANOVA was planned to analyse behavioural data. Interval (short or long) and Target (T1 or T2) were within-subjects factors and Group (meditators vs controls) the between-subjects factor. The dependent variable was AB accuracy, defined as the percentage of correctly responded to trials (T1 and T2 identified correctly). This tested hypothesis RH1, with post-hoc tests planned to assess the specific hypothesis that meditators showed a reduced AB effect (defined by increased short interval T2 accuracy) if an interaction between Group, Target and Interval were present. Where possible, Bayesian analyses were also performed using JASP (Love et al., 2019) to provide the strength of evidence for either the null or alternative hypotheses (for all of the behavioural, demographic, and EEG comparisons), and a small number of follow up exploratory linear mixed models were used to test our explanations for significant results (described in full in the supplementary materials, section 3).

## Results

### ERP Comparisons

To test our first primary hypothesis (PH1) that meditators would show a smaller allocation of attentional-related neural resources to T1, reflected by a lower amplitude of the P3b neural response strength to T1 stimuli in meditators compared to controls, the GFP test was performed on the P3b time window (from 350 to 600ms following T1, consistent with Slagter et al., 2007). No difference was detected for the main effect of Group in GFP averaged across the P3b period (p = 0.798, FDR-p = 0.798, η_p_^2^= 0.001, see Table 2 and Figure 1), nor was there a significant interaction between Group and Interval (p = 0.732, η_p_^2^= 0.004). To test the strength of evidence for the null hypothesis, averaged P3b GFP values from within the time window of interest (350 to 600ms) were submitted to Bayesian statistics. This result showed that the null hypothesis was more likely than the alternative hypothesis for both the Group factor and the interaction between Group and Interval. Comparing models including Group and a Group by Interval interaction to the model only including Interval provided BF01 = 6.520, while comparing the main effect of Group independently to equivalent models stripped of the Group effect and excluding higher-order interactions, BFexcl = 1.835, and for the interaction between Group and Interval, BFexcl = 3.553. Our single electrode analyses, which focused on time windows and electrodes reported to be significant by Slagter et al. (2007), showed similarly null results (supplementary materials section 3b).

**Table 2:**
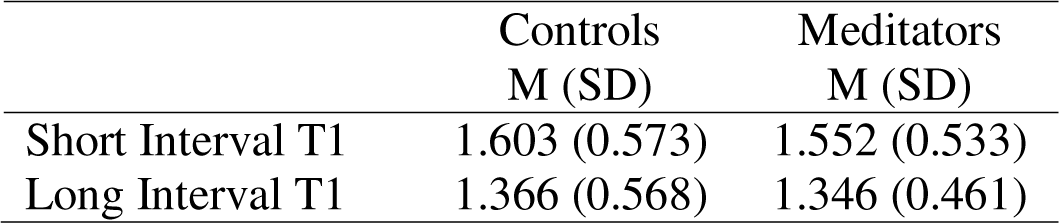
Global field potential (GFP) values averaged across the P3b period of interest.

**Figure 1.**
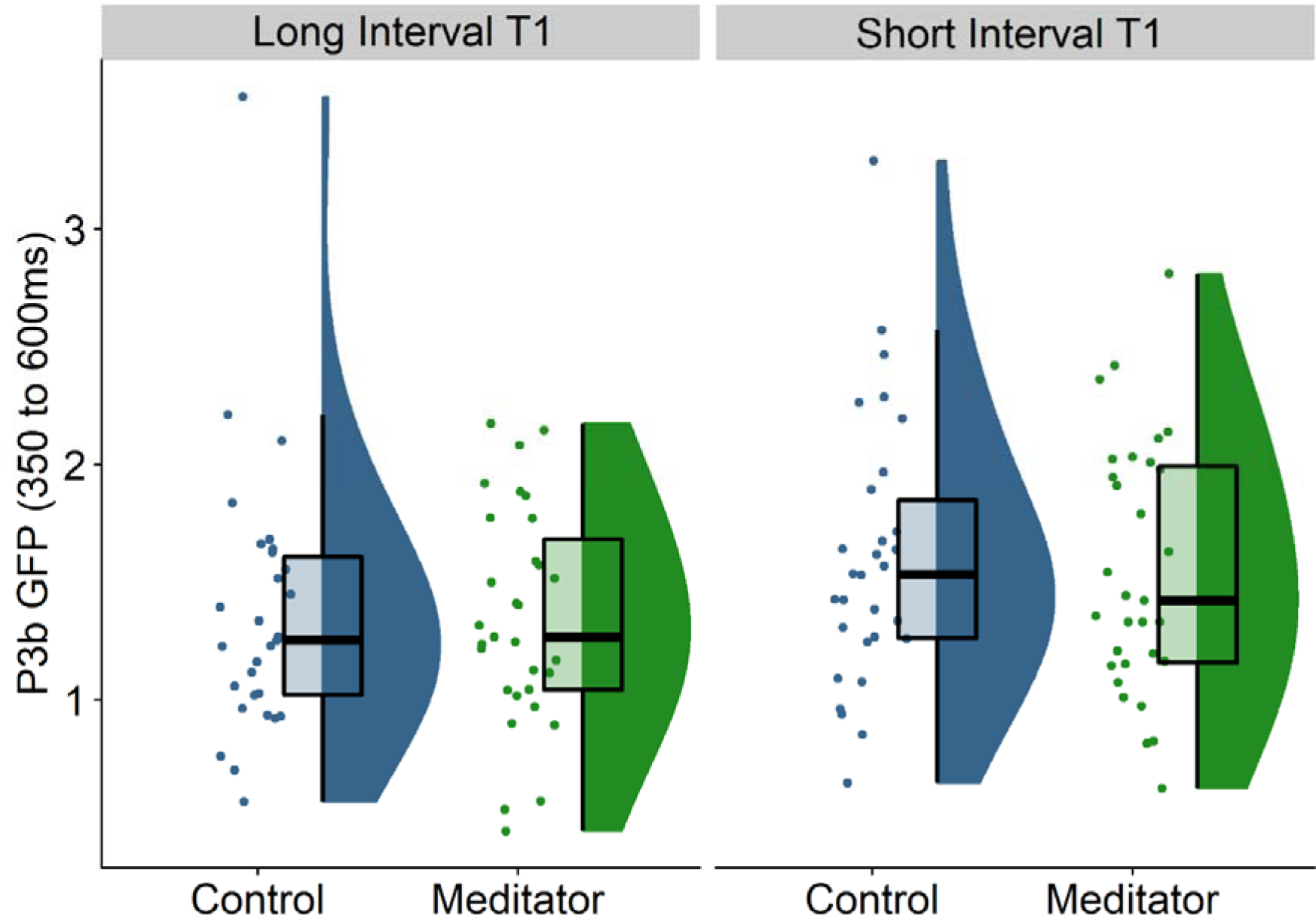
Global field potential (GFP) values averaged across the P3b period of interest (from 350 to 600ms after T1 presentation).

As mentioned in our hypotheses, while Slagter et al (2007) focused on the P3b in response to T1 only, our view is that it is sensible to hypothesise that that effects might occur in components other than the P3b, that ERP amplitudes time locked to T2 might be increased in meditators (EH1), or for the relationship between ERP amplitudes time locked to T1 and T2 to be different in meditators (EH2). To test these exploratory hypotheses (EH1 and EH2), a GFP test was performed across the entire epoch, including all conditions (both T1 and T2 targets and short / long intervals). This test showed a significant interaction between Group and Target from 214 to 258ms following the stimuli (averaged across this time interval: p = 0.002, η_p_^2^ = 0.0914, see Figure 2), which survived multiple comparison controls for duration (global duration control = 41ms). This effect falls within the typical posterior-N2 time window. Within this interaction, controls showed significantly higher GFP amplitudes in response to T1 compared to T2 (p = 0.022, η_p_^2^= 0.1657), while meditators showed no difference between T1 and T2 (p = 0.279, η_p_^2^= 0.0403). When group comparisons were restricted to short interval T1 stimuli only (averaged within the 214 to 258ms window), meditators showed significantly lower posterior-N2 GFP amplitudes than controls (p = 0.029, η_p_^2^= 0.0784, see Figures 2 and 3). To determine the strength of evidence for this significant interaction between Group and Target, averaged GFP values for each participant across both short and long intervals were calculated for both T1 and T2 targets separately and submitted to a repeated measures Bayesian ANOVA design. When comparing the interaction effect against models that did not include the interaction effect, the Bayes Factor showed moderate evidence for the effect (BFincl = 3.411). As such, while hypothesis EH1 was not supported (as meditators did not show larger amplitude ERPs following T2 stimuli), hypothesis EH2 was supported, as meditators showed a more equal distribution of ERP amplitudes between T1 and T2 than controls (although not within the P3b window). Finally, in our test of the exploratory hypothesis that the distribution of ERPs would differ between meditators and controls (EH3), the TANOVA showed no significant main effect of Group or interaction involving Group that exceeded multiple comparison controls for the number of comparisons across the epoch (all p > 0.05).

**Figure 2.**
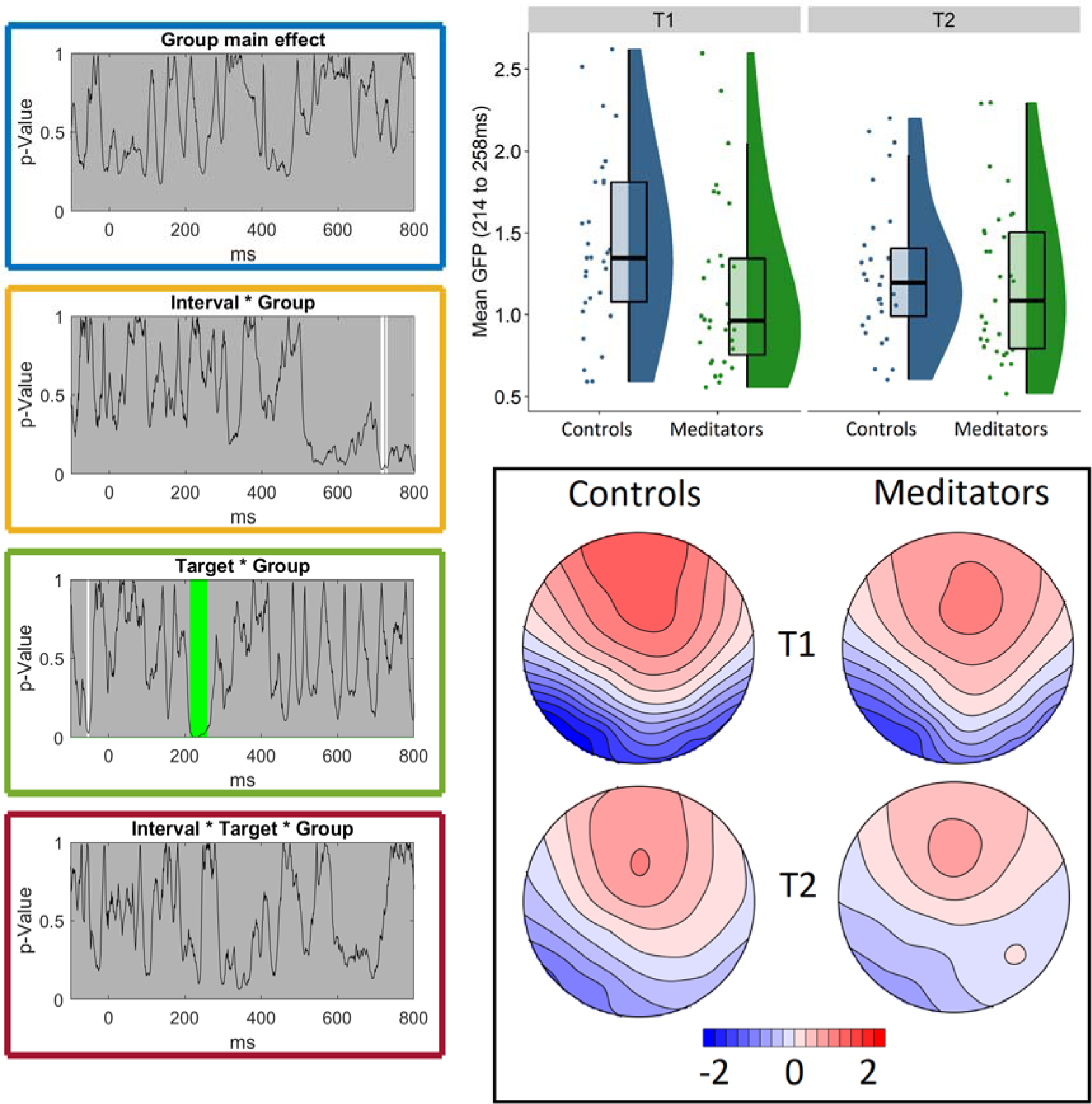
Left: p-value graphs for the main effect of group and interactions involving group for the whole epoch comparisons of the event-related potential (ERP) global field potential (GFP). The black line reflects the p-value, white areas reflect significant time points, and green periods reflect windows where the effect passed global duration controls. Top right: GFP activity in response to the first target (T1) and second target (T2), averaged over the significant window for the test of the interaction between Group and Target (from 214 to 258ms following the stimuli) and averaged across both short and long intervals. Bottom right: the mean (non-normalised) topography within the significant 214 to 258ms period from each group, averaged across T1 and T2 locked epochs separately.

**Figure 3.**
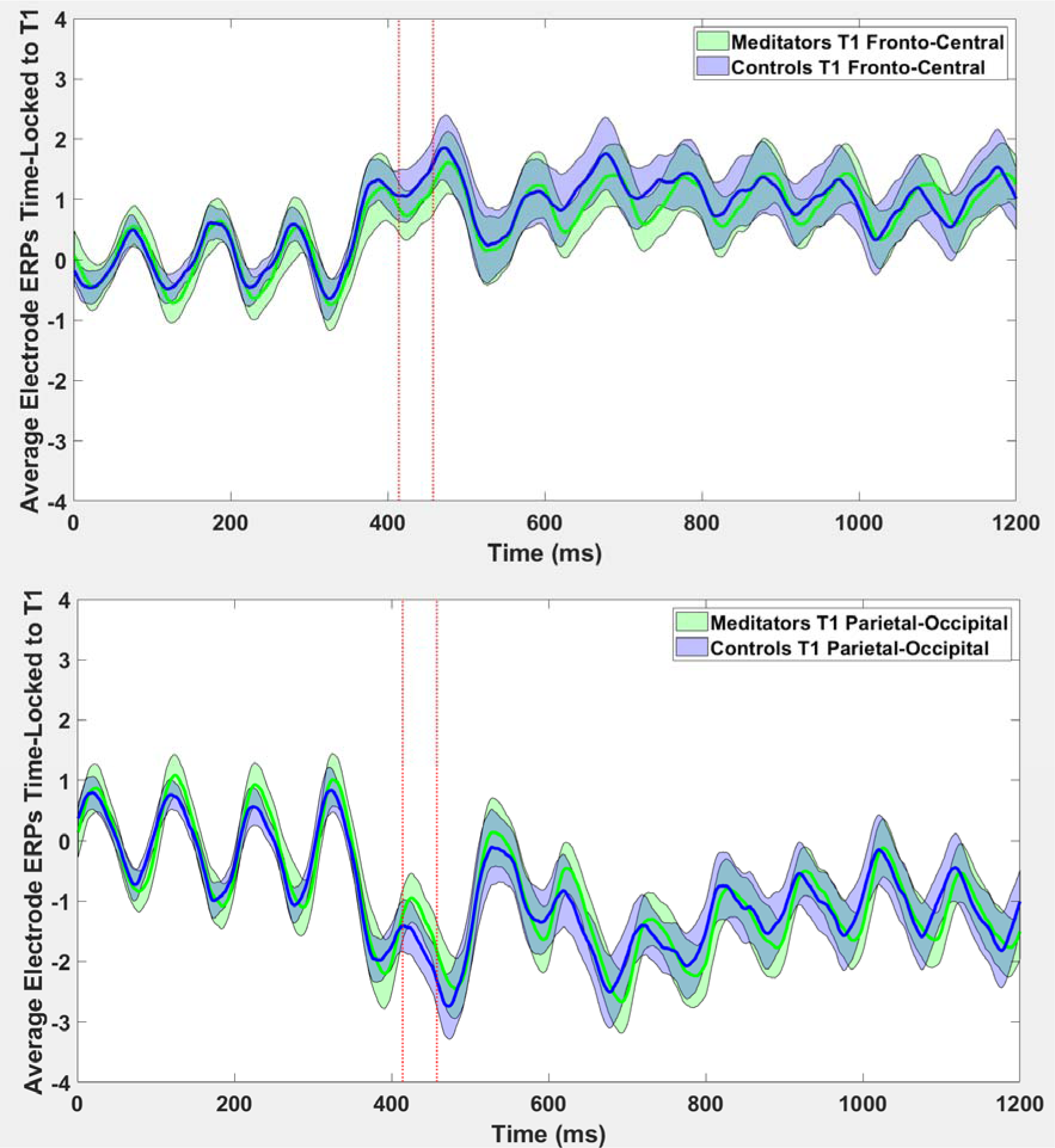
Averaged event-related potentials (ERPs) averaged within fronto-central (top) (F1 Fz F2 FC1 FCz FC2) and parietal-occipital (bottom) electrodes (PO7 PO5 PO6 PO8 O1 Oz O2) time-locked to T1 with the significant period marked (red dashed lines). Note that our analyses were based on the GFP, so while the averaged electrodes demonstrate the difference (with N2 ERPs showing smaller amplitudes in meditators regardless of polarity), our significance tests were not based on these values. Note also the oscillatory pattern in the alpha frequency, synchronised to the stimuli presentation rate.

In the supplementary materials we report exploratory linear mixed models and generalised linear mixed models to explore the potential associations between single trial GFP values within the posterior-N2 effect and whether single trials were responded to correctly to assess potential explanations for this result (section 3b). In brief, the these exploratory analyses showed that correct identification of short interval T2 stimuli was associated with lower posterior-N2 GFP time-locked to T1 (similar to the pattern shown by the meditators) (Figure S3). This suggests that when fewer attentional resources were devoted to processing T1, T2 could be more accurately identified. Additionally, in single trial analysis, the relationship between T2 posterior-N2 GFP, trial number and response accuracy differed between the Groups. To begin with, both meditators and controls were less likely to identify T2 stimuli if their T2 posterior-N2 GFP was high. Controls showed the same pattern throughout the task. However, by the end of the task, this pattern switched for the meditators who were more likely to identify T2 targets when they showed high posterior-N2 GFP values.

### Theta Phase Synchronisation Comparisons

The TCT for TPS showed consistent neural activity across groups and conditions from -280ms across the first 600ms after stimulus presentation, with TCT inconsistency in controls locked to the T1 stimuli prior to this time that did not overlap with any of our significant effects in the RMS test, but did overlap with some of the significant effects w
ithin the TANOVA tests. This demonstrated that our RMS TPS results were not driven simply by inconsistent topographical activation within a single group or condition (supplementary materials section 3c, Figure S5). For our test of hypothesis PH2, that meditators would show higher TPS following short interval T2, RMS TPS was averaged within short interval T2 trials across the 121 to 501ms window for direct comparison with Slagter et al. (2009) (who found an effect within this window, maximal at Fz and FC6). No significant difference was detected (p = 0.086, FDR-p = 0.173, η_p_^2^= 0.0482, BF01 = 1.104). Similarly, for the 309 to 558ms period (where Slagter et al. 2009 found an effect within this window that was maximal at electrode T8), no significant difference was detected (p = 0.118, η_p_^2^= 0.0418, BF01 = 1.373).

However, when all conditions and time points were included in an exploratory analysis of RMS TPS, a significant interaction between Group, Target, and Interval was present from 117 to 295ms (averaged across the significant window: p = 2e-4, η_p_^2^= 0.2358, Figure 4). This effect lasted longer than the duration controls for multiple comparisons over time used by Slagter et al. (2009) (175.1ms). When RMS TPS was averaged within the significant window, Bayesian analysis of the interaction indicated strong support for the alternative hypothesis (BFincl = 41.612), and the model including this Group, Target, and Interval interaction effect as well as the nested comparisons was 5.502e+9 times more likely than the null model (BF10 = 5.502e+9). In assessing the cause of the 3-way interaction with reduced ANOVA designs, our results indicated it was driven by two features: firstly, controls showed larger RMS TPS during long interval T2 trials than short interval T2 trials, while meditators showed very little difference in RMS TPS between the short and long interval conditions (p = 0.0094, η_p_^2^= 0.1718, BFincl = 29.574). Secondly, the interaction was also driven by an effect where meditators showed a more even distribution of RMS TPS between T1 and T2, in comparison to controls who showed higher RMS TPS values to T1 compared to T2 (short interval T1 vs short interval T2) (p = 0.0022, η_p_^2^= 0.1626, BFincl = 25.192). However, counter to the results of Slagter et al. (2009), the interaction was not driven specifically by a difference between groups in short interval T2 TPS (averaged within the 117 to 295ms window showing the significant interaction, there was no significant difference between the groups in short interval T2 RMS TPS, p = 0.136, η_p_^2^= 0.0373). Single electrode analyses replicating Slagter et al.’s (2009) electrode and window of interest showed the same pattern of results as the effect we detected within the 117 to 295ms window, with Bayesian evidence supporting the alternative hypothesis for the interaction between Group and Interval for T2 stimuli (BFincl = 4.621 within Slagter et al.’s (2009) time window, and BFincl = 35.908 when restricted to the significant time period detected in our exploratory analysis, reported in full in the supplementary materials section 3d, Figure S6).

**Figure 4.**
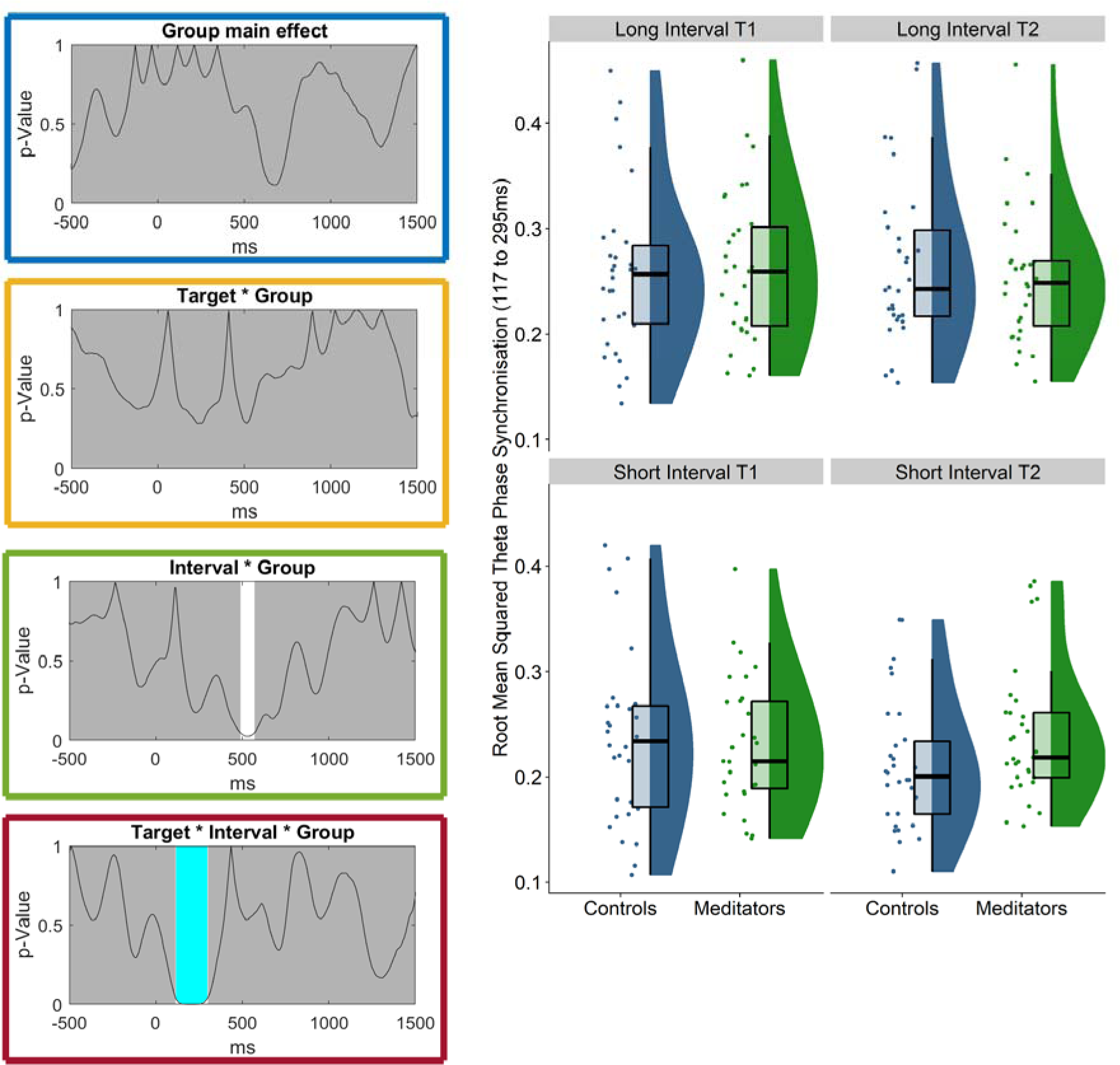
Root mean squared (RMS) comparisons of group, target, and interval for theta phase synchronisation. Left: P-graphs for the main effect of Group and interactions involving Group. The black line reflects the p-value, white areas reflect significant time points and the light blue area indicates the effect that passed Slagter et al.’s (2009) duration control. Right: Theta phase synchronisation RMS showing the significant interaction of interest between Group, Interval and Target from the averaged activity within the 117 to 295ms window (p = 0.004, η_p_^2^= 0.2526, BFincl = 41.612).

To assess whether these differences in TPS might have behavioural relevance, we performed Pearson’s correlations between TPS and percentage correct from short interval T2 trials across both groups together. These results indicated that TPS from all conditions correlated with short interval T2 accuracy (statistics reported in full in Table 3, and scatterplots for these comparisons can be viewed in Figure 5). There was also an interaction between Group and Interval from 455 to 560ms (averaged across the significant window: p = 0.0218, η_p_^2^= 0.1520). However, this period did not survive the 175.1ms minimum duration used by Slagter et al. (2009). No other main effect or interaction involving group was significant for any part of the epoch (all p > 0.10).

**Table 3:** Pearson’s Correlations between percent correct responses to the second target stimuli (T2) in short interval trials and the averaged root mean squared (RMS) theta phase synchronisation (TPS) within the 117 to 295ms period in response to both the first target stimulus (T1) and T2.

| Variable | Short Interval T1<br>RMS TPS<br>Pearson's r (*p-value*) | Long Interval T1<br>RMS TPS<br>Pearson's r (*p-value*) | Short Interval T2<br>RMS TPS<br>Pearson's r (*p-value*) | Long Interval T2<br>RMS TPS<br>Pearson's r (*p-value*) |
| --- | --- | --- | --- | --- |
| Percentage Correct Short Interval T2 | .324* (.012) | .381** (.003) | .269* (.037) | .293* (.023) |
| BF10: | 2.159 | 13.087 | 1.333 | 2.014 |

\*p < .05, \*\*p < .01, \*\*\*p<.001

**Figure 5.**
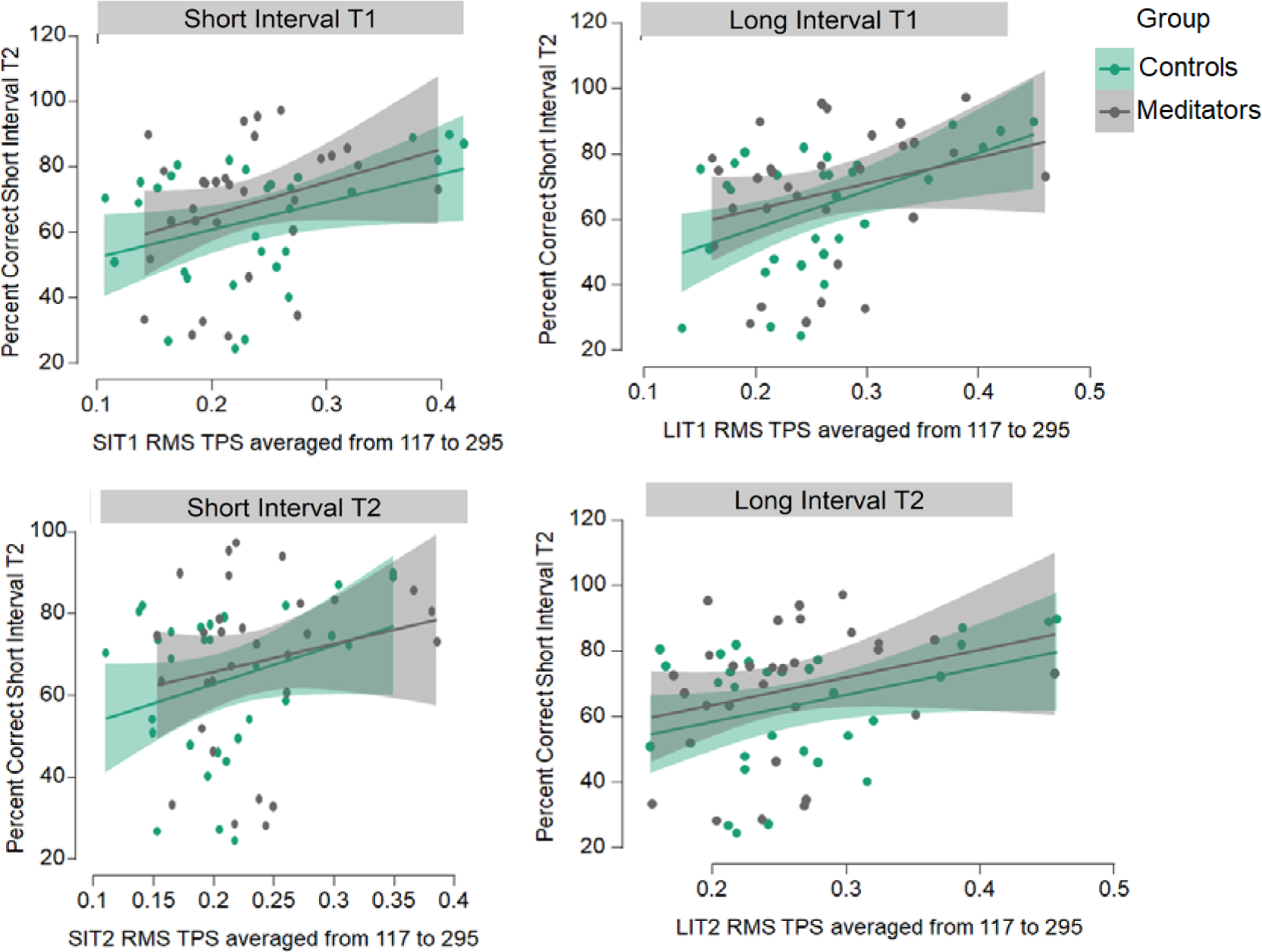
Scatterplots depicting the correlations between root mean squared (RMS) theta phase synchronisation (TPS) averaged within the significant window (117 to 295ms) from each condition and accuracy at detecting the second target stimuli (T2) in short interval trials. Note the common pattern across all groups and conditions. The grey and light green areas reflect relative variance from the line of best fit at point on the x-axis.

With regards to our exploratory hypothesis that the scalp distribution of TPS would differ between groups (EH4), the TANOVA, including all conditions and all time points, showed an interaction between Group and Target during the presentation of the distractor stimuli from -385 to -100ms prior to T1, which lasted longer that Slagter et al.’s (2009) duration control for multiple comparisons (175.1ms). When averaged across the significant window the statistics were: p = 0.001, η_p_^2^= 0.0609 (see Figure 6). When the interaction was explored by averaging TPS within the significant period and performing TANOVA comparisons between the groups for T1 and T2 stimuli separately, the effect was shown to be driven by a difference in TPS distribution between the groups prior to T1 stimuli (p = 0.018, η_p_^2^= 0.0336), with meditators showing more TPS in occipital electrodes (meditator minus control t-max at Oz = 2.908) and meditators showing less TPS in right frontal electrodes (meditator minus control t-min at F6 = -3.384). Groups did not differ in TPS locked to T2 stimuli (p = 0.1358). It is worth noting that the period that showed the significant result overlapped with a period of topographical inconsistency in T1-locked TPS in the control group (with inconsistent topographical distributions across the control group prior to - 280ms). This suggests that at least part of the interaction may have been driven by an inconsistent topographical pattern in the control group (rather than a between-group difference during that time period). No other differences were present in any of the main effect or interactions involving group within any time point in the epoch (all p > 0.05).

**Figure 6.**
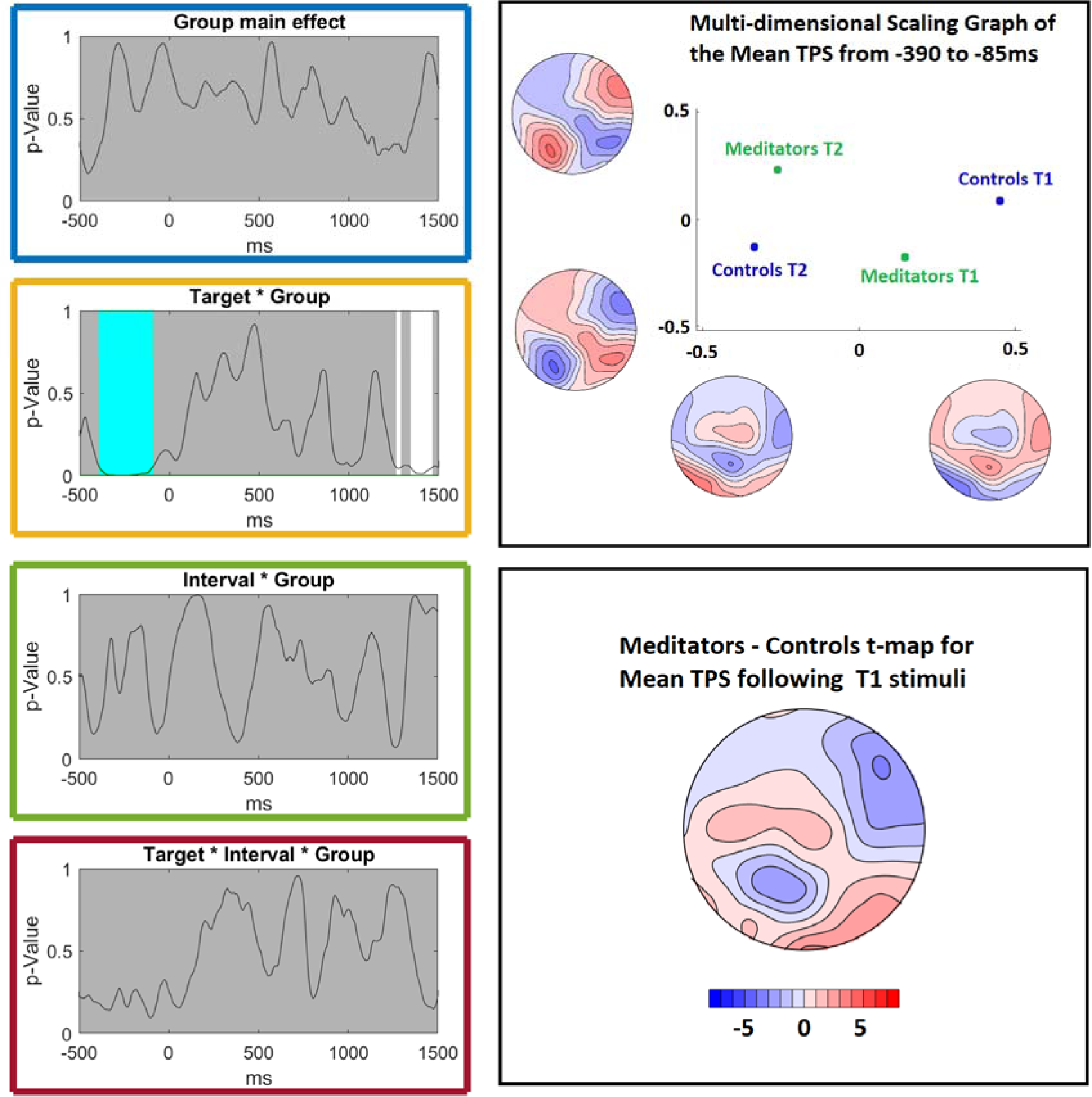
Topographical Analysis of Variance (TANOVA) test results for the theta phase synchronisation (TPS). Left: p-graphs for the main effect of Group and each interaction involving Group. The black line reflects the p-value, the white areas reflect significant time points, and the light blue periods reflect windows where the effect passed Slagter et al.’s (2009) duration controls. Right top: A multi-dimensional scaling graph depicting the differences between each group’s TPS topographies in response to the first (T1) and second (T2) target stimuli averaged during the window of the significant Group x Target interaction (-390 to -85ms around the target). Within the multi-dimensional scaling graph, the topography maps indicate the ends of the eigenvector spectrum in each of the x- and y-axis, and the points on the graph indicate where each group and condition’s mean topography lay on that spectrum (for both the x and y axis) relative to the other points in the graph (note that the topographies along the x and y axis do not represent the actual topography for a group / condition). As such, the interaction between Group and Target in topographical activation is demonstrated by the graph. Right bottom: the t-map for the meditator minus control theta phase synchronisation topography for T1 stimuli (averaged from -390 to -85ms around T1), after normalisation for overall amplitude (so that all individuals had a GFP = 1). Red indicates areas where meditators showed higher values, blue indicates areas where controls showed higher values (indicating that topographical differences were present, without suggesting that TPS was higher in the control group in a specific electrode, due to the normalisation for amplitude).

### Alpha-power Comparisons

The TCT for RMS alpha-power showed consistent neural activity across all groups and conditions from -400ms until the end of the epoch, indicating our alpha-power results were not driven simply by inconsistent topographical activation within a single group or condition (details are reported in the supplementary materials section 3e, Figure S7). When RMS alpha-power was averaged across the -31 to 160ms window for direct comparison with Slagter et al. (2009) and test of our third primary hypothesis (PH3 – that meditators would show greater alpha-power around T1 presentation), no significant difference was detected (p = 0.2976, FDR-p = 0.3968, η_p_^2^= 0.0189, BF01 = 2.379). The exploratory RMS test for alpha-power, including all time points within the epoch time-locked to T1 stimuli, showed a significant main effect of Group from 475 to 685ms, in which meditators showed less alpha-power (averaged within this window: p = 0.023, η_p_^2^ = 0.0844, see Figure 7). This effect passed the duration controls implemented by Slagter et al. (2009) (83.5ms for alpha). No interaction was detected between Interval and Group in RMS alpha-power that lasted longer than Slagter et al.’s (2009) duration controls. Nor was there any Group main effect or interaction between Group and Interval in the alpha-power TANOVA (all p > 0.05), providing a null result for hypothesis EH5 (that there would be differences between the groups in the scalp distribution of alpha-power).

**Figure 7.**
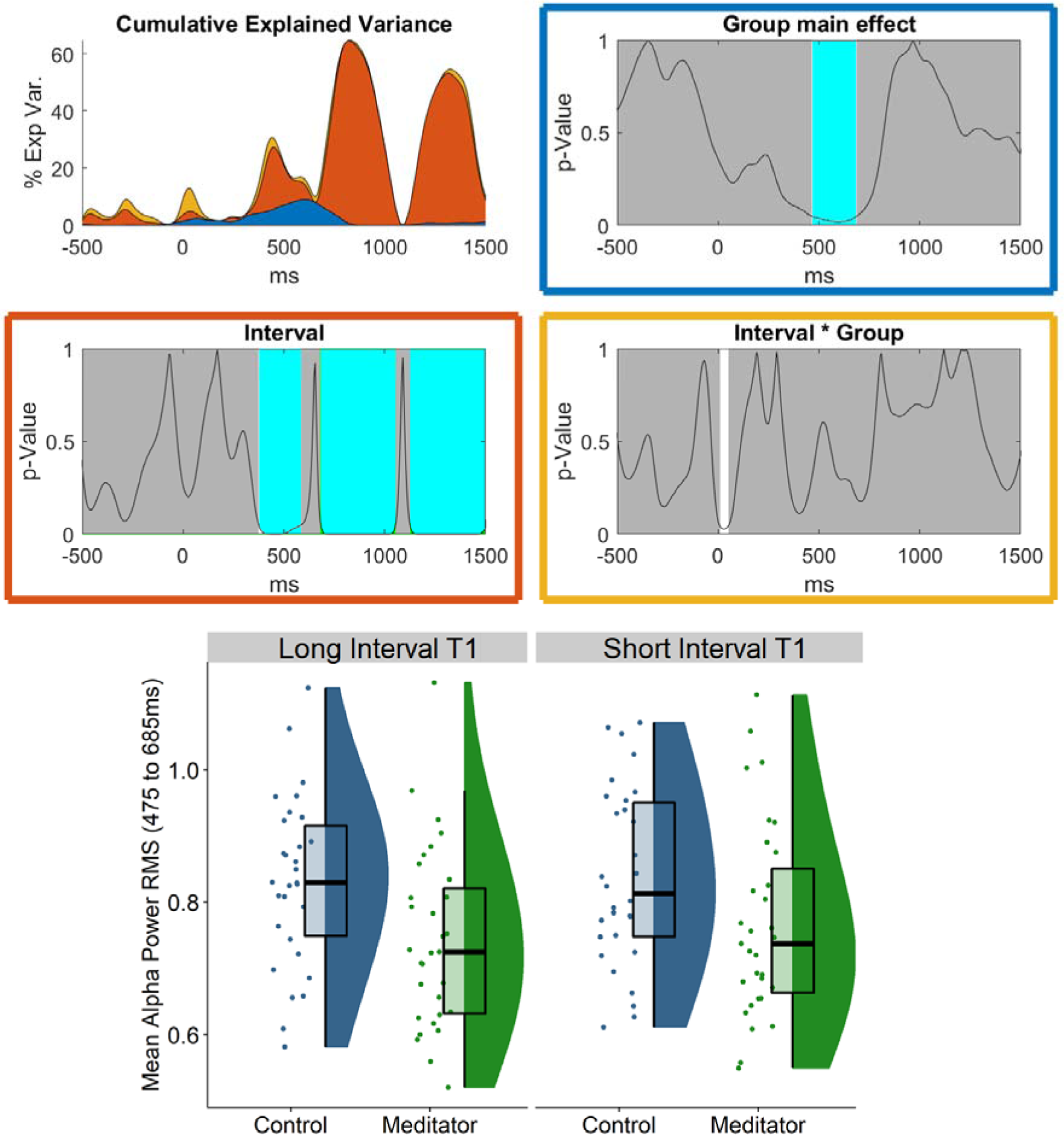
Root mean squared (RMS) alpha-power comparisons time-locked to T1 stimuli onset. Top left: The cumulative variance explained (η_p_^2^) at each time point across the epoch by each main effect and condition, with each colour reflecting the η_p_^2^ from the effect being tested, colour coded to match the p-graphs. Top right and middle: the p-graphs for the main effect of interval (orange, middle left), group (blue) and interaction between interval and group (yellow, middle right). The black line reflects the p-value, white areas reflect significant time points, and light blue periods reflect windows where the effect passed Slagter et al.’s (2009) duration controls. Bottom: Mean RMS alpha-power averaged within the 475 to 685ms window following the first target stimuli (T1) for each participant (baseline corrected to alpha-power across the entire epoch).

To explore potential explanations for these results, we performed a number of additional tests of the pattern of relationships between trial number, single trial accuracy (to assess potential learning across the task), and alpha-power within this significant period (these are reported in the supplementary materials section 3e). In brief, the baseline corrected RMS alpha-power within the 475 to 685ms window decreased across trials as participants completed the task, which was concurrent with improved performance across the task, suggesting participants may have been learning attention-based strategies to enable improved short interval T2 detection. However, across all participants, averaged baseline corrected alpha-power RMS within the 475 to 685ms window after T1 did not correlate with the accuracy of short interval T2 detection. Further, an exploratory linear mixed model indicated that *incorrect* responses were associated with slightly, but significantly, lower short interval RMS alpha-power than correct responses (supplementary materials section 3e, Figure S11). However, lower short interval trial RMS alpha-power within a later 685 to 1050ms window was strongly associated with *correct* responses (Figure S12). Short interval alpha-power RMS was also strongly correlated between these two periods. This relationship was stronger within incorrect trials than for correct trials, and long interval RMS alpha-power increased in the later 685 to 1050ms window compared to the earlier period in both groups. This suggests that lower short interval RMS alpha-power in the later 685 to 1050ms window was required to identify the T2 stimuli. As such, perhaps the lower RMS alpha-power in the earlier period might have been a compensatory mechanism on trials when participants noticed their attention waning, reflecting an attempt to regulate alpha-power in the later period, during which low alpha-power was vital for stimulus processing.

### Alpha Phase Synchronisation Comparisons

In test of our fourth primary hypothesis (PH4 -that APS would be reduced in the meditation group during the presentation of the distractor stimuli prior to T1 stimuli), we conducted an RMS test of APS time-locked to T1 stimuli averaged across the period where distractor stimuli were presented prior to T1 (within the -414 to -214ms window for direct comparison with Slagter et al. (2009), our results indicated a non-significant main effect of Group, where meditators showed higher APS, which is in the opposite direction to the findings provided by Slagter et al. (2009) (p = 0.061, FDR-p = 0.173, η_p_^2^= 0.0586). Additionally, our exploratory analysis of APS across the entire epoch showed a significant main effect of Group from -258 to -90ms, and from 288 to 1500ms (both of which survived Slagter et al.’s (2009) duration controls of 83.5ms, see Figure 8). Within both the shorter pre-stimulus and longer post-stimulus period, meditators showed larger RMS APS (averaged within the -258 to -90ms period = 0.031, η_p_^2^= 0.072, BFincl = 2.089, averaged within the 288 to 1500ms period: p = 0.018, η_p_^2^= 0.092, BFincl = 1.752, with the best model including the main effect of group and the main effect of interval, BF10 = 42.23 for the average interval from 288 to 1500ms). Our results also indicated a brief significant interaction between Group and Interval in APS RMS (706 to 786ms) which did not pass Slagter et al.’s (2009) duration controls.

**Figure 8.**
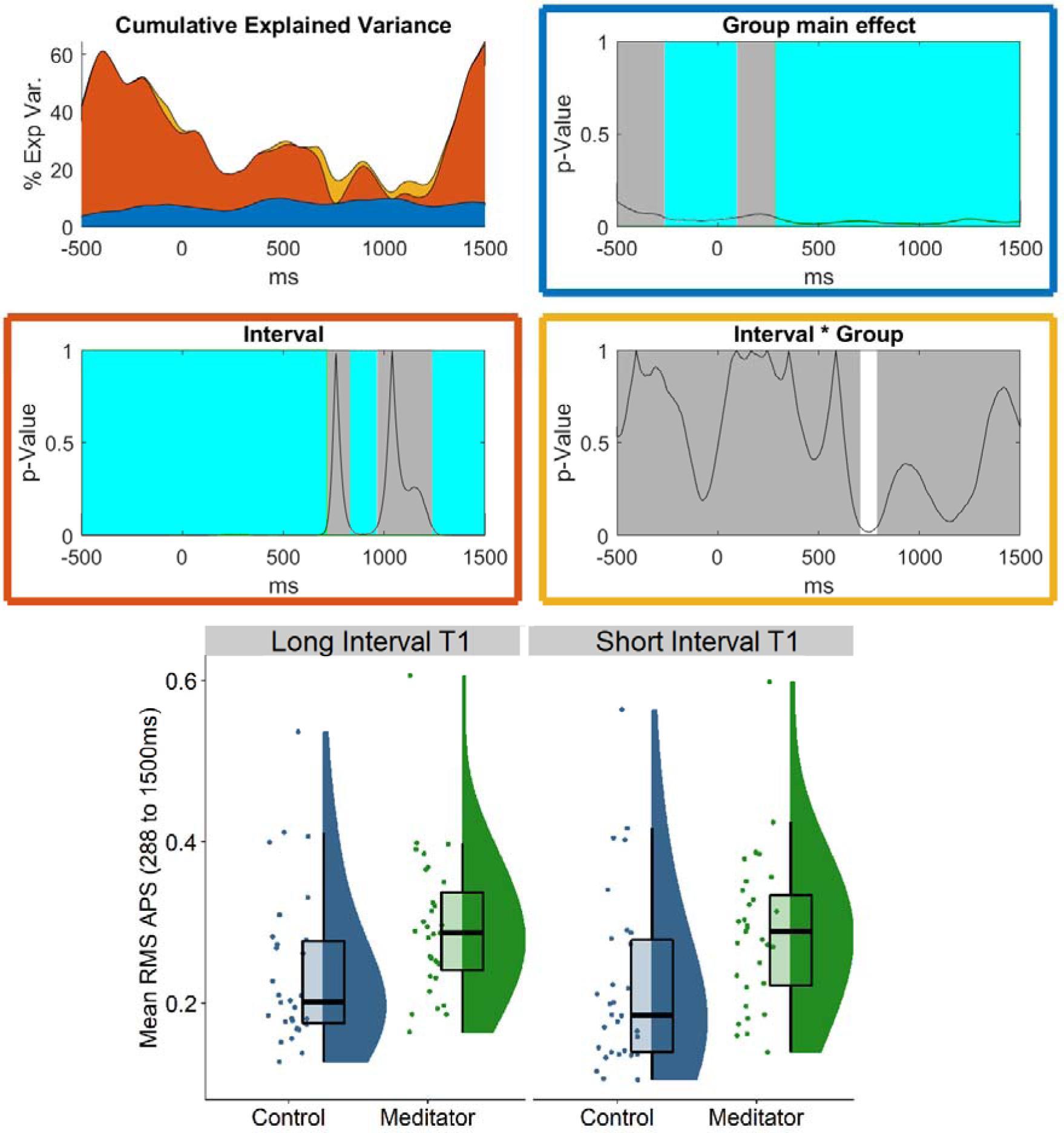
Root mean squared (RMS) alpha phase synchronisation (APS) comparisons time-locked to T1 stimuli onset. Top left: The cumulative variance explained (η_p_^2^) at each time point across the epoch by each main effect and condition, with each colour reflecting the η_p_^2^from the effect being tested, colour coded to match the p-graphs. Top right and middle: the p-graphs for the main effect of interval (orange, middle left), group (blue) and interaction between interval and group (yellow, middle right). The black line reflects the p-value, white areas reflect significant time points, and light blue periods reflect windows where the effect passed Slagter et al.’s (2009) duration controls. Bottom: Mean root mean squared alpha phase synchronisation (RMS APS) from each group in response to T1 long (LIT1) and short (SIT1) interval trials, averaged within the significant window from the RMS APS test.

With regards to the TANOVA test of APS (which tested exploratory hypothesis EH6 – that meditators would show a different scalp distribution of APS), a significant Group main effect was detected from 990 to 1500ms where meditators showed higher APS values in fronto-central and parieto-occipital electrodes and lower APS values in lateral central electrodes (p = 0.007, η_p_^2^= 0.0408, with a meditator minus control t-max of 3.417 at PO5 and t-min of -3.035 at C5, see Figure 9). This effect passed Slagter et al.’s (2009) duration control (83.5ms). There was also a Group main effect in the TANOVA from -244 to -2ms (p = 0.030, η_p_^2^ = 0.0329, and brief significant interaction between Group and Interval in the APS TANOVA (150 to 280ms, p = 0.011, η_p_^2^ = 0.0364), both of which passed Slagter et al.’s (2009) duration control (83.5ms).

**Figure 9.**
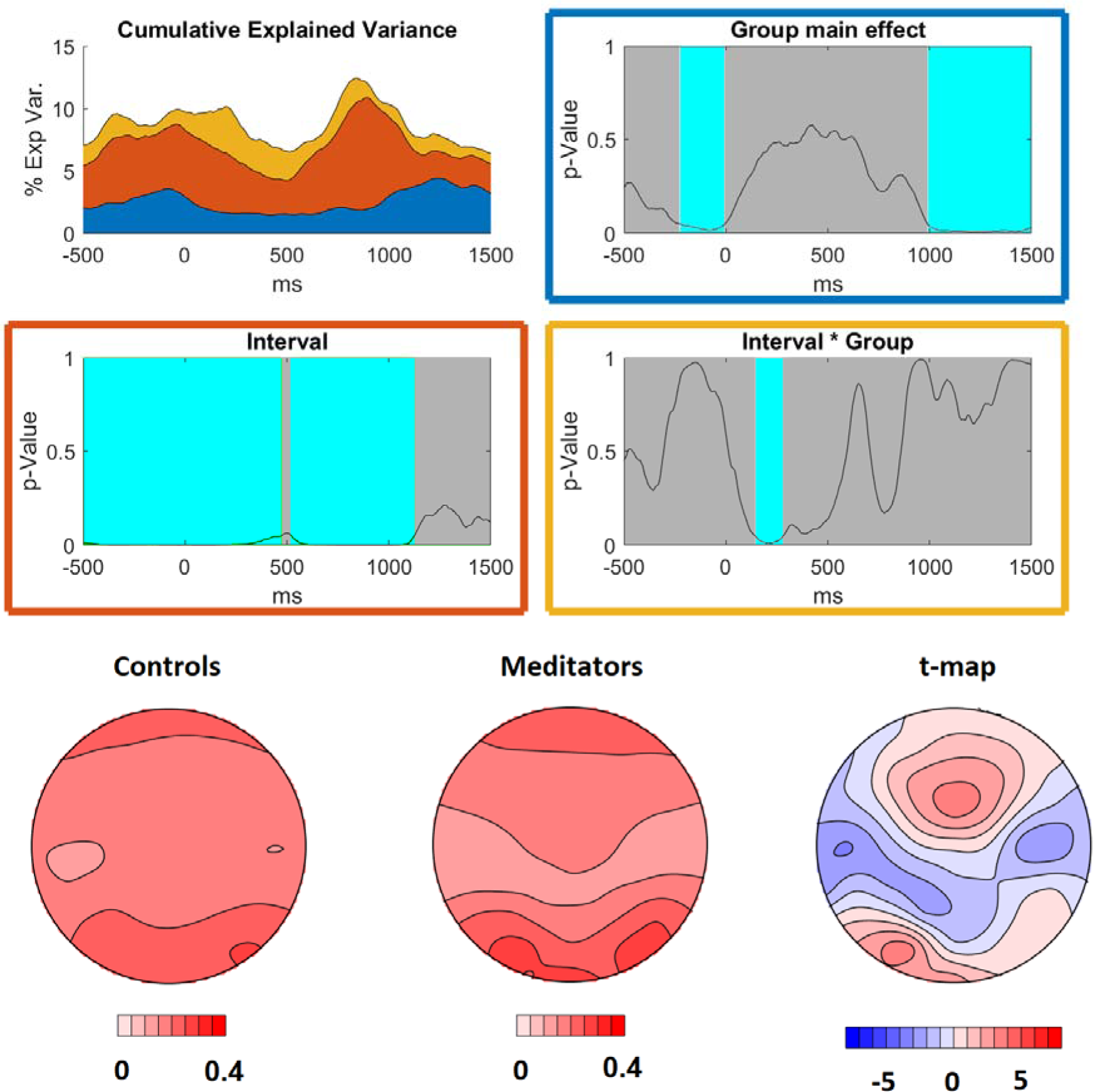
Alpha phase synchronisation (APS) topographical analysis of variance (TANOVA) comparisons time-locked to the onset of the first target stimuli (T1). Top left: The cumulative variance explained (η_p_^2^) at each time point across the epoch by each main effect and condition, with each colour reflecting the η_p_^2^ from the effect being tested, colour coded to match the p-graphs. Top right and middle: the p-graphs for the main effect of interval (orange, middle left), group (blue) and interaction between interval and group (yellow, middle right). The black line reflects the p-value, white areas reflect significant time points, and light blue periods reflect windows where the effect passed Slagter et al.’s (2009) duration controls. Bottom: Topography maps for APS averaged within the 990 to 1500ms period for each group and the t-map of meditator APS minus control APS after normalisation for overall amplitude (so that all individuals had a GFP = 1). Red indicates areas where meditators showed higher values, blue indicates areas where controls showed higher values (indicating that topographical differences were present, without suggesting that APS was higher in the control group in a specific electrode, due to the normalisation for amplitude).

RMS APS averaged within the 282 to 1500ms period significantly correlated to percentage correct for short interval T2 trials, in both short interval and long interval trials - for the correlation between APS RMS during short interval T1 trials and T2 short interval percentage correct: Pearson’s r = 0.314, p = 0.014, BF10 = 3.093, and for the correlation between APS RMS during long interval T1 trials and T2 short interval percentage correct: Pearson’s r = 0.307, p = 0.016, BF10 = 2.717. Scatterplots depicting these correlations can be viewed in the supplementary materials (Figure S14). These correlations may indicate that participants who synchronised their alpha oscillations more consistently with the stimulus stream (which was presented at 10Hz, within the alpha frequency) were better able to perceive and correctly identify the T2 stimuli. It is worth noting that the T2 stimuli in short interval trials were presented at 300ms, just after the point at which the meditation group showed higher alpha synchronisation to the stimuli.

### Behavioural and EEG Epoch Inclusion Comparisons

Levene’s test indicated the assumption of equality of variances was met for all conditions within the analysis of the behavioural data (all p > 0.15). However, the Shapiro-Wilk test indicated significant deviations from normality for 8/10 of the Condition x Group combinations, so robust statistics were implemented in R, using the mixed ANOVA (bwtrim function) from the WRS2 package (Field and Wilcox, 2017). Violations of the assumptions of traditional parametric ANOVAs (including normality violations) do not affect these robust statistics. However, only Group x Condition designs are currently available (rather than group x condition x condition), so this analysis was restricted to a Group x Interval comparison for T2 responses only (as the primary comparison of interest), and the originally planned parametric statistical analyses are reported in the supplementary materials (section 3a). Means, standard deviations, and both parametric and robust statistics are presented in Table 4, and the data can be viewed in Figure 10.

**Table 4.** Attentional Blink Behavioural Performance Means (M), Standard Deviations (SD) and Statistics for Each Group and Condition. T1 = the first target stimuli. T2 = the second target stimuli.

|  | Meditators<br>*M (SD)* | Controls<br>*M (SD)* | Statistical test result | p-value | Effect Size | Bayesian Factor |
| --- | --- | --- | --- | --- | --- | --- |
| Long interval T1<br>percentage correct | 92.593<br>(4.961) | 91.410<br>(6.290) | Group main effect: $F(159) = 0.698$<br>Robust statistics, Group main effect for T2 only:<br>value (133.997) = 0.3250 | $p = 0.407$<br>$p = 0.5724$ | $\eta^2G = 0.006$ | $BF_{excl} = 4.140$ |
| Short interval T1<br>percentage correct | 92.321<br>(4.577) | 90.800<br>(4.981) | Target main effect: $F(1,59) = 160.152$ | $p < 0.001^{**}$ | $\eta^2G = 0.407$ | $BF_{incl} = 9.008e+12$ |
| Long interval T2<br>percentage correct | 77.509<br>(12.820) | 75.409<br>(15.332) | Interval main effect: $F(159) = 26.682$ | $p < 0.001^{**}$ | $\eta^2G = 0.049$ | $BF_{incl} = 5.920e+13$ |
| Short interval T2<br>percentage correct | 67.059<br>(20.848) | 63.913<br>(18.929) | Group x Target: $F(159) = 0.149$ | $p = 0.700$ | $\eta^2G = 6.411e-4$ | $BF_{excl} = 4.197$ |
| | | | Group x Interval: $F(159) = 0.098$ | $p = 0.755$ | $\eta^2G = 1.898e-4$ | $BF_{excl} = 3.957$ |
| | | | Target x Interval: $F(159) = 25.415$ | $p < 0.001^{**}$ | $\eta^2G = 0.042$ | $BF_{incl} = 4.523e+17$ |
| | | | Group x Target x Interval: $F(159) = 0.029$ | $p = 0.866$ | $\eta^2G = 4.973e-5$ | $BF_{excl} = 4.019$ |
| | | | Robust Statistics: Group x Interval for T2 only:<br>value (133.898) = 0.212 | $p = 0.642$ | | |

$^{**}p < 0.001$ .

**Figure 10.**
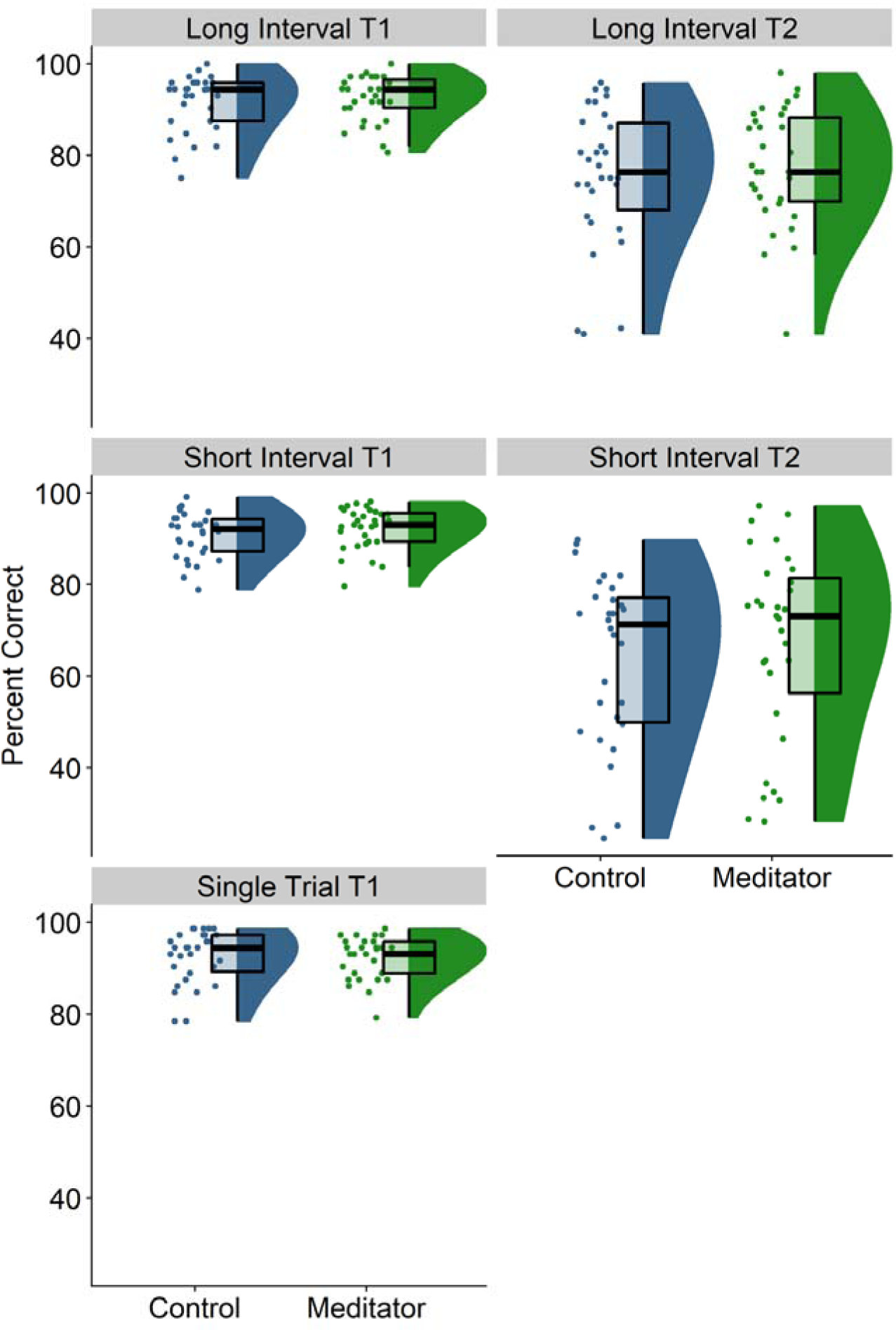
Attentional Blink performance, measured in percentage correct for each group and condition. Long interval refers to conditions in which the T2 stimulus was presented 700ms after T1. Short interval refers to conditions in which the T2 stimulus was presented 300ms after T1. Figures on the left (T1) indicate the percentage of T1 stimuli correctly identified by each participant, whilst figures on the right (T2) indicate the percentage of T2 stimuli correctly identified by each participant. The single trial T1 label refers to T1-only trials (where no T2 stimulus was presented).

In testing our first replication hypothesis (RH1 - that our meditation group would show a reduced AB effect, with more correct responses to short interval T2 stimuli), the robust statistics showed no main effect of Group for percent correct in response to T2: value (1,33.997) = 0.325, p = 0.572, and no interaction between Group and Interval: value (1,33.898) = 0.220, p = 0.642. The parametric statistics showed the same pattern of null results. The Bayesian statistical model including Group or interactions that involved Group as a factor was 259.326 less likely than the model that only included Target, Interval, and the interaction between Target and Interval (BF01 = 259.326). These results suggest it is highly unlikely that the meditation group showed higher percentage correct in any condition compared to the control group.

No main effects or interactions involving Group were found for the number of epochs provided by each participant for each condition (all p > 0.1). The TCT for the ERP data also showed mostly consistent neural activity across groups and conditions, with a brief period of inconsistency that did not overlap with any of our significant effects. These two tests indicate our ERP results were not driven simply by differences in the number of epochs included in ERP averages or inconsistent topographical activation within a single group or condition (details of these tests are reported in the supplementary materials, section 3a and 3b, Table S1 and Figure S1).

## Discussion

This study aimed to comprehensively examine if neurophysiological markers of attention differed between community meditators and non-meditator controls. In our sample of meditators with typical daily MM practice, our results did not show support for our primary hypotheses regarding the neurophysiological markers obtained from within our time windows of interest (the P3b, TPS, alpha-power, and APS, with the windows of interest overlapping with the significant effects reported by Slagter et al. (2007, 2009)). No differences were found between meditators and non-meditators in the amplitude or distribution of the P3b neural following T1 or T2 stimuli in the attention blink task. Nor were there any differences between meditators and non-meditators in TPS, alpha-power, or APS within our a priori selected time windows of interest. Frequentist statistics provided null results, and Bayesian statistics provided weak to moderate evidence against these primary hypotheses.

However, our exploratory analyses (which included all time points within the epochs around all T1/T2 and short/long interval conditions) did show significant effects, which were further supported by very strong Bayesian evidence in favour of the alternative hypothesis. In particular, meditators showed more equal posterior-N2 amplitudes across T1 and T2 stimuli than non-meditators (who showed larger posterior-N2 amplitudes to T1 than T2). Similarly, meditators showed more equal TPS values between the first and second target in short interval trials, and meditators showed similar TPS values to T2 in both short and long interval trials, in comparison to controls who showed higher TPS following the first target, and higher TPS to T2 in long interval compared to short interval trials. Meditators also showed lower alpha-power than controls during a period where short interval T2 stimuli would be processed, and increased APS to T1 stimuli. These effects are aligned with theoretical perspectives on the effects of mindfulness, and align with Slagter et al.’s (2007, 2009) explanation of their results that meditators distribute their neural activity more equally across stimuli, rather than biasing responses towards T1 (however, our results did not align with Slagter et al.’s time windows of significant results). Each pattern of neural activity shown by the meditation group was also associated with higher performance, either correlated with percentage correct across all participants, or associated with correct responses rather than incorrect responses in single trial analyses, suggesting the activity shown by meditators might reflect functionally relevant attentional mechanisms. However, unexpectedly, our analyses of behavioural performance provided non-significant frequentist results, and our results showed strong overall Bayesian evidence against any main effect or interaction that involved group. We discuss the details and implications of these findings in the following.

### The posterior-N2 and P3b

Our primary analysis did not detect a difference in the P3b following T1 stimuli in our sample of community meditators. However, our exploratory analyses showed that the meditator group generated an equal amplitude posterior-N2 response across T1 and T2 stimuli, while controls showed higher posterior-N2 responses to T1 stimuli than T2 stimuli. As such, while our study did not replicate Slagter et al.’s (2007) findings with regards to the P3b, our result is conceptually similar, suggesting that meditators distributed attentional resources more equally distributed across the two stimuli. Previous research in healthy control individuals has also demonstrated a reduced posterior-N2 to T2 stimuli following short interval trials, suggested to reflect a lack of attentional engagement to enable stimuli processing (Zivony et al., 2018). As such, our results suggest that meditators are more equally distributing the engagement of attentional resources across the two AB stimuli. In support of this, an exploratory single trial analysis of the posterior-N2 GFP showed that correct identification of short interval T2 stimuli was associated with a smaller posterior-N2 GFP time-locked to T1, suggesting that when fewer attention resources were devoted to processing T1, T2 could be more accurately identified. As such, although the meditation group did not show higher task performance overall, their neural activity averaged within each condition showed the same pattern that was associated with higher performance.

It is not clear, however, why our study detected differences in the posterior-N2 rather than the P3b. This inconsistency might be explained by a progressive change of neural activity during the AB task with more intensive meditation experience. Slagter et al.’s (2007) sample underwent a 3-month intensive retreat, while our participants were experienced meditating members of the lay public (although with an average of six years of meditation experience, and an average of approximately 7 hours per week of practice at the time of the study). However, if this is the case, it is not clear why the less experienced meditators in our study would show altered T1 processing at a shorter delay following T1 presentation than Slagter et al.’s (2007) sample of more experienced meditators. It could be that age is a driving factor, as Slagter’s participant’s median age was 41, whereas the median age of our meditation group was 35, and ERP latency is known to increase with age (Polich, 1997). Another possible explanation is that the EEG pre-processing and analysis techniques used in the current study were updated to fit more recent perspectives on best practice for pre-processing and analysing EEG data compared to those used by Slagter et al. (2007, 2009). Perhaps most importantly, the current study used a high pass filter of 0.25Hz, whereas Slagter et al. (2007) used a high pass filter of 1Hz. The amplitude of ERPs, and particularly the P3b, have been shown to be considerably affected by high pass filtering out < 1Hz data, as the P3b is produced at least in part by < 1Hz activity (Rousselet, 2012; Tanner et al., 2016). As such, the P3b data Slagter et al. (2007) analysed may have had considerable signal removed from the P3b, and their analysis may have been adversely affected.

### Theta Phase Synchronisation

Our primary analysis of TPS to short interval T2 trials showed no difference between meditators and controls. In contrast, our exploratory test of TPS showed strong Bayesian evidence that while controls showed higher TPS to T2 for long interval trials than short interval trials, meditators showed similar TPS to T2 for both short and long interval trials. Strong Bayesian evidence also indicated that meditators showed a more even distribution of TPS between the first and second target in short interval trials, in comparison to controls who showed higher TPS following the first target. Multiple validation checks of this test demonstrated the same result (including single electrode analyses averaged within our a priori time window of interest, and a repeat of the test that excluded participants who provided fewer epochs, ensuring the test possessed maximal validity). These results align with Slagter et al.’s (2009) interpretation that theta synchronisation reflects increased consistency of neural processes, allowing increased attention as a result of meditation training. Our results also support this interpretation, indicating that theta synchronisation was higher following T2 in long interval trials than short interval trials (suggesting theta synchronisation to T2 is disrupted by T1 processing in short interval trials) and that higher theta synchronisation was related to performance.

However, despite the association between increased theta synchronisation and performance and the higher TPS in our meditation group, we found evidence against increased AB task accuracy in our meditation group. Our significant result also only overlapped with the first half of the window in which Slagter et al. (2009) detected increased TPS in their meditators after the retreat, and unlike Slagter et al. (2009), our TPS result was not present when the analysis was focused specifically on the difference between meditators and controls in TPS following short interval T2 trials. This may suggest that while typical community meditation is associated with an effect on theta synchronisation attentional mechanisms, the theta synchronisation after stimulus presentation is not as prolonged as in post-intensive-retreat meditators, and the effect is weaker, only appearing relative to the non-short interval T2 conditions (in which theta synchronisation is perhaps less vital for task performance than it is in the commonly attentional blinked short interval T2 condition). However, the more equal distribution of TPS to short interval T2 stimuli in meditators in our study does still suggest that the meditation group is distributing limited attentional resources to better encode the T2 stimuli. The efficacy of this neural strategy seems to be reflected in the correlation between higher TPS and higher accuracy at accurately identifying short interval T2 stimuli. Our exploratory analysis of the distribution of TPS also indicated that meditators showed more TPS in occipital electrodes prior to T1 stimuli than controls. There was also a more consistent topographical distribution of activity within the meditation group than within the control group, perhaps indicating a consistent synchronisation of oscillations to the target stream in a functionally relevant brain region in preparation for the detection of the relevant stimuli. However, if this interpretation is correct, it is not clear why the meditation group did not show higher accuracy than the control group. As such, our exploratory results require replication, and it may be that ultimately research will show there is no significant difference in TPS between meditators and non-meditators.

### Alpha-power

The current study did not find a significant difference in our primary analyses focused on specific alpha-power and alpha phase synchronisation time windows (with time windows of interest derived from Slagter et al. 2009). However, in our exploratory analysis, the meditation group showed a larger reduction in the level of ongoing alpha-power from 475 to 685ms following T1 stimuli (relative to the alpha-power across the rest of the epoch). Higher alpha-power has been associated with the inhibition of non-relevant brain regions during attention tasks, with the suggestion that this allows the brain to prioritise processing in brain regions that are relevant to the task, without the relevant brain regions being “distracted” by processing in non-relevant regions (Klimesch et al., 2007). In contrast, lower alpha-power is found in brain regions where active processing is required to complete the task, such that alpha-power can be increased to inhibit processing or decreased to enable processing in specific brain regions (Klimesch et al., 2007). In support of this interpretation of the function of alpha-power, previous research has shown higher levels of brain region-specific alpha-power modulation in experienced meditators when attention is required to either tactile oddball or visual working memory stimuli (Wang et al., 2020). Results in that study indicated that alpha-power increased or decreased in specific task-relevant regions, dependent on the specific task demands, that meditators produced stronger task-relevant increases or decreases, and as such, performed the task more accurately (Wang et al., 2020). The current study provides further support for the interpretation of alpha as an inhibitory mechanism, with alpha-power remaining high during distractor stimuli presentation but decreasing (releasing inhibition) earlier in short interval trials in alignment with short interval T2 processing, and decreasing later in long interval trials, in alignment with long interval T2 processing (see Figures S8-9 in the supplementary materials section 3e for demonstration of this point). This decrease in alpha-power during short interval T2 stimuli processing and increase in alpha within long interval trials during the same time period likely reflects a ‘gating’ mechanism, with both release of inhibition to process target stimuli and increase of inhibition to reduce distractor processing. Indeed, lower alpha-power RMS within a 685 to 1050ms window was strongly associated with short interval T2 correct responses (see the Supplementary Materials section 3e, Figure S12).

As such, the results of the current study might suggest that the reduction in alpha-power immediately following the timing of the presentation of short interval T2 stimuli in the meditation group reflects an attentional mechanism enabling increased neural processing during the period where neural activity would process the short interval T2 stimuli. This appears to occur regardless of whether the short interval T2 stimuli was presented or not presented (in the case that the trial was a long interval trial). Two possible interpretations of the fact that meditators showed this prolonged alpha-power reduction to enable short interval T2 processing even for long interval trials is that it may reflect a neural activity pattern prioritising either awareness in general, or carefulness. The increased processing of stimuli, regardless of whether they might be task-relevant, might reflect increased general awareness. Alternatively, the increased processing of the time period during which T2 might be present may indicate increased carefulness in anticipation of a potential T2 stimuli being presented. Some previous research has reported results that suggest the “increased awareness” interpretation is more likely - research using mathematical modelling of performance in a behavioural task has suggested that the improved attention function from mindfulness is related to enhancements in an individual’s ability to extract higher information quality during a working memory task rather than increased caution in responding (Van Vugt & Jha, 2011), a finding supported by neuroimaging research showing earlier activation of working memory related brain regions in meditators (Bailey et al., 2020). Our task did not require participants to respond quickly, so it did not provide the ability to assess reaction times. However, previous results indicated meditators have shown increased performance without reaction time slowing (Van Vugt & Jha, 2011) and increased accuracy across both fast and slow reaction times (van den Hurk et al., 2010). In contrast, other research has indicated that meditators perform better in a movement task when the action required to meet the task goals is ambiguous and changing, and that they achieve this by performing a speed-accuracy trade-off for slower but more accurate responses (Naranjo & Schmidt, 2012). Trait mindfulness has also been shown to reduce the accelerating but accuracy-reducing effects of worry on performance (Hallion et al., 2020), supporting the “increased carefulness” interpretation.

Further research may be able to elucidate the reasons for this pattern further. While this pattern whereby meditators may have shown prolonged alpha-power reduction to enable short interval T2 processing even for long interval trials and our suggested interpretations of the pattern would have had no effect on task-relevant stimulus perception and, therefore, could not lead to improved task performance, the pattern does align with the “non-judgemental” aspect of mindfulness practice – maintaining awareness of the present moment as it is, without evaluation. This contrasts with the pattern shown by the controls, which indicates they reduced the processing of non-target distractor stimuli within the short interval T2 period, eliminating the distractor stimuli from awareness. As might be expected, given the lack of relevance to task performance of this neural strategy, across all participants, averaged alpha-power within the time window where meditators showed reduced alpha activity did not correlate with the accuracy of short interval T2 detection. In fact, our exploratory analysis indicated that *incorrect* responses on short interval trials were associated with slightly, but significantly, lower alpha-power within this window than correct responses (supplementary materials section 3e, S11). This might provide support for a conjecture that the careful or non-judgemental neural strategy of the meditators prioritised present moment awareness at the expense of accurate task performance. However, alpha-power RMS was also strongly correlated between the earlier (during-T2 processing) and later (post-T2 processing) alpha power time periods, and this relationship was stronger within incorrect trials than for correct trials. As such, it may be that the alpha-power reduction during the earlier (during-T2 processing) period might reflect a preparatory mechanism that attempted to engage attention when attention had drifted, so that the neural activity required for successful task performance in the later (post-T2 processing) window would be present. We note that at this stage, these explanations are conjecture, and alternatively, it may simply be that the lower alpha-power in meditators during the earlier (during-T2 processing) period reflects a non-optimal neural activation in the context of the task. Further research may be able to determine which explanation is correct.

### Alpha Phase Synchronisation

Similar to the alpha-power results, our study did not find a significant difference in our primary analysis focused on alpha phase synchronisation time windows in replication of those reported by Slagter et al. (2009). However, in contrast with the lower alpha-power during the short interval T2 stimuli time window, the meditation group showed a prolonged period of *higher* alpha synchronisation to T1. Meditators also showed a different scalp distribution of alpha synchronisation to T1, with more parietal and frontal APS than controls. While alpha-power has been associated with the inhibition of non-relevant brain regions during attention tasks that require processing for other reasons (Klimesch et al., 2007), the same relationship has not been reported for APS. Indeed, the correlation between APS and task performance in our study, along with the more occipital distribution in the meditation group, suggests that inhibition of non-relevant brain regions (in our visual task) is not likely to be the explanation for the higher APS in our meditation group. Instead, we suspect the increased APS in our meditation group reflects synchronisation to the ongoing stream of stimuli presentation timing (as stimuli were presented at 10Hz, within the alpha frequency). Previous research has suggested that the synchronisation of ongoing endogenous neural oscillations to external stimuli may increase the likelihood of neurons firing in response to those stimuli, which is then related to the increased encoding of that stimuli into working memory (Buzsáki & Moser, 2013; Fujisawa & Buzsáki, 2011; Lisman & Buzsáki, 2008; O’Neill et al., 2013). This process is likely to reflect a mechanism underlying attention function, and a similar phenomenon may underlie the alpha synchronisation to stimuli in the current study. As such, it may be that the attentional training the meditation group had undertaken increased their ability to time lock their alpha oscillations to stimuli in occipital regions responsible for processing the visual stimuli, and frontal regions responsible for attending to the stimuli. We note here that it might be valuable to analyse connectivity between these regions in future research. However, if these explanations are accurate, it is unclear why no performance differences were found between the groups. The lack of behavioural difference between the groups was unlikely to be due to a ceiling effect, as mean accuracy in the groups was 67.06% for meditators and 63.91% for controls. As such, the exploratory APS results require replication before we can confirm these explanations.

### Potential explanations for our null results

While our results suggest differences in neural activity in meditators that align with improved attention function, the meditator and control groups did not differ in task performance. There are a number of potential explanations for this null result, as well as the null results for our primary analyses. For the sake of brevity, these are summarised here, and explained in full in the supplementary materials (section 4). Firstly, the behavioural effects of meditation in the AB task may be dependent on a meditation-induced mindful state, or particular types of meditative practices. Secondly, it may be that more meditation experience is required before differences in AB task performance are detected, or that the AB task was not sensitive enough to detect differences between our groups. Age may have also been a factor - perhaps meditation protects against age-related decline in AB performance, and our young meditation group had not aged enough to show this effect. However, these explanations seem unlikely given our meditators were more experienced than those included in many studies, our task replicated a number of previous AB task studies that did detect differences, and some research has indicated older meditators showed improved AB task performance compared to both age-matched controls *and* a younger control group (van Leeuwen et al., 2009). Next, our study design differed from Slagter et al. (2007, 2009) – their study involved the repetition of the AB task before and after an intensive retreat. It may be that MM is not associated with generalised better performance in the AB task, but rather an increased ability to learn the task and as a result, increased performance on the second repetition of the task following meditation practice. This features also meant that Slagter et al.’s (2007, 2009) within subject design also controlled for interindividual variability, while our between-groups study did not.

Our study also included updated EEG analysis methods from Slagter et al. (2007, 2009). Most notably, the current study used a high pass filter of 0.25Hz, whereas Slagter et al. (2007) used a high pass filter of 1Hz. The amplitude of ERPs, including the P3b, has been shown to be produced at least in part by < 1Hz activity, and are adversely affected by high pass filtering out < 1Hz data (Rousselet, 2012; Tanner et al., 2016). As such, the P3b data Slagter et al. (2007) analysed may have had considerable signal removed from the P3b, and their analysis may have been adversely affected. Lastly, it may be that either our result or the results reported by Slagter et al. (2007, 2009) are spurious, reflecting a sampling bias, chance-like effect, or similar “non-effect of interest”. However, we note that a spurious chance-like result is less likely in studies with a larger sample size, as per the current study (Agrillo & Petrazzini, 2012).

As such, our results indicate that the specific alterations detected by previous research, including those to the P3b (within a specific window of interest), increased T2-locked TPS, and improved performance on short interval AB trials, are not necessarily markers of regular mindfulness meditation practice. Despite the potential explanations outlined in the previous sections for the differences between the meditator and control group in our study, these findings were exploratory and were not controlled for experiment-wise multiple comparisons. As such, it is possible that there are simply no differences between groups and that ultimately, previous mindfulness experience may not result in behavioural improvements in the AB task (although unlikely given the number of positive findings, even if the findings were exploratory). Although our EEG findings are uncertain, our results provide confidence in the null result for differences in task performance. This was surprising as it conflicted with previous findings (Slagter et al., 2007; Slagter et al., 2009). It was especially surprising considering that the meditators in the current study reported at least two years of meditative practice, which we expected would be sufficient to produce differences in attention performance if MM did indeed affect attention. From our perspective, the most likely explanation for the difference between our results and those of Slagter et al. (2007, 2009) is that our participants were regular meditators, whereas theirs were tested before and after a 3-month retreat. As such, when viewing both studies together, our results suggest that differences in AB performance among meditators may be exclusively present following intensive meditation interventions.

It may be that the type of attention captured by the AB task is less relevant to the attention trained through mindfulness meditation practice. This interpretation is supported by our alpha-power findings, which suggested meditators may not have engaged alpha to inhibit distractor processing when short interval T2 stimuli were absent as strongly as the controls. Other EEG markers or neuroimaging methods using different attention tasks may be better suited to detect differences between meditation and control groups, and the null results for behavioural analyses in the current study may help refine our understanding of exactly which mechanisms are altered (and which are not altered) by meditation practice. With AB literature suffering from a lack of published replications, the present study also underscores the importance of replication studies in different populations and contexts, as some of the effects of meditation may be specific to certain populations only (Bailey et al., 2019b; Osborn et al., 2022; Vago et al., 2019; Van Dam et al., 2018). Slagter et al. (2007) have been cited over 1000 times, yet this is the first even partial replication attempt, which, despite using a larger sample size, revealed null results for our replication of the outcome measures reported by Slagter et al. (2007, 2009).

### Limitations and Future Directions

The most obvious limitation of our study is that it utilised a cross-sectional design. A longitudinal approach, assessing participants before and after meditation practice, may allow for the determination of causality. However, we note that this is difficult to achieve with the level of meditation experience tested in the current study. Another limitation of this study was that it utilised a broad definition of meditation (Kabat-Zinn, 1994) and included both “focused attention” and “open monitoring” practitioners. Meditation literature is unclear on the direct impact of different varieties of meditation practice on AB performance, with research suggesting both focused attention and open monitoring meditation affect AB performance (van Leeuwen et al., 2009), other research suggesting AB performance is exclusively impacted by open monitoring meditation (Colzato et al., 2015), and some studies suggest neither practice affects AB performance (Sharpe et al., 2021). While delineating between the different MM practices and their potential impacts may be valuable, the conclusions that can be drawn from our broad sample may be more reflective of everyday mindfulness meditators in the community. For additional strengths and limitations of the study, see the supplementary materials.

## Acknowledgements

We gratefully acknowledge the Dhamma Aloka Vipassana meditation centre in Melbourne and the Melbourne Insight Meditation centre for their assistance with the recruitment of several meditators who took part in the study.

## Data Availability Statement

The data that support the findings of this study are available from the corresponding author, NWB, upon reasonable request.

## References

Atchley, R., Klee, D., Memmott, T., Goodrich, E., Wahbeh, H., & Oken, B. (2016). Event-related potential correlates of mindfulness meditation competence. Neuroscience, 320, 83–92. https://doi.org/10.1016/j.neuroscience.2016.01.051

Baer, R. A., Smith, G. T., Hopkins, J., Krietemeyer, J., & Toney, L. (2006). Using Self-Report Assessment Methods to Explore Facets of Mindfulness. Assessment, 13(1), 27–45. https://doi.org/10.1177/1073191105283504

Bailey, N., Biabani, M., Hill, A. T., Miljevic, A., Rogasch, N. C., McQueen, B., Murphy, O. W., & Fitzgerald, P. (2022a). Introducing RELAX (the Reduction of Electroencephalographic Artifacts): A fully automated pre-processing pipeline for cleaning EEG data-Part 1: Algorithm and Application to Oscillations. bioRxiv. https://doi.org/https://doi.org/10.1101/2022.03.08.483548

Bailey, N., Freedman, G., Raj, K., Sullivan, C., Rogasch, N., Chung, S. W., Hoy, K., Chambers, R., Hassed, C., Van Dam, N., & Fitzgerald, P. (2019). Mindfulness meditators show altered distributions of early and late neural activity markers of attention in a response inhibition task. PLoS One, 14(8), e0203096. https://doi.org/10.1101/396259

Bailey, N., Geddes, H., Zannettino, I., Humble, G., Payne, J., Baell, O., Emonson, M., Chung, S. W., Hill, A. T., & Rogasch, N. C. (2022). Meditators probably show increased behaviour-monitoring related neural activity. Mindfulness, 374(14), 33–49.

Bailey, N., Hill, A., Biabani, M., Murphy, O., Rogasch, N., McQueen, B., Miljevic, A., & Fitzgerald, P. (2022b). Introducing RELAX (the Reduction of Electroencephalographic Artifacts): A fully automated pre-processing pipeline for cleaning EEG data – Part 2: Application to Event-Related Potentials. bioRxiv, 2022.2003.2008.483554. https://doi.org/https://doi.org/10.1101/2022.03.08.483554

Bailey, N. W., Freedman, G., Raj, K., Spierings, K. N., Piccoli, L. R., Sullivan, C. M., Chung, S. W., Hill, A. T., Rogasch, N. C., & Fitzgerald, P. B. (2020). Mindfulness meditators show enhanced accuracy and different neural activity during working memory. Mindfulness, 11, 1762–1781.

Bailey, N. W., Raj, K., Freedman, G., Fitzgibbon, B. M., Rogasch, N. C., Van Dam, N. T., & Fitzgerald, P. B. (2019b). Mindfulness meditators do not show differences in electrophysiological measures of error processing. Mindfulness, 10(7), 1360–1380.

Beck, Epstein, Brown, & Steer. (1988). An Inventory for Measuring Clinical Anxiety: Psychometric Properties. Journal of Consulting and Clinical Psychology, 56(6), 893. https://doi.org/10.1037/0022-006X.56.6.893

Beck, Ward, Mendelson, Mock, & Erbaugh. (1961). An Inventory for Measuring Depression. Archives of General Psychiatry, 4(6), 561–571. https://doi.org/10.1001/archpsyc.1961.01710120031004

Benjamini, Y., & Hochberg, Y. (2000). On the adaptive control of the false discovery rate in multiple testing with independent statistics. Journal of educational and Behavioral Statistics, 25(1), 60–83. https://doi.org/https://doi.org/10.3102/10769986025001060

Bigdely-Shamlo, N., Mullen, T., Kothe, C., Su, K.-M., & Robbins, K. A. (2015). The PREP pipeline: standardized preprocessing for large-scale EEG analysis. Frontiers in neuroinformatics, 9, 16. https://doi.org/https://doi.org/10.3389/fninf.2015.00016

Britton, W. B., Davis, J. H., Loucks, E. B., Peterson, B., Cullen, B. H., Reuter, L., Rando, A., Rahrig, H., Lipsky, J., & Lindahl, J. R. (2018). Dismantling Mindfulness-Based Cognitive Therapy: Creation and validation of 8-week focused attention and open monitoring interventions within a 3-armed randomized controlled trial. Behaviour Research and Therapy, 101, 92–107. https://doi.org/https://doi.org/10.1016/j.brat.2017.09.010

Buzsáki, G., & Moser, E. I. (2013). Memory, navigation and theta rhythm in the hippocampal-entorhinal system. Nature neuroscience, 16(2), 130–138. https://doi.org/https://doi.org/10.1038/nn.3304

Cahn, & Polich, J. (2009). Meditation (Vipassana) and the P3a event-related brain potential. Int J Psychophysiol, 72(1), 51–60. https://doi.org/10.1016/j.ijpsycho.2008.03.013

Castellanos, N. P., & Makarov, V. A. (2006). Recovering EEG brain signals: Artifact suppression with wavelet enhanced independent component analysis. Journal of Neuroscience Methods, 158(2), 300–312. https://doi.org/10.1016/j.jneumeth.2006.05.033

Chambers, R., Gullone, E., & Allen, N. B. (2009). Mindful emotion regulation: An integrative review. Clinical Psychology Review, 29(6), 560–572. https://doi.org/10.1016/j.cpr.2009.06.005

Cramer, H., Hall, H., Leach, M., Frawley, J., Zhang, Y., Leung, B., Adams, J., & Lauche, R. (2016). Prevalence, patterns, and predictors of meditation use among US adults: A nationally representative survey. Scientific reports, 6(1), 1–9. https://doi.org/https://doi.org/10.1038/srep36760

Crane, R. S., Brewer, J., Feldman, C., Kabat-Zinn, J., Santorelli, S., Williams, J. M. G., & Kuyken, W. (2017). What defines mindfulness-based programs? The warp and the weft. Psychological Medicine, 47(6), 990–999. https://doi.org/http://dx.doi.org/10.1017/S0033291716003317

Dell’Acqua, R., Dux, P. E., Wyble, B., Doro, M., Sessa, P., Meconi, F., & Jolicœur, P. (2015). The attentional blink impairs detection and delays encoding of visual information: Evidence from human electrophysiology. Journal of Cognitive Neuroscience, 27(4), 720–735. https://doi.org/10.1162/jocn_a_00752

Delorme, A., & Makeig, S. (2004). EEGLAB: an open source toolbox for analysis of single-trial EEG dynamics including independent component analysis. Journal of Neuroscience Methods, 134(1), 9–21. https://doi.org/https://doi.org/10.1016/j.jneumeth.2003.10.009

Di Lollo, V., Kawahara, J.-i., Ghorashi, S. S., & Enns, J. T. (2005). The attentional blink: Resource depletion or temporary loss of control? Psychological research, 69(3), 191–200. https://doi.org/10.1007/s00426-004-0173-x

Falkenstein, M., Hohnsbein, J., & Hoormann, J. (1993). Late visual and auditory ERP components and choice reaction time. Biological Psychology, 35(3), 201–224. https://doi.org/10.1016/0301-0511(93)90002-p

Falkenstein, M., Hohnsbein, J., Hoormann, J., & Blanke, L. (1991). Effects of crossmodal divided attention on late ERP components. II. Error processing in choice reaction tasks. Electroencephalography and clinical neurophysiology, 78(6), 447–455. https://doi.org/https://doi.org/10.1016/0013-4694(91)90062-9

Fitzgibbon, S., DeLosAngeles, D., Lewis, T., Powers, D., Grummett, T., Whitham, E., Ward, L., Willoughby, J., & Pope, K. (2016). Automatic determination of EMG-contaminated components and validation of independent component analysis using EEG during pharmacologic paralysis. Clinical Neurophysiology, 127(3), 1781–1793. https://doi.org/10.1016/j.clinph.2015.12.009

Fujisawa, S., & Buzsáki, G. (2011). A 4 Hz oscillation adaptively synchronizes prefrontal, VTA, and hippocampal activities. Neuron, 72(1), 153–165. https://doi.org/10.1016/j.neuron.2011.08.018

Habermann, M., Weusmann, D., Stein, M., & Koenig, T. (2018). A Student’s Guide to Randomization Statistics for Multichannel Event-Related Potentials Using Ragu. Frontiers in Neuroscience, 12. https://doi.org/10.3389/fnins.2018.00355

Hallion, L. S., Kusmierski, S. N., & Caulfield, M. K. (2020). Worry alters speed-accuracy tradeoffs but does not impair sustained attention. Behaviour Research and Therapy, 128, 103597. https://doi.org/https://doi.org/10.1016/j.brat.2020.103597

Hayes, S. C. (2012). Acceptance and commitment therapy the process and practice of mindful change (2nd ed. ed.). New York : Guilford Press.

Hölzel, B. K., Lazar, S. W., Gard, T., Schuman-Olivier, Z., Vago, D. R., & Ott, U. (2011). How Does Mindfulness Meditation Work? Proposing Mechanisms of Action From a Conceptual and Neural Perspective. Perspectives on Psychological Science, 6(6), 537–559. https://doi.org/10.1177/1745691611419671

Jha, A. P., Krompinger, J., & Baime, M. J. (2007). Mindfulness training modifies subsystems of attention. Cognitive, Affective, & Behavioral Neuroscience, 7(2), 109–119. https://doi.org/10.3758/CABN.7.2.109

Kabat-Zinn, J. (1994). Wherever you go, there you are : mindfulness meditation in everyday life (1st ed. ed.). New York : Hyperion.

Kiken, L. G., Garland, E. L., Bluth, K., Palsson, O. S., & Gaylord, S. A. (2015). From a state to a trait: Trajectories of state mindfulness in meditation during intervention predict changes in trait mindfulness. Personality and Individual Differences, 81, 41–46. https://doi.org/https://doi.org/10.1016/j.paid.2014.12.044

Klimesch, W. (2012). α-band oscillations, attention, and controlled access to stored information. Trends in Cognitive Sciences, 16(12), 606–617. https://doi.org/10.1016/j.tics.2012.10.007

Klimesch, W., Sauseng, P., & Hanslmayr, S. (2007). EEG alpha oscillations: the inhibition– timing hypothesis. Brain research reviews, 53(1), 63–88. https://doi.org/10.1016/j.brainresrev.2006.06.003

Koenig, T., Kottlow, M., Stein, M., & Melie-García, L. (2011). Ragu: A Free Tool for the Analysis of EEG and MEG Event-Related Scalp Field Data Using Global Randomization Statistics. Computational Intelligence and Neuroscience, 2011, 938925. https://doi.org/10.1155/2011/938925

Koenig, T., & Melie-garcía, L. (2010). A Method to Determine the Presence of Averaged Event-Related Fields Using Randomization Tests. Brain Topography, 23(3), 233–242. https://doi.org/http://dx.doi.org/10.1007/s10548-010-0142-1

Kuyken, W., Byford, S., Taylor, R. S., Watkins, E., Holden, E., White, K., Barrett, B., Byng, R., Evans, A., Mullan, E., & Teasdale, J. D. (2008). Mindfulness-Based Cognitive Therapy to Prevent Relapse in Recurrent Depression. Journal of Consulting and Clinical Psychology, 76(6), 966–978. https://doi.org/10.1037/a0013786

Lachaux, J.-P., Rodriguez, E., Martinerie, J., & Varela, F. J. (1999). Measuring phase synchrony in brain signals. Human Brain Mapping, 8(4), 194–208. https://doi.org/10.1002/(SICI)1097-0193(1999)8:4

Lisman, J., & Buzsáki, G. (2008). A neural coding scheme formed by the combined function of gamma and theta oscillations. Schizophrenia bulletin, 34(5), 974–980. https://doi.org/10.1093/schbul/sbn060

Love, J., Selker, R., Marsman, M., Jamil, T., Dropmann, D., Verhagen, J., Ly, A., Gronau, Q. F., Šmíra, M., & Epskamp, S. (2019). JASP: Graphical statistical software for common statistical designs. Journal of Statistical Software, 88, 1–17. https://doi.org/10.18637/jss.v088.i02

Lutz, Slagter, Dunne, & Davidson. (2008). Attention regulation and monitoring in meditation. Trends in Cognitive Sciences, 12(4), 163–169. https://doi.org/10.1016/j.tics.2008.01.005

Lutz, Slagter, H., A, Rawlings, N., B, Francis, D., A, Greischar, L., L, & Davidson, J., D. (2009). Mental Training Enhances Attentional Stability: Neural and Behavioral Evidence. The Journal of Neuroscience, 29(42), 13418 –13427. https://doi.org/10.1523/JNEUROSCI.1614-09.2009

Martens, S., & Wyble, B. (2010). The attentional blink: past, present, and future of a blind spot in perceptual awareness. Neuroscience and biobehavioral reviews, 34(6), 947–957. https://doi.org/10.1016/j.neubiorev.2009.12.005

Mizuhara, H., & Yamaguchi, Y. (2007). Human cortical circuits for central executive function emerge by theta phase synchronization. Neuroimage, 36(1), 232–244. https://doi.org/10.1016/j.neuroimage.2007.02.026

Naranjo, J. R., & Schmidt, S. (2012). Is it me or not me? Modulation of perceptual-motor awareness and visuomotor performance by mindfulness meditation. BMC neuroscience, 13(1), 1–17. https://doi.org/https://doi.org/10.1186/1471-2202-13-88

O’Neill, P.-K., Gordon, J. A., & Sigurdsson, T. (2013). Theta oscillations in the medial prefrontal cortex are modulated by spatial working memory and synchronize with the hippocampus through its ventral subregion. Journal of Neuroscience, 33(35), 14211–14224. https://doi.org/10.1523/JNEUROSCI.2378-13.2013

Olivers, C. N., & Meeter, M. (2008). A boost and bounce theory of temporal attention. Psychological review, 115(4), 836. https://doi.org/10.1037/a0013395

Oostenveld, R., Fries, P., Maris, E., & Schoffelen, J.-M. (2011). FieldTrip: Open Source Software for Advanced Analysis of MEG, EEG, and Invasive Electrophysiological Data. Computational Intelligence and Neuroscience, 2011(2011). https://doi.org/10.1155/2011/156869

Osborn, M., Shankar, S., Szymanski, O., Gunningham, K., Caldwell, B., Perera, M. P. N., Michael, J., Wang, M., Fitzgerald, P. B., & Bailey, N. W. (2022). Meta-analysis Provides Weak Evidence for an Effect of Mindfulness on Neural Activity Related to Error-Processing in Healthy Individuals Only. Mindfulness, 1–25.

Pion-Tonachini, L., Kreutz-Delgado, K., & Makeig, S. (2019). ICLabel: An automated electroencephalographic independent component classifier, dataset, and website. Neuroimage, 198, 181–197. https://doi.org/https://doi.org/10.1016/j.neuroimage.2019.05.026

Polich, J. (1997). EEG and ERP assessment of normal aging. Electroencephalography and Clinical Neurophysiology/Evoked Potentials Section, 104(3), 244–256. https://doi.org/10.1016/s0168-5597(97)96139-6

Potter, M. C., Chun, M. M., Banks, B. S., & Muckenhoupt, M. (1998). Two attentional deficits in serial target search: the visual attentional blink and an amodal task-switch deficit. Journal of Experimental Psychology: Learning, Memory, and Cognition, 24(4), 979. https://doi.org/10.1037//0278-7393.24.4.979

Raimondo, F., Kamienkowski, J. E., Sigman, M., & Fernandez Slezak, D. (2012). CUDAICA: GPU optimization of infomax-ICA EEG analysis. Computational Intelligence and Neuroscience, 2012. https://doi.org/https://doi.org/10.1155/2012/206972

Rousselet, G. A. (2012). Does filtering preclude us from studying ERP time-courses? Frontiers in psychology, 3, 131. https://doi.org/10.3389/fpsyg.2012.00131

Sergent, C., Baillet, S., & Dehaene, S. (2005). Timing of the brain events underlying access to consciousness during the attentional blink. Nature neuroscience, 8(10), 1391–1400. https://doi.org/https://doi.org/10.1038/nn1549

Shapiro, K. L., Raymond, J. E., & Arnell, K. M. (1997). The attentional blink. Trends in Cognitive Sciences, 1(8), 291–296. https://doi.org/10.1016/S1364-6613(97)01094-2

Sharpe, P., Whalley, B., & Mitchell, C. J. (2021). Does brief focused attention and open monitoring meditation affect the attentional blink? Mindfulness, 12(10), 2430–2438. https://doi.org/https://doi.org/10.1007/s12671-021-01709-2

Sheehan, D. V., Lecrubier, Y., Sheehan, K. H., Amorim, P., Janavs, J., Weiller, E., Hergueta, T., Baker, R., & Dunbar, G. C. (1998). The Mini-International Neuropsychiatric Interview (MINI): the development and validation of a structured diagnostic psychiatric interview for DSM-IV and ICD-10. Journal of clinical psychiatry, 59(20), 22–33. https://doi.org/https://pubmed.ncbi.nlm.nih.gov/9881538/

Slagter, H. A., Lutz, A., Greischar, L. L., Francis, A. D., Nieuwenhuis, S., Davis, J. M., & Davidson, R. J. (2007). Mental Training Affects Distribution of Limited Brain Resources. PLoS Biology, 5(6), e138. https://doi.org/10.1371/journal.pbio.0050138

Slagter, H. A., Lutz, A., Greischar, L. L., Nieuwenhuis, S., & Davidson, R. J. (2009). Theta Phase Synchrony and Conscious Target Perception: Impact of Intensive Mental Training. Journal of Cognitive Neuroscience, 21(8), 1536–1549. https://doi.org/10.1162/jocn.2009.21125

Somers, B., Francart, T., & Bertrand, A. (2018). A generic EEG artifact removal algorithm based on the multi-channel Wiener filter. Journal of neural engineering, 15(3), 036007. https://doi.org/10.1088/1741-2552/aaac92

Tang, Hölzel, B. K., & Posner, M.. (2015). The neuroscience of mindfulness meditation. Nature Reviews. Neuroscience, 16(4), 213–225. https://doi.org/http://dx.doi.org/10.1038/nrn3916

Tang, Ma, Y., Wang, J., Fan, Y., Feng, S., Lu, Q., Yu, Q., Sui, D., Rothbart, M. K., Fan, M., & Posner, M. I. (2007). Short-term meditation training improves attention and self-regulation.(PSYCHOLOGY)(Clinical report). Proceedings of the National Academy of Sciences of the United States, 104(43), 17152. https://doi.org/10.1073/pnas.0707678104

Tanner, D., Norton, J. J., Morgan-Short, K., & Luck, S. J. (2016). On high-pass filter artifacts (they’re real) and baseline correction (it’s a good idea) in ERP/ERMF analysis. Journal of Neuroscience Methods, 266, 166–170. https://doi.org/10.1016/j.jneumeth.2016.01.002

Ueno, T., Hirano, S., Hirano, Y., Kanba, S., Kobayashi, S., & Onitsuka, T. (2009). Locked to Stimulation: Significance Level of the Phase-Locking Factor. 2nd International Congress on Image and Signal Processing, 1–4. https://doi.org/10.1109/CISP.2009.5304010

Vago, D. R., Gupta, R. S., & Lazar, S. W. (2019). Measuring cognitive outcomes in mindfulness-based intervention research: a reflection on confounding factors and methodological limitations. Current opinion in psychology, 28, 143–150. https://doi.org/10.1016/j.copsyc.2018.12.015

Van Dam, N. T., van Vugt, M. K., Vago, D. R., Schmalzl, L., Saron, C. D., Olendzki, A., Meissner, T., Lazar, S. W., Kerr, C. E., Gorchov, J., Fox, K. C. R., Field, B. A., Britton, W. B., Brefczynski-Lewis, J. A., & Meyer, D. E. (2018). Mind the Hype: A Critical Evaluation and Prescriptive Agenda for Research on Mindfulness and Meditation. Perspectives on Psychological Science, 13(1), 36–61. https://doi.org/10.1177/1745691617709589

van den Hurk, P. A., Giommi, F., Gielen, S. C., Speckens, A. E., & Barendregt, H. P. (2010). Greater efficiency in attentional processing related to mindfulness meditation. Quarterly journal of experimental psychology, 63(6), 1168–1180. https://doi.org/10.1080/17470210903249365

van Leeuwen, S., Müller, N. G., & Melloni, L. (2009). Age effects on attentional blink performance in meditation. Consciousness and Cognition, 18(3), 593–599. https://doi.org/https://doi.org/10.1016/j.concog.2009.05.001

Van Vugt, M. K., & Jha, A. P. (2011). Investigating the impact of mindfulness meditation training on working memory: A mathematical modeling approach. Cognitive, Affective, & Behavioral Neuroscience, 11(3), 344–353. https://doi.org/10.3758/s13415-011-0048-8

Varela, F., Lachaux, J.-P., Rodriguez, E., & Martinerie, J. (2001). The brainweb: Phase synchronization and large-scale integration [Review Article]. Nature Reviews Neuroscience, 2, 229. https://doi.org/10.1038/35067550

Vogel, E. K., Luck, S. J., & Shapiro, K. L. (1998). Electrophysiological evidence for a postperceptual locus of suppression during the attentional blink. Journal of Experimental Psychology: Human Perception and Performance, 24(6), 1656. https://doi.org/10.1037//0096-1523.24.6.1656

Wang, M. Y., Freedman, G., Raj, K., Fitzgibbon, B. M., Sullivan, C., Tan, W.-L., Van Dam, N., Fitzgerald, P. B., & Bailey, N. W. (2020). Mindfulness meditation alters neural activity underpinning working memory during tactile distraction. Cognitive, Affective, & Behavioral Neuroscience, 20, 1216–1233. https://doi.org/https://doi.org/10.3758/s13415-020-00828-y

Wang, Y., Xiao, L., Gong, W., Chen, Y., Lin, X., Sun, Y., Wang, N., Wang, J., & Luo, F. (2021). Mindful non-reactivity is associated with improved accuracy in attentional blink testing: A randomized controlled trial. Current Psychology, 1–13. https://doi.org/https://doi.org/10.1007/s12144-021-01377-4

Ward, R., Duncan, J., & Shapiro, K. (1996). The Slow Time-Course of Visual Attention. Cognitive Psychology, 30(1), 79–109. https://doi.org/10.1006/cogp.1996.0003

Zivony, A., Allon, A. S., Luria, R., & Lamy, D. (2018). Dissociating between the N2pc and attentional shifting: An attentional blink study. Neuropsychologia, 121, 153–163. https://doi.org/https://doi.org/10.1016/j.neuropsychologia.2018.11.003

