## Supplementary Materials for "Experienced Meditators Show Multifaceted Attention-Related Differences in Neural Activity"

### Supplementary Materials Section 1 - Introductory Points

It is worth noting that the degree to which the attentional blink (AB) is reduced by mindfulness meditation (MM) appears to be affected by different contextual and individual characteristics of MM practitioners. For example, previous studies have found MM-related changes in the AB to be influenced by the age of meditators (van Leeuwen et al., 2009), type of MM (van Vugt & Slagter, 2014), trait mindfulness (Makowski et al., 2019), meditative state (Colzato et al., 2015; May et al., 2011), and the length of MM training (Roca & Vazquez, 2020; Slagter et al., 2007; Wang et al., 2021).

It is also worth noting that evidence supporting one analytical model of the AB phenomenon does not necessarily negate the mechanisms and functional processes proposed by another. Indeed, it may be that because each of these conceptual constructs view attention as a unitary phenomenon and thus may be prematurely assuming that there is a singular process underlying attention, they are all at least partially incorrect. Recent developments in attentional theory have been provided by Hommel et al. (2019), who argue for a synthetic approach to attention in which the priority is no longer delineating between neural functions, but exploring the many cognitive and functional mechanisms that contribute to and synthesise under the umbrella of ‘attention’. This perspective also aligns with that of the predictive coding framework, which views brain function (and resulting cognitive processes) as being driven simply by the minimisation of prediction errors, occurring when there is an energetically expensive mismatch between the brain’s prior model of its environment and the posterior evidence it obtains from its environment through sensory apparatus (Friston, 2010; Hohwy, 2012). While limited research has examined the AB through these lenses, we consider them to be important for understanding attention function, and mindfulness research is increasingly considering the predicting coding perspective (Deane et al., 2020; Laukkonen & Slagter, 2021; Lutz et al., 2019; Manjaly & Iglesias, 2020; Verdonk & Trousselard, 2021).

### Supplementary Materials Section 2 - Methods

Note also that the participants in this study comprise a separate sample from the participants included in our previous studies: [BLINDED FOR REVIEW].

#### Section 2a - Task Visual Specifications

Each trial started with a 1780ms fixation cross, followed by a rapid serial stream of 19 stimuli. Each stream contained distracter stimuli and one or two target stimuli. Distractor stimuli were uppercase letters randomly drawn from the alphabet (excluding B, I, J, O, Q and S due to visual similarity to numbers). Target stimuli were numbers randomly drawn from the set 2-9. Stimuli were 8.25cm (height) by 6.75 to 10.5cm (width) and were presented in black on a grey background. Each stimulus was presented for 66.67ms followed by a 33.33ms blank screen on a monitor with a 60Hz refresh rate.

#### Section 2b - TPS and APS Computation

To calculate the theta-phase synchronisation (TPS) and alpha-phase synchronisation (APS), phase locking factor values were calculated by first submitting the EEG signal from each accepted epoch to a Morlet Wavelet Transform with a Fourier output (5 oscillation cycles with steps of 1Hz and 5ms resolution) in the theta frequency range (4 to 8.5Hz) and alpha frequency range (8.5 to 15Hz, in replication of Slagter et al., 2009). We then divided the Fourier spectrum by the amplitude, summed the angles, then took the absolute value of this sum and divided it by the number of trials in the specific condition and participant to obtain TPS (Delorme & Makeig, 2004).

#### Section 2c - ERP Statistical Comparisons

Note that while Slagter et al. (2007) analysed a 350 to 650ms period, all of their significant effects were found prior to 600ms, providing the rationale for the window selected in our study. In addition to the comparisons reported in the main manuscript, to assess the likelihood of our explanations for a significant interaction between group and target, we conducted a small number of exploratory generalised linear mixed models of averaged GFP from within a significant time window including individual epochs from all participants, with correct and incorrect response and group as fixed effects factors and participant as the random effect factor.

#### Section 2d - Theta Phase Synchronisation Statistical Comparisons

To compare TPS between the groups, root mean squared (RMS) and TANOVA tests were used to conduct repeated measures ANOVA design, examining 2 group (meditators vs controls) x 2 condition (short and long interval) comparisons for TPS data surrounding T2 onset. To make comparisons with Slagter et al. (2009), RMS and TANOVA tests were averaged within the 121 to 501ms window (where Slagter et al. 2009 detected an effect that was maximal at electrodes FC6 and Fz) and the 309 to 558ms window (where Slagter et al. 2009 detected an effect that was maximal at electrode T8) after the T2 stimuli. An additional exploratory analysis was performed, including T1 stimuli in a repeated measures ANOVA design examining 2 groups (meditators vs controls) x 2 conditions (short and long interval) x 2 conditions (T1 and T2) for TPS data from -500 to 1500ms around the stimuli to determine if any effects were missed by the analysis focused only on T2.

#### Section 2e - Alpha phase synchronisation and alpha-power statistical comparisons

To replicate the comparisons of alpha-power and alpha phase synchronisation (APS) conducted by Slagter et al. (2009), RMS and TANOVA tests were used to conduct repeated measures ANOVA design comparisons of these measures, examining 2 group (meditators vs controls) x 2 condition (short and long interval) comparisons for data -500 to 1500ms surrounding T1 onset. Slagter et al. (2009) reported reductions in APS from -414 to -214ms before T1 stimuli in meditators from pre- to post-retreat (maximal at Oz). They also reported increases in alpha-power between -31 to 160ms around T1 stimuli in meditators from pre- to post-retreat (maximal at POz). We did not analyse alpha-power time-locked to T2, as the epochs that were time-locked to T1 were long enough to include the T2 stimuli, and Slagter et al. (2009) did not test differences in alpha-power time-locked to T2 stimuli. Finally, we conducted a small number of exploratory generalised linear mixed models of RMS alpha-power from individual epochs from all participants, including correct and incorrect response and group as fixed effects factors and participant as the random effect factor, in order to assess the likelihood of our explanations for significant group effects.

#### Section 2f - Single Electrode Replication Statistical Comparisons

In addition to the RAGU analysis, traditional single electrode comparisons were conducted for comparison with previous research. ERP data from electrode Pz was averaged during the early phase of the P3b between 394 and 450ms and from electrode CP3 during its later phase between 488 and 551ms post T1 onset. TPS data from electrodes Fz and FC6 were averaged between 121 to 500ms, and from electrode T8 averaged from 309 to 558ms post T2_._ The statistical program JASP (JASP Team, 2019) was used to perform single electrode analyses. Bayesian as well as frequentist repeated measures ANOVAs were used to conduct a 2 group (meditators vs controls) x 2 response (short vs long interval) x 2 electrode (Pz and CP3) comparison in direct replication of Slagter et al. (2007). Similarly, a Bayesian as well as frequentist repeated measures ANOVA was used to conduct a 2 group (meditators vs controls) x 2 response (short vs long interval) comparison for TPS at Fz and FC6 during the 121 to 500ms window and from electrode T8 during the 309 to 558ms post T2 window separately in direct replication of Slagter et al. (2009). Sphericity violations were resolved through the Greenhouse-Geisser correction (Greenhouse & Geisser, 1959). Bayes Factor analyses were used to calculate the probability of the null hypothesis in contrast to the alternative hypothesis, where null results were found (Rouder et al., 2017). The suggested comparison between models containing a hypothesised effect to equivalent models stripped of the effect (excluding higher-order interactions) was performed for these analyses.

### Supplementary Materials Section 3 - Results

#### Section 3a - Behavioural and epochs included analyses

The originally planned parametric analyses of the behavioural data showed the same results as the robust statistics, with no difference between groups found in overall percentage correct (F(1,59) = 0.698, p = 0.407, η^2^G = 0.006, BFexcl = 4.140), nor were interactions found between group and target (F(1,59) = 0.149, p = 0.700, η^2^G = 6.411e – 4, Bfexcl = 4.197), group and interval (F(1,59) = 0.098, p = 0.755, η^2^G = 1.898e – 4, Bfexcl = 3.957), nor between group, target, and interval (F(1,59) = 0.029, p = 0.866, η^2^G = 4.973e – 5, Bfexcl = 4.019). Overall, Bayesian statistical models including group or interactions including group as a factor were 259.326 less likely than the model that only included target, interval, and the interaction between target and interval (BF01 = 259.326). This suggests it is highly unlikely that the meditation group showed higher percentage correct in any condition than the control group. Means and standard deviations are presented in Table 2 of the main manuscript, and the data can be viewed in Figure 1 of the main manuscript. Similarly, no difference was found between groups for the difference in percentage correct between short on long interval responses to T2 (t(59) = 0.284, p = 0.777, Cohen’s d = 0.073, BF01 = 3.708). Additionally, because Slagter et al. (2017) focused on the percentage correct for T2 triggers specifically, an analysis was run excluding T1 trigger percentage correct. This also showed no main effect of group (F(1,59) = 0.453, p = 0.503, η^2^G = 0.006, BFexcl = 2.519) and no interaction between interval and group (F(1,59) = 0.062, p = 0.804, η^2^G = 2.368e-4, BFexcl = 3.827). Finally, while the groups did not significantly differ in age and years of education, they were not directly matched. In order to address the potential effects of these demographic variables, an analysis of covariance (ANCOVA) analysis was performed comparing short interval T2 percentage correct between the groups, including these variables as covariates. No significant differences were present between the groups (F(1,57) = 1.059, p = 0.308, η^2^G = 0.018, BFexcl = 2.968). Nor were there any significant effects of age or years of education (both p > 0.11).

In order to confirm that a difference in the number of epochs from each group did not affect our results, we compared the number of epochs from each group for each condition. A repeated measures ANOVA of the number of epochs remaining after incorrect responses and artifacts were excluded including group, interval and target as factors showed no differences or interactions involving group (all p > 0.14, BF01 = 177.523 when comparing the model including group and interactions involving group to the null hypothesis). The results for these analyses can be viewed in Tables S1 and S2.

Table S1. Statistics for the number of epochs provided by each participant for each condition. Note there was no significant main effect of group or interaction involving group.

| Effect | F | p-value |
| --- | --- | --- |
| Target | 151.493 | < 0.001 |
| Target * Group | 1.209 | 0.276 |
| Interval | 114.486 | < 0.001 |
| Interval * Group | 0.137 | 0.713 |
| Target * Interval | 84.263 | < 0.001 |
| Target * Interval * Group | 0.002 | 0.968 |

Table S2. Descriptive statistics for the number of epochs provided by each participant for each condition.

| Interval | Interval | Group | Mean | SD |
| --- | --- | --- | --- | --- |
| Short | T1 | Controls | 161.100 | 22.497 |
|  |  | Meditators | 168.226 | 14.160 |
|  | T2 | Controls | 114.033 | 36.274 |
|  |  | Meditators | 123.194 | 37.349 |
| Long | T1 | Controls | 54.300 | 8.789 |
|  |  | Meditators | 56.484 | 4.404 |
|  | T2 | Controls | 45.433 | 11.773 |
|  |  | Meditators | 49.323 | 9.053 |

In order to assess our potential explanations for our results, we conducted a generalised linear mixed model analysis with the binomial family and logit link function based on single trials with correct/incorrect response as the dependent variable, trial number and group as fixed effect factors, and participants as the random effect factor. This analysis showed that participants significantly improved their performance as the task progressed (see Tables S3 and S4).

Table S3. Generalised linear mixed model analysis results of behavioural performance (accuracy of responding to the second target stimuli in short interval trials) against trial number.

| Effect | ChiSq | p-value |
| --- | --- | --- |
| Group | 0.065 | 0.799 |
| Trial Number | 35.341 | < 0.001 |
| Group * Trial Number | 1.876 | 0.171 |

Table S4. Estimated marginal means from the generalised linear mixed model analysis results of behavioural performance (proportion of correct responses to the second target stimuli in short interval trials) against trial number. Note that estimates are on the response scale, and that both groups show improved performance as the trials progress.

|  |  |  |  | 95% CI | |
| --- | --- | --- | --- | --- | --- |
| Group | Trial Number | Estimate (SIT2 response accuracy in proportion correct) | Standard Error | Lower Bound | Upper Bound |
| Controls | 41.383 | 0.704 | 0.048 | 0.602 | 0.789 |
| Meditators | 41.383 | 0.736 | 0.044 | 0.640 | 0.813 |
| Controls | 98.593 | 0.745 | 0.045 | 0.647 | 0.823 |
| Meditators | 98.593 | 0.790 | 0.039 | 0.704 | 0.857 |
| Controls | 155.804 | 0.781 | 0.043 | 0.685 | 0.854 |
| Meditators | 155.804 | 0.836 | 0.034 | 0.758 | 0.893 |

#### Section 3b - Posterior-N2 and P3b

The topographical consistency test (TCT) demonstrated consistent neural activity, except for a short period around 320ms following both short interval T1 and T2 targets, and long interval T1 targets for the control participants, and for long interval T1 targets only for meditators, none of which overlapped with the significant effects we detected (see Supplementary Figure S1). The ERP TANOVA averaged showed no main effect of Group nor any interaction involving Group that passed global duration controls (all p > 0.05, see Supplementary Figure S2).

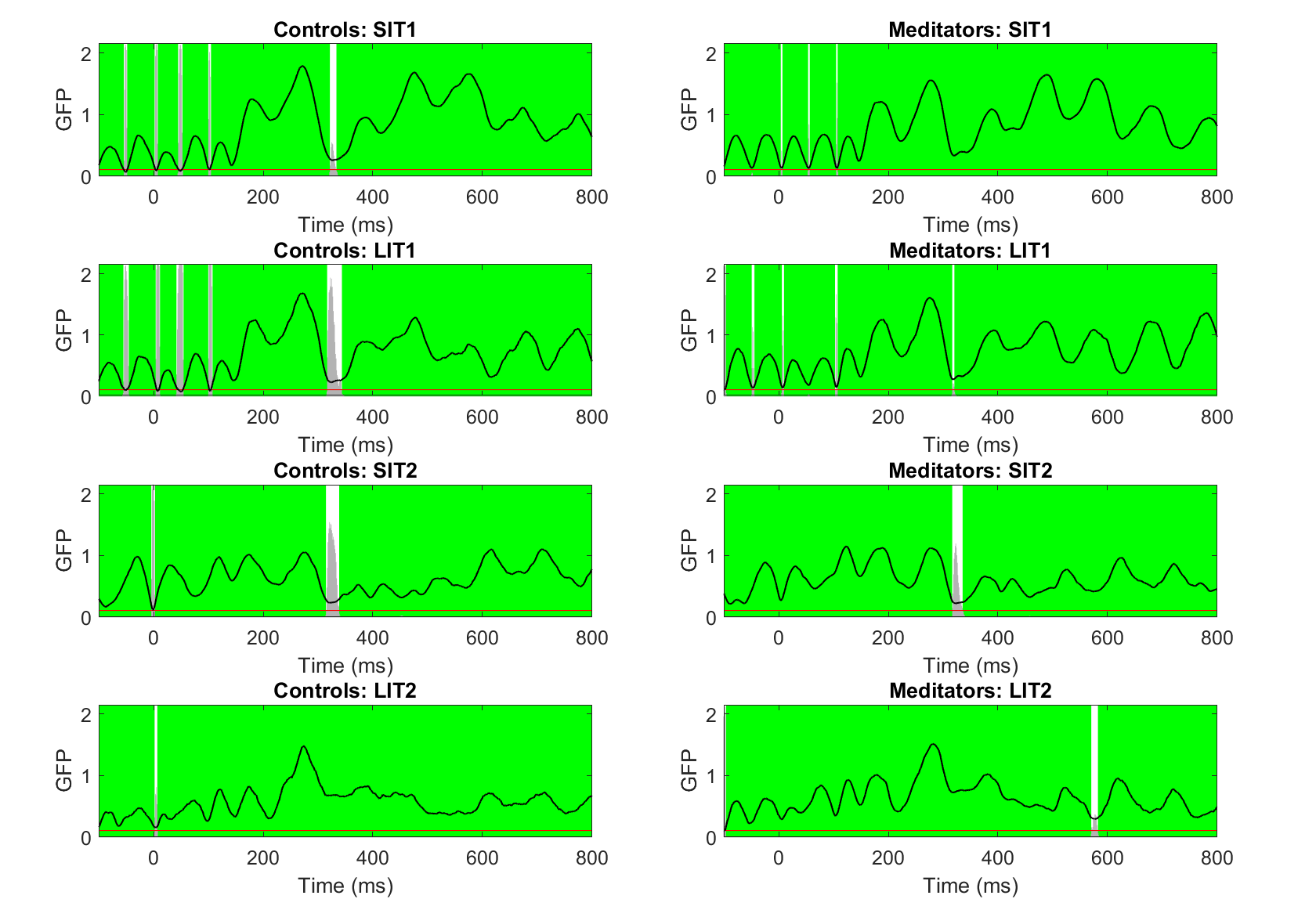

Figure S1. Topographical consistency test (TCT) results for event-related potential (ERP) data. The black line represents global field potential (GFP) values across the epoch. Green periods reflect periods of significant topographical consistency within the group/condition. Grey bars reflect the p-value, with the red line at the bottom indicating the p < 0.05 level. SIT1 reflects short interval RMS TPS following the first target stimuli (T1), SIT2 reflects short interval RMS TPS following the second target stimuli (T2), LIT1 reflects long interval RMS TPS following the first target stimuli (T1), LIT2 reflects long interval RMS TPS following the second target stimuli (T2). All groups and conditions showed within group topographical consistency with the exception of brief periods that did not overlap with our significant effect of interest. Note the oscillatory pattern in the GFP line, which oscillates within the alpha frequency, and as such demonstrates synchronisation of neural activity to the stimuli presentation rate.

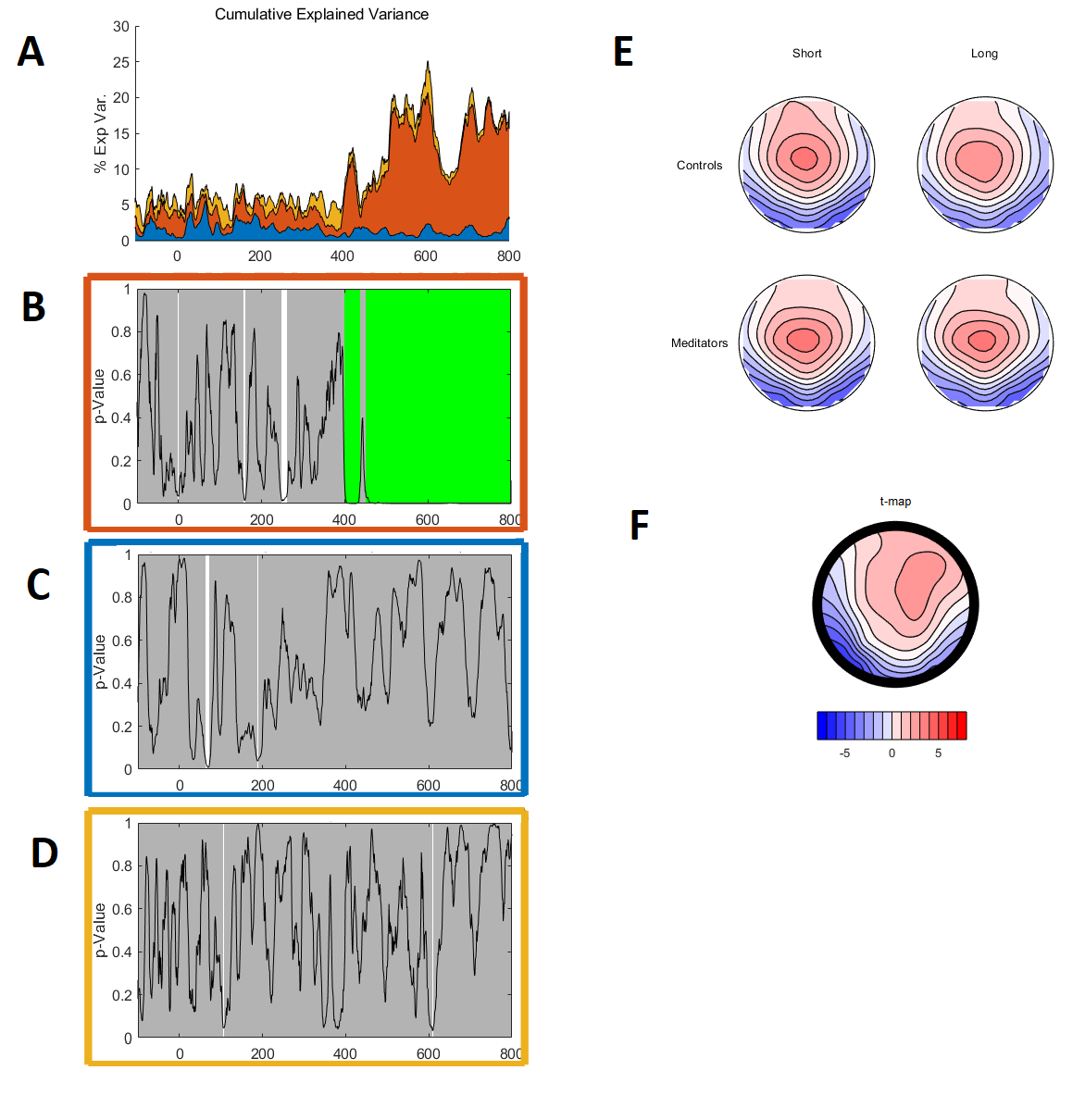

Figure S2. Topographical analysis of variance (TANOVA) for ERP comparisons following T1 stimuli for each interval condition (short and long) and each group. A. The cumulative variance explained (η_p_^2^) at each time point across the epoch by each main effect and condition, with each colour reflecting the η_p_^2^ from the effect being tested, colour coded to match the p-graphs. B. p-graph for the main effect of interval. The black line reflects the p-value, white areas reflect significant time points, and green periods reflect windows where the effect passed global duration controls. C. p-graph for the main effect of group. D. p-graph for the interaction between group and interval. E. topographical maps averaged within the P3b period of interest (350 to 600ms) for each condition and group. F. t-map for the difference between short and long interval T1 trial responses averaged within the P3b period of interest after averaging across meditators and controls.

To test whether the lack of significant differences in the GFP test or TANOVA of the P3b was due to the inclusion of all electrodes in the analyses, we restricted our analyses to the specific electrodes and time windows showing the maximum effect in Slagter et al. (2007) (Pz from 394 to 450ms and CP3 from 488 to 551ms). At Pz, this analysis showed no significant main effect of group (F(1,59) = 1.290, p = 0.261, η^2^G = 0.020, BFexcl = 1.269), nor interaction between group and interval (F(1,59) = 1.903, p = 0.173, η^2^G = 0.002, BFexcl = 1.756). The model, including interval, group, and interaction between the two, was 3.459 times less likely than the null model (BF01 = 3.459). Similar null results were found at CP3, with the main effect of group showing no significant difference (F(1,59) = 0.146, p = 0.704, η^2^G = 0.002, BFexcl = 1.843), and no significant interaction between group and interval (F(1,59) = 0.168, p = 0.683, η^2^G = 1.950e-4, BFexcl = 3.706). The overall model including interval, group, and interaction between the two was 32.747 times less likely than the null model (BF01 = 32.747).

In order to assess whether the posterior-N2 activity that showed a significant interaction between group and target in our exploratory ERP analyses was related to task accuracy (as previous research has suggested it reflects attentional engagement, (Zivony et al., 2018)), a generalised linear mixed model was conducted to assess the relationship between correct/incorrect responses from single trials, and single trial posterior-N2 GFP from both short interval T1 and T2 stimuli. This model included participant as the random effect variable, group and averaged GFP during the posterior-N2 period (227 to 254ms) following short interval T1 and T2 stimuli response type as the fixed effects variables, and correct/incorrect identification of the short interval T2 stimuli as the dependent variable. The analysis indicated that correct responses were associated with lower T1 posterior-N2 GFP (ChiSq = 125.32, p < 0.001, see Figure S3 and Table S5). Additionally, the analysis indicated that correct responses were also associated with lower T2 posterior-N2 GFP, although this relationship was very small (ChiSq = 19.00, p < 0.001, see Figure S4 and Table S5). Interestingly, the analysis indicated there was an interaction between posterior-N2 GFP from T1 and T2 (ChiSq = 29.82, p < 0.001), such that if posterior-N2 T1 values were low, then low posterior-N2 T2 values were associated with the best performance (see Table S5). However, if posterior-N2 T1 values were high, then high posterior-N2 T2 values were associated with the best performance. This suggests that perhaps the brain efficiently allocates neural resources, such that if it can accurately identify T2 stimuli with a low amplitude posterior N2 response to T2 because it has minimally engaged these posterior N2 attention resources to T1, then it does so. In contrast, if the posterior N2 to T1 was high in amplitude, suggesting over engagement of attentional resources to that earlier stimulus, then the posterior N2 response to T2 also needs to be large to enable processing of this stimuli.

Finally, to assess whether the relationship between posterior-N2 and accuracy changed over time, perhaps reflecting a learning mechanism, a generalised linear mixed model was conducted with response as the dependent variable, short interval T1 posterior-N2, short interval T2 posterior-N2, group, and trial number as fixed effect variables, and participant as the random effect variable. Notably, this analysis showed an interaction between group, short interval T2 posterior-N2, and trial number (see Tables S6, S7 and S8). The estimated marginal means from this analysis showed that, to begin with, both groups showed better performance with both low T1 posterior-N2 GFP values and low T2 posterior-N2 GFP values. This pattern was maintained for the control group, but for the meditation group, the pattern switched so that while low T1 posterior-N2 GFP values were still associated with correct responses, high T2 posterior-N2 GFP values were associated with correct responses in the later trials (Tables S7 and S8). Since the posterior-N2 is associated with attention engagement to stimuli, it may be that meditators learned to use this attention engagement to improve their performance as the task progressed.

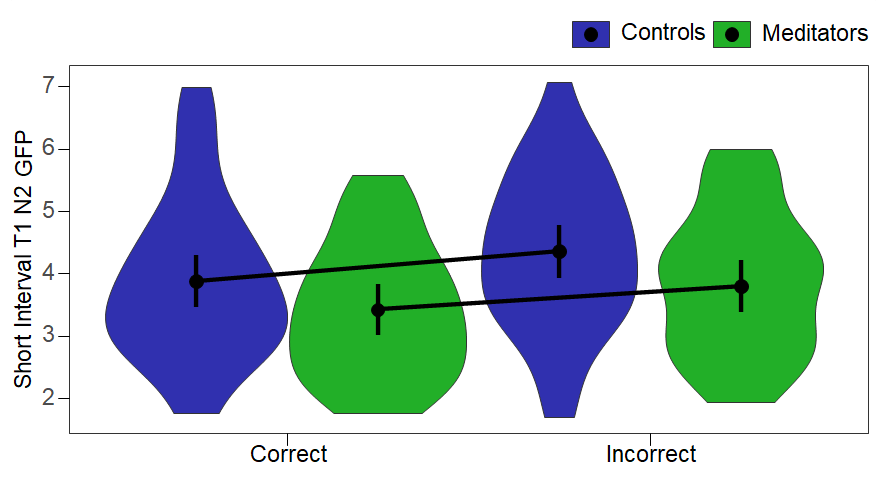

Figure S3. Estimated marginal means for posterior-N2 global field potential (GFP) to short interval T1 trials, averaged across the time period of significant difference between meditators and controls (227 to 254ms), for both groups and split by whether the T2 stimuli were responded to correctly. Note that the GFP values were extracted from each short interval T2 single epoch separately, and estimated marginal means were calculated from these single trials. As such, these values are higher than those presented for the analysis within RAGU, which averaged the activity across all short interval T2 epochs first, then calculated the GFP after that averaging.

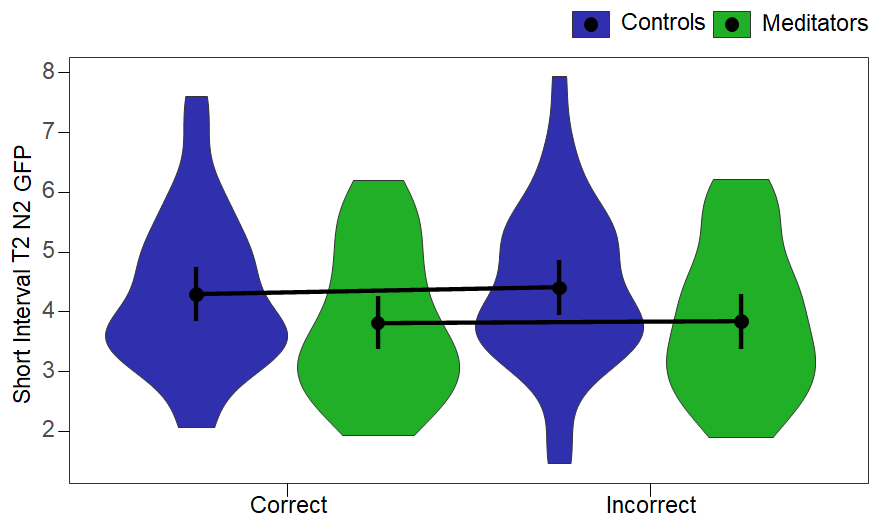

Figure S4. Estimated marginal means for posterior-N2 global field potential (GFP) to short interval T2 trials, averaged across the time period of significant difference between meditators and controls (227 to 254ms), for both groups and split by whether the T2 stimuli were responded to correctly. Note that the GFP values were extracted from each short interval T2 single epoch separately, and estimated marginal means were calculated from these single trials. As such, these values are higher than those presented for the analysis within RAGU, which averaged the activity across all short interval T2 epochs first, then calculated the GFP after that averaging.

Table S5. Estimated Marginal Means from the generalised linear mixed model for posterior-N2 global field potential (GFP) to the first target stimuli in short interval trials (SIT1) and the second target stimuli in short interval trials (SIT2), averaged across both the time-period of significant difference between meditators and controls (227 to 254ms) and averaged across groups. Note that estimates are on the response scale (so reflect a proportion of correct responses that are associated with each SIT1 and SIT2 posterior-N2 GFP value). Note the pattern indicates that regardless of the size of the posterior-N2 GFP to SIT2, smaller amplitude poster-N2 GFP to SIT1 is related to higher accuracy.

|  |  |  |  | 95% CI | |
| --- | --- | --- | --- | --- | --- |
| SIT1 posterior-N2 GFP | SIT2 posterior-N2 GFP | Estimate (SIT2 response accuracy in proportion correct) | Standard Error | Lower Bound | Upper Bound |
| 1.923 | 2.182 | 0.923 | 0.011 | 0.897 | 0.942 |
| 3.651 | 2.182 | 0.868 | 0.017 | 0.830 | 0.898 |
| 5.379 | 2.182 | 0.784 | 0.027 | 0.727 | 0.832 |
| 1.923 | 3.983 | 0.910 | 0.012 | 0.883 | 0.932 |
| 3.651 | 3.983 | 0.863 | 0.018 | 0.825 | 0.894 |
| 5.379 | 3.983 | 0.796 | 0.025 | 0.744 | 0.840 |
| 1.923 | 5.784 | 0.897 | 0.015 | 0.864 | 0.922 |
| 3.651 | 5.784 | 0.858 | 0.019 | 0.818 | 0.891 |
| 5.379 | 5.784 | 0.808 | 0.024 | 0.757 | 0.850 |

Table S6. Statistics from the generalised linear mixed model with correct/incorrect response to the second target in short interval trials (SIT2) as the outcome measure, and posterior-N2 global field potential (GFP) after short interval T1 (SIT1) and T2 (SIT2) stimuli, averaged across the time period of significant difference between meditators and controls (227 to 254ms) as well as trial number as the fixed effect variables. Significant p-values are presented in bold.

| Effect | Chi-Sq Value | p-value |
| --- | --- | --- |
| Posterior-N2 GFP SIT1 | 56.963 | **< 0.001** |
| Posterior-N2 GFP SIT2 | 13.781 | **< 0.001** |
| Trial Number | 0.938 | 0.333 |
| Group | 0.258 | 0.611 |
| Posterior-N2 GFP SIT1  *  Posterior-N2 GFP SIT2 | 14.412 | **< 0.001** |
| Posterior-N2 GFP SIT1  *  Trial Number | 0.148 | 0.701 |
| Posterior-N2 GFP SIT2  *  Trial Number | 1.138 | 0.286 |
| Posterior-N2 GFP SIT1  *  Group | 0.003 | 0.957 |
| Posterior-N2 GFP SIT2  *  Group | 1.324 | 0.250 |
| Trial Number  *  Group | 1.337 | 0.248 |
| Posterior-N2 GFP SIT1 *   Posterior-N2 GFP SIT2  *  Trial Number | 0.083 | 0.774 |
| Posterior-N2 GFP SIT1  *   Posterior-N2 GFP SIT2  *  Group | 0.150 | 0.698 |
| Posterior-N2 GFP SIT1  *  Trial Number  *  Group | 0.002 | 0.968 |
| Posterior-N2 GFP SIT2  *  Trial Number  *  Group | 4.244 | **0.039** |
| Posterior-N2 GFP SIT1  *   Posterior-N2 GFP SIT2  *  Trial Number  *  Group | 0.292 | 0.589 |

Table S7. Estimated marginal means from the generalised linear mixed model with correct/incorrect responses as the outcome measure, and posterior-N2 global field potential (GFP) after the first target stimuli in short interval trials (SIT1), averaged across the time period of significant difference between meditators and controls, and split by whether T2 was responded to correctly (227 to 254ms) as the fixed effect variable. Estimates are on the response scale. Note that the estimated marginal means showed that both groups showed better performance with low T1 posterior-N2 GFP, and that although performance improved across trials, this pattern remained the same throughout the task.

|  |  |  |  |  | 95% CI | |
| --- | --- | --- | --- | --- | --- | --- |
| posterior-N2 SIT1 GFP | Trial Number | Group | Estimate | Standard Error | Lower | Upper |
| 1.923 | 56.258 | Controls | 0.883 | 0.022 | 0.832 | 0.920 |
| 3.651 | 56.258 | Controls | 0.835 | 0.028 | 0.772 | 0.884 |
| 5.379 | 56.258 | Controls | 0.774 | 0.036 | 0.695 | 0.837 |
| 1.923 | 141.949 | Controls | 0.905 | 0.018 | 0.863 | 0.935 |
| 3.651 | 141.949 | Controls | 0.867 | 0.023 | 0.814 | 0.907 |
| 5.379 | 141.949 | Controls | 0.817 | 0.031 | 0.750 | 0.870 |
| 1.923 | 227.640 | Controls | 0.924 | 0.016 | 0.887 | 0.949 |
| 3.651 | 227.640 | Controls | 0.894 | 0.020 | 0.848 | 0.927 |
| 5.379 | 227.640 | Controls | 0.854 | 0.026 | 0.794 | 0.899 |
| 1.923 | 56.258 | Meditators | 0.891 | 0.020 | 0.844 | 0.925 |
| 3.651 | 56.258 | Meditators | 0.838 | 0.028 | 0.777 | 0.885 |
| 5.379 | 56.258 | Meditators | 0.767 | 0.037 | 0.686 | 0.832 |
| 1.923 | 141.949 | Meditators | 0.916 | 0.016 | 0.879 | 0.942 |
| 3.651 | 141.949 | Meditators | 0.879 | 0.021 | 0.830 | 0.915 |
| 5.379 | 141.949 | Meditators | 0.829 | 0.029 | 0.764 | 0.879 |
| 1.923 | 227.640 | Meditators | 0.935 | 0.013 | 0.905 | 0.957 |
| 3.651 | 227.640 | Meditators | 0.910 | 0.017 | 0.872 | 0.938 |
| 5.379 | 227.640 | Meditators | 0.877 | 0.023 | 0.824 | 0.916 |

Table S8. Estimated marginal means from the generalised linear mixed model with correct/incorrect responses as the outcome measure, and posterior-N2 global field potential (GFP) after the second target stimuli in short interval trials (SIT2), averaged across the time period of significant difference between meditators and controls, and split by whether T2 was responded to correctly (227 to 254ms) as the fixed effect variable. Estimates are on the response scale. Note that the estimated marginal means showed that to begin with, both groups showed better performance with low T2 posterior-N2 GFP. This pattern was maintained for the control group, but for the meditation group, the pattern switched so that high SIT2 posterior-N2 GFP values were associated with correct responses.

|  |  |  |  |  | 95% CI | |
| --- | --- | --- | --- | --- | --- | --- |
| posterior-N2 SIT2 GFP | Trial Number | Group | Estimate | Standard Error | Lower | Upper |
| 2.182 | 56.258 | Controls | 0.837 | 0.029 | 0.771 | 0.887 |
| 3.983 | 56.258 | Controls | 0.830 | 0.029 | 0.765 | 0.880 |
| 5.784 | 56.258 | Controls | 0.822 | 0.032 | 0.751 | 0.875 |
| 2.182 | 141.949 | Controls | 0.875 | 0.023 | 0.822 | 0.913 |
| 3.983 | 141.949 | Controls | 0.862 | 0.024 | 0.808 | 0.903 |
| 5.784 | 141.949 | Controls | 0.849 | 0.027 | 0.789 | 0.895 |
| 2.182 | 227.640 | Controls | 0.904 | 0.019 | 0.859 | 0.936 |
| 3.983 | 227.640 | Controls | 0.890 | 0.021 | 0.842 | 0.924 |
| 5.784 | 227.640 | Controls | 0.873 | 0.024 | 0.817 | 0.914 |
| 2.182 | 56.258 | Meditators | 0.842 | 0.028 | 0.780 | 0.890 |
| 3.983 | 56.258 | Meditators | 0.828 | 0.029 | 0.763 | 0.878 |
| 5.784 | 56.258 | Meditators | 0.812 | 0.033 | 0.739 | 0.868 |
| 2.182 | 141.949 | Meditators | 0.868 | 0.024 | 0.815 | 0.908 |
| 3.983 | 141.949 | Meditators | 0.874 | 0.022 | 0.823 | 0.912 |
| 5.784 | 141.949 | Meditators | 0.879 | 0.023 | 0.828 | 0.917 |
| 2.182 | 227.640 | Meditators | 0.890 | 0.021 | 0.842 | 0.925 |
| 3.983 | 227.640 | Meditators | 0.909 | 0.017 | 0.869 | 0.938 |
| 5.784 | 227.640 | Meditators | 0.925 | 0.016 | 0.887 | 0.951 |

#### Section 3c - Theta Phase Synchronisation – Validation Check

The TCT results for RMS TPS indicated topographical consistency within all groups and conditions that overlapped with the significant effects reported in our main manuscript (see Figure S5). Within our main TPS analyses, in order to maximise our statistical power, we included all participants in our analysis of TPS. However, we note that Slagter et al. (2009) excluded participants who provided fewer than 40 epochs for analysis to maximise the validity of the TPS measure. After excluding participants who provided fewer than 40 correct response and artifact-free epochs for all conditions (11 participants total excluded, 3 meditators), the same TPS RMS interaction between group, target, and interval from 117 to 295ms post stimulus was significant (p = 0.0012, η_p_^2^ = 0.1966 when averaged across the 117 to 295ms time period). Unfortunately, while it would be informative to perform a single trial analysis of TPS to assess whether higher TPS is associated with correct responses (as we have for the posterior-N2 and alpha-power), TPS is calculated as the synchronisation of theta phase to stimuli across multiple trials, so single trial analysis is not possible.

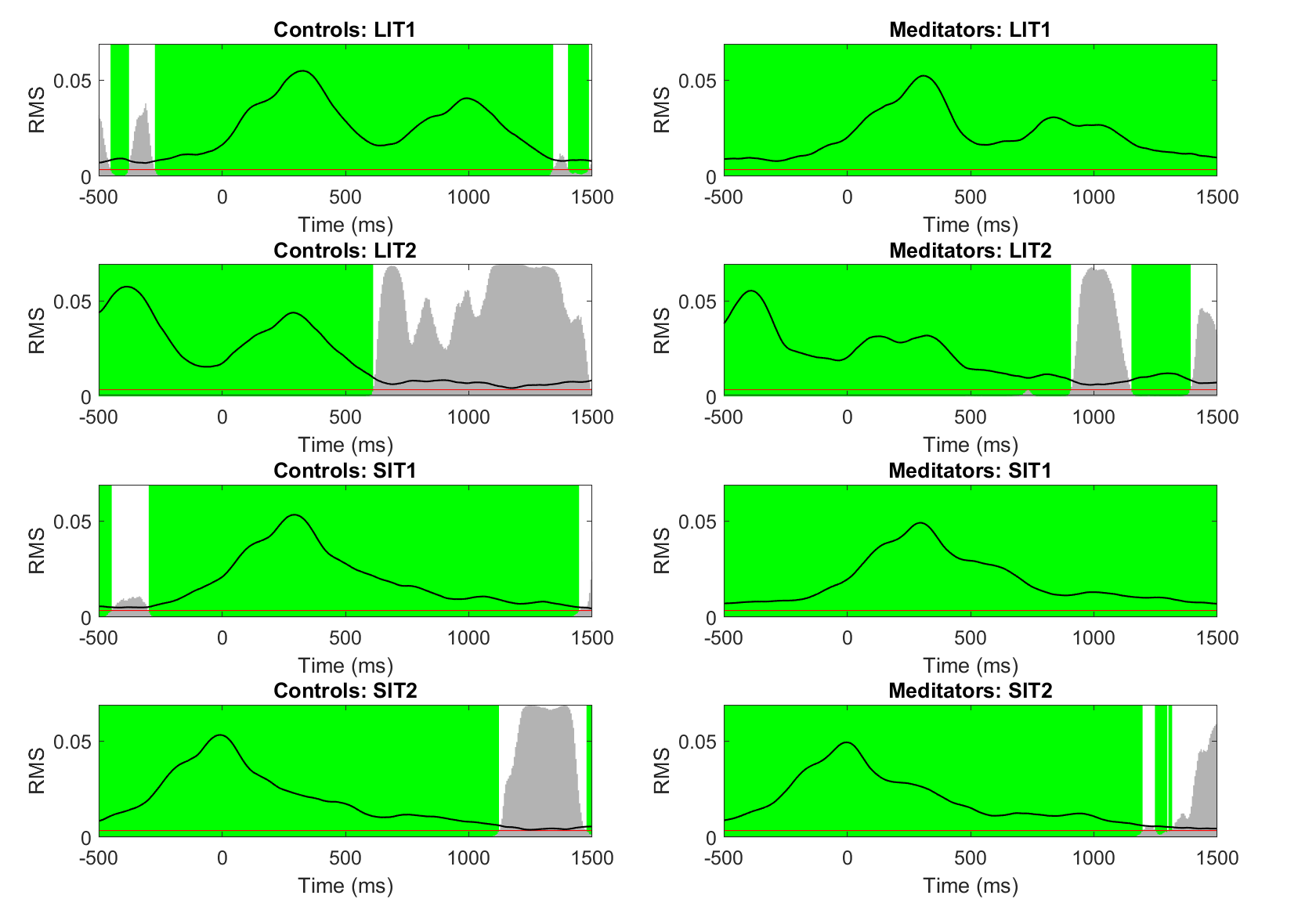

Figure S5. Topographical consistency test (TCT) results for root mean squared (RMS) theta phase synchronisation (TPS) data. The black line represents root mean squared (RMS) values across the epoch. Green periods reflect periods of significant topographical consistency within the group/condition. Grey bars reflect the p-value, with the red line at the bottom indicating the p < 0.05 level. SIT1 reflects short interval RMS TPS following the first target stimuli (T1), SIT2 reflects short interval RMS TPS following the second target stimuli (T2), LIT1 reflects long interval RMS TPS following the first target stimuli (T1), LIT2 reflects long interval RMS TPS following the second target stimuli (T2). Note the double peaks in long interval trials (indicating theta phase synchronisation to both T1 and T2 stimuli) but only the single peak for short interval trials (suggesting neural mechanisms underpinning theta phase synchronisation to T2 are not engaged, perhaps because those mechanisms were still recovering from engagement to process T1).

#### Section 3d - Theta Phase Synchronisation – Single Electrode Analysis

In addition to the RMS and TANOVA tests, which included all electrodes, we examined specific single electrodes that showed significant differences in the study by Slagter et al. (2009). In alignment with our RMS TPS test results (rather than the TANOVA results), the single electrode comparisons of TPS following T2 stimuli during the 121 to 500ms window at electrodes Fz and FC6 showed a significant interaction between group and interval (F(1,59) = 7.058, p = 0.01, η^2^G = 0.010, BFincl = 4.621, see Figure S6). Within this interaction, the meditator group showed similar TPS values following short interval T2 trials compared to their long interval T2 trials, while the control group showed reduced TPS following short interval T2 trials compared to long interval T2 trials. The meditator group also showed larger TPS values following short interval T2 trials than the control group in these electrodes. Given the RMS test showed a significant result for only part of this time period, it is likely that the effect size would have been even larger if restricted to the time window from our data rather than in replication of Slagter et al. (2009). Indeed, this was the case, with the same analysis restricted to the 117 to 295ms window showing a larger effect size for the interaction between group and interval (F(1,59) = 14.454, p < 0.001, η^2^G = 0.018, BFincl = 35.908). No other main effects or interactions involving group were significant for comparisons involving these electrodes and either time window (all p > 0.3). The effects involving group from electrode T8 during the 309 to 558ms post T2 window were also not significant, with no main effect of group (F(1,59) = 2.075, p = 0.155, η^2^G = 0.025, BFexcl = 1.542), and no interaction between group and interval (F(1,59) = 0.054, p = 0.816, η^2^G = 0.025, BFexcl = 3.819).

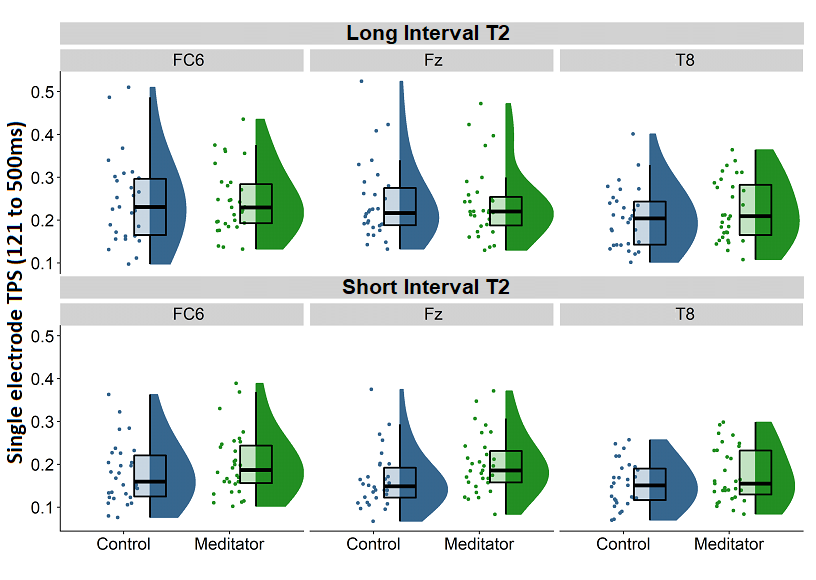

Figure S6. Theta phase synchronisation (TPS) from single electrode analyses. FC6, Fz, and T8 refer to the electrode of interest.

#### Section 3e - Alpha-power RMS Exploratory Analyses

The TCT results for RMS alpha-power indicated topographical consistency within all groups and conditions across the entire epoch (see Figure S7). Our results indicated the meditators showed a reduction in alpha-power from 475 to 685ms following T1 presentation (Figures S8 and S9). This period began 175ms after the short interval T2 stimuli were presented, so the reduction in alpha-power in the meditation group occurred while they were processing short interval T2 stimuli. To assess the likelihood of the potential explanation of the lower RMS alpha-power (from 475 to 685ms following T1 presentation) in the meditator group that was presented in our discussion, we computed the average RMS alpha-power within this time period from T1 short interval epochs. We then used the MATLAB function ‘polyfit’ to compute the slope of averaged alpha-power RMS within this time-period across trials (from the first epoch to the last epoch available) for each participant. This produced a mean slope of -9.498e-04 (SD = 0.0014), which a single sample t-test indicated was significantly different from 0 (t(60) = -5.436, p < 0.0001, 95% CI = -0.0013 to -0.0006). This demonstrated that RMS alpha-power reduced as the task progressed. An independent samples t-test indicated that the two groups did not differ in average RMS slope as the task progressed (t(59) = -0.480, p = 0.633). Despite the indication that RMS alpha-power during this time window (baseline corrected to the entire epoch) reduced as the task progressed, overall average RMS alpha-power during this time window did not correlate with short interval T2 percentage correct across all participants (r = -0.002, p = 0.985). We also noted that there was a main effect of Interval from 685 to 1050ms after T1, whereby long interval trials showed an increase in RMS alpha-power (Figures S8, S9 and S10).

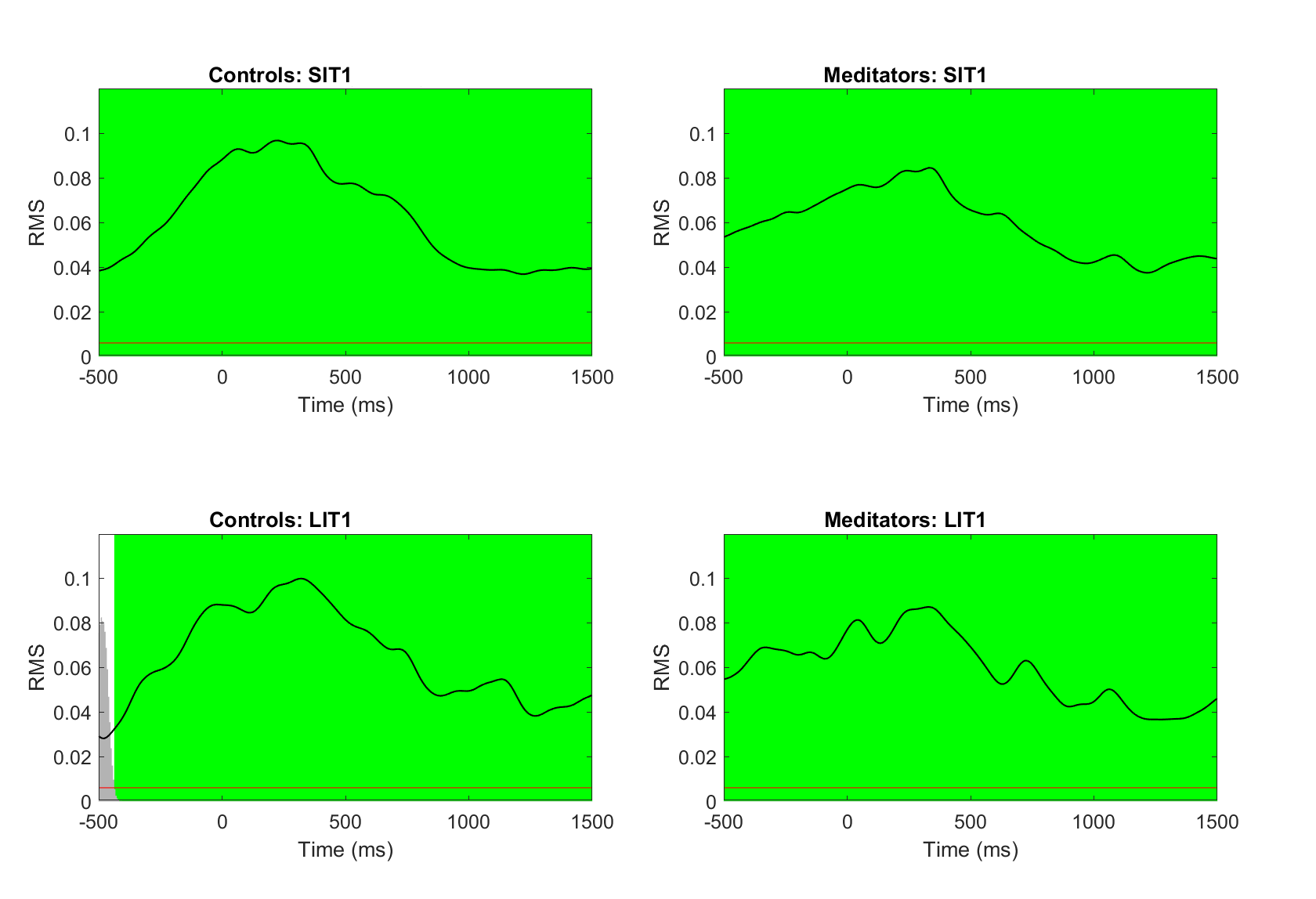

Figure S7. Topographical Consistency Test (TCT) results for root mean square (RMS) alpha-power data. The black line represents RMS values across the epoch. Green periods reflect periods of significant topographical consistency within the group/condition. Grey bars reflect the p-value, with the red line at the bottom indicating the p < 0.05 level. SIT1 reflects short interval RMS TPS following the first target stimuli (T1), LIT1 reflects long interval RMS TPS following the first target stimuli (T1).

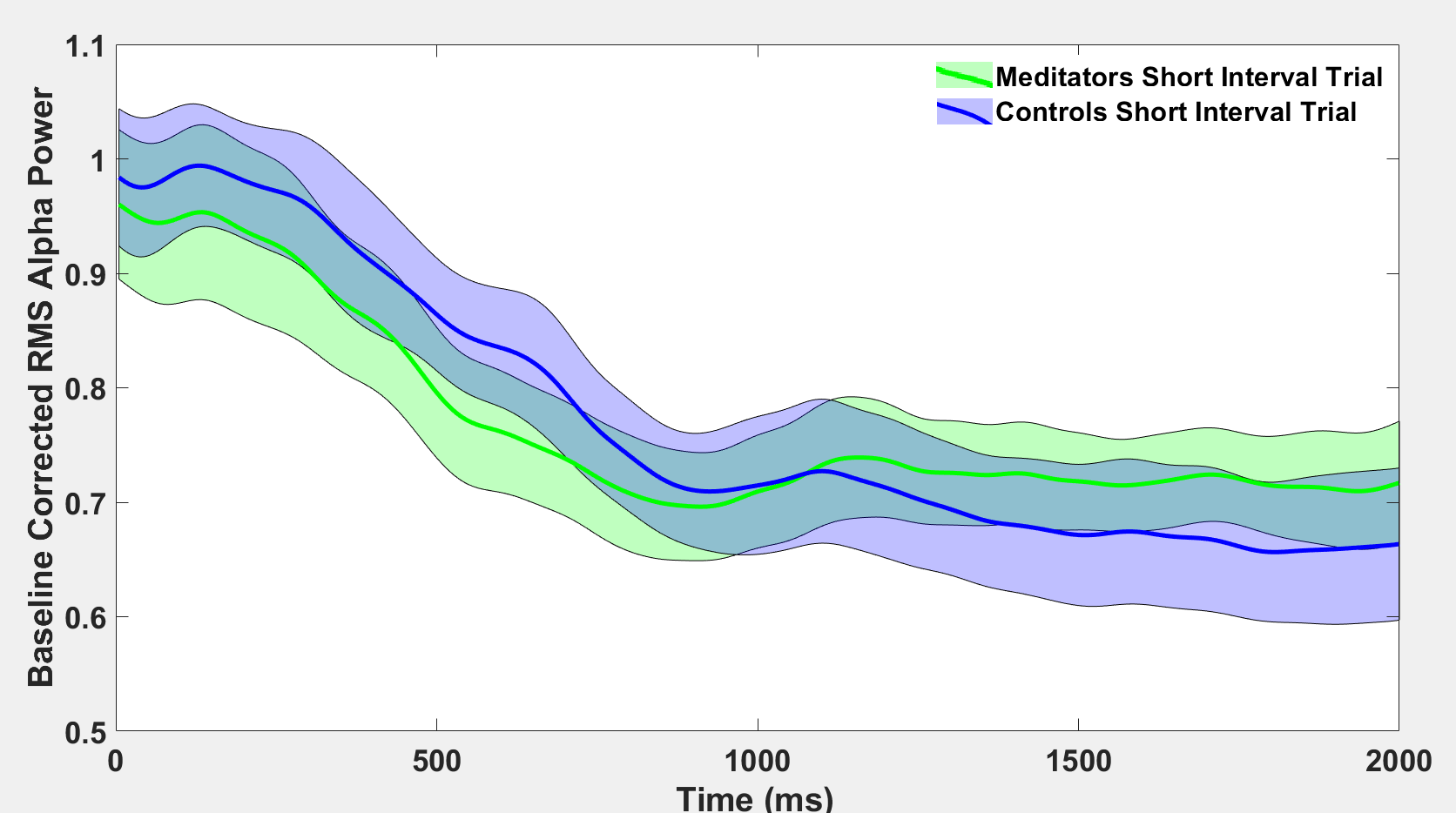

Figure S8. Baseline corrected root mean squared (RMS) alpha-power for short interval trials following the first target stimuli (T1) from each group.

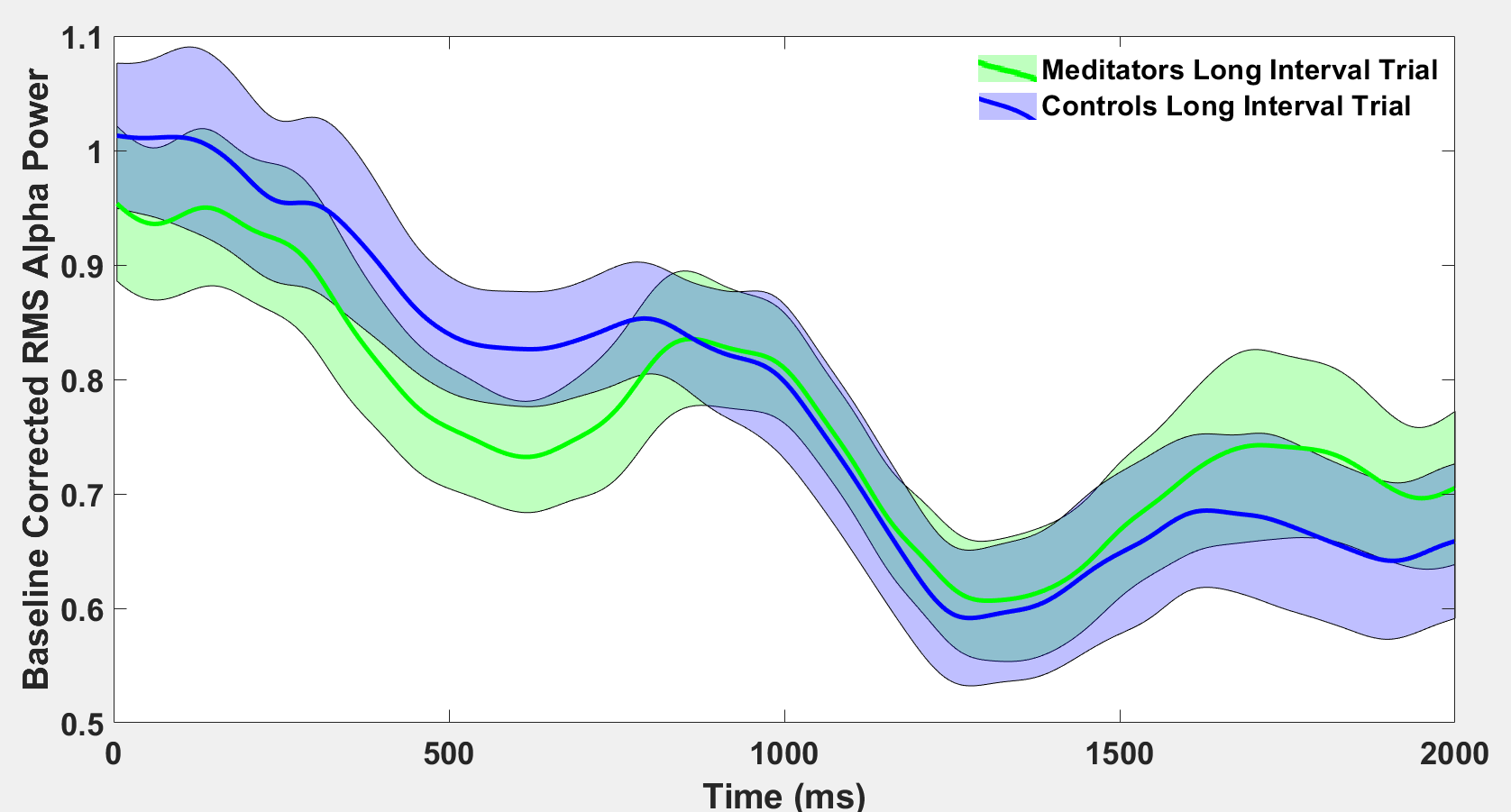

Figure S9. Baseline corrected root mean squared (RMS) alpha-power for long interval trials following the first target stimuli (T1) from each group.

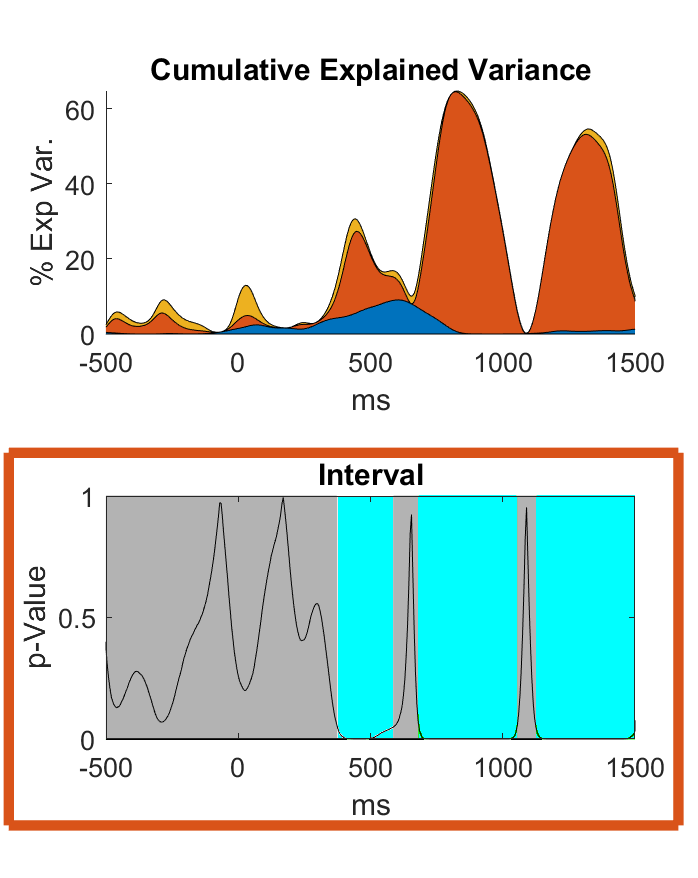

Figure S10. Top: Root mean squared (RMS) alpha-power cumulative explained variance (η_p_^2^) at each time point across the epoch by each main effect and condition, with each colour reflecting the η_p_^2^ from the effect being tested, colour coded by the effect (orange is the main effect of Interval, blue is the main effect of Group, and yellow is the interaction between Group and Interval). Bottom: p-graph for the main effect of interval. The black line reflects the p-value, the white areas reflect significant time points, and the light blue periods reflect windows where the effect passed Slagter et al.’s (2009) duration control. Note the first two time periods of significance, which Figures S8 and S9 indicate show higher alpha-power in long interval trials in the first period (perhaps reflecting inhibition of the processing of distractor stimuli), and lower alpha-power in long interval trials in the second period (perhaps reflecting the release of inhibition to process the long interval T2 stimuli).

To determine the relationship between alpha-power during these periods and correct/incorrect responses, we conducted a generalised linear mixed model (using the binomial family and logit model test, with participant as the random effect variable and group, RMS alpha-power from 475 to 685ms and also from 685 to 1050ms as the fixed effects variables, and correct/incorrect responses as the dependent variable) indicated that *incorrect* responses were associated with lower baseline corrected alpha-power RMS from 475 to 685ms following T1 presentation (ChiSq = 8.334, p = 0.004, see Figure S11 and Table S9). However, in support of theoretical perspectives that suggest alpha-power reflects inhibition of brain regions, lower RMS alpha-power in the later period (685 to 1050ms) was associated with *correct* responses in short interval T2 trials (ChiSq = 43.724, p < 0.001, see Figure S12 and Table S10), with a much stronger effect than the opposite effect reported for the earlier time-period. Additionally, we found that across all epochs from all participants, RMS alpha-power during the 475 to 685ms following T1 presentation correlated with the RMS alpha-power during the 685 to 1050ms time-period (r = 0.398, p < 0.001).

Finally, to assess whether single trial response accuracy modulated the relationship between the task-relevant RMS alpha-power from 685 to 1050ms and RMS alpha-power in the earlier time period, a linear mixed model with correct/incorrect response, RMS alpha-power from 475 to 685ms, and group as fixed effect variables, participants as the random effects variable, and alpha-power RMS from 685 to 1050ms as the dependent variable. This analysis showed that the relationship between the two RMS alpha-power time periods was stronger for the incorrect trials than the correct trials (estimated trend for correct trials = 0.268, estimated trend for incorrect trials 0.342). This result might suggest that the lower alpha-power in meditators during the earlier 475 to 685ms period might reflect a preparatory mechanism, engaging attention when attention had drifted, so that the neural activity required for successful task performance in the later 685 to 1050ms window might be more likely to be present. Note that for this linear mixed model, JASP provided an error message that the model fit was singular, so the design was reduced to exclude group, response type, and the interaction between these two as random effects. Applying this setting did not change the pattern or significance of the results.

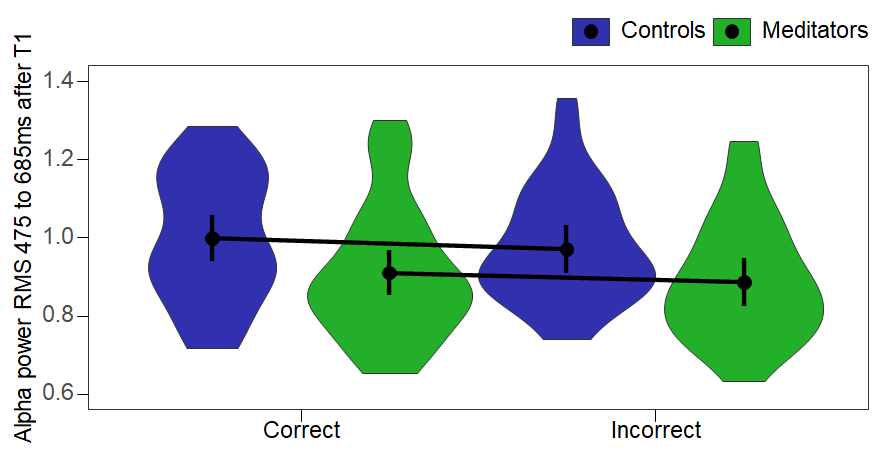

Figure S11. Estimated marginal means for baseline corrected root mean squared (RMS) alpha-power averaged across the time period of significant difference between meditators and controls (475 to 685ms), for both groups and split by whether the T2 stimuli were responded to correctly.

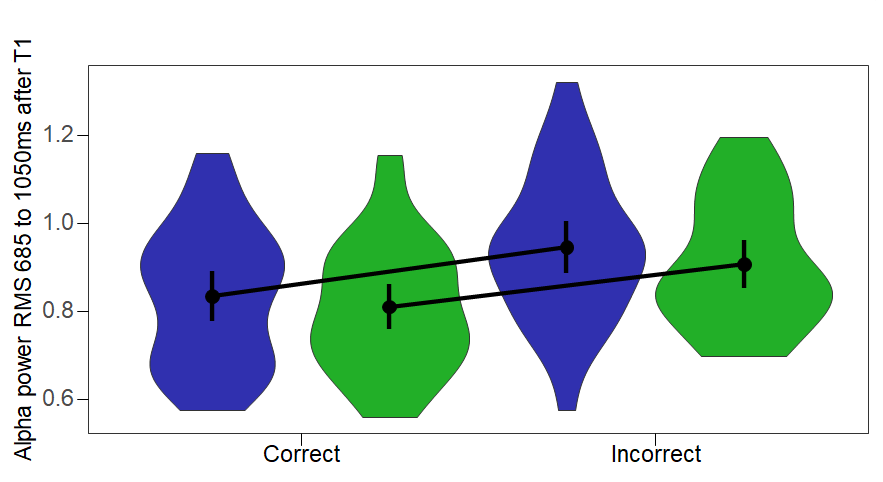

Figure S12. Estimated marginal means for baseline corrected root mean squared (RMS) alpha-power averaged across the task performance relevant time-period (685 to 1050ms), for both groups and split by whether the T2 stimuli were responded to correctly.

Table S9. Estimated Marginal Means with baseline corrected root mean square (RMS) alpha-power averaged across the time-period of significant difference between meditators and controls (475 to 685ms) as the outcome variable, and Correct/Incorrect Response and Group as the fixed-effect variables.

|  | | | | | | | | | | |
| --- | --- | --- | --- | --- | --- | --- | --- | --- | --- | --- |
|  | | | | | | | | 95% CI | | |
| Group | | Response | | Estimate | | SE | | Lower | | Upper |
| Controls |  | Correct |  | 0.998 |  | 0.030 |  | 0.940 |  | 1.057 |
| Meditators |  | Correct |  | 0.910 |  | 0.029 |  | 0.853 |  | 0.967 |
| Controls |  | Incorrect |  | 0.970 |  | 0.031 |  | 0.908 |  | 1.032 |
| Meditators |  | Incorrect |  | 0.887 |  | 0.031 |  | 0.826 |  | 0.947 |

Table S10. Estimated Marginal Means with baseline corrected root mean square (RMS) alpha-power averaged across the task performance relevant period (685 to 1050ms) as the outcome variable, and Correct/Incorrect Response and Group as the fixed-effect variables.

|  | | | | | | | | | | |
| --- | --- | --- | --- | --- | --- | --- | --- | --- | --- | --- |
|  | | | | | | | | 95% CI | | |
| Group | | Response | | Estimate | | SE | | Lower | | Upper |
| Controls |  | Correct |  | 0.836 |  | 0.029 |  | 0.779 |  | 0.892 |
| Meditators |  | Correct |  | 0.811 |  | 0.026 |  | 0.759 |  | 0.863 |
| Controls |  | Incorrect |  | 0.946 |  | 0.031 |  | 0.886 |  | 1.006 |
| Meditators |  | Incorrect |  | 0.907 |  | 0.028 |  | 0.852 |  | 0.961 |

These results support for the interpretation of alpha as an inhibitory mechanism, with alpha-power decreasing from T1 presentation to the short interval T2 processing period (with a main effect of interval showing significance from 685 to 1050ms after T1 - short interval T2 stimuli were presented at 300ms, then the alpha decrease occurred later 385ms, allowing for initial processing before task-relevant identification was likely to be engaged). In contrast, a decrease in alpha-power occurred at the same time period after T1 presentation in long interval trials (as participants could become aware at this time that a short interval T2 stimuli was not presented). Then later in the long interval trials, alpha was decreased again following T2 presentation (1125 to 1500ms, ~425ms after long interval T2 presentation). In alignment with theories of the function of alpha-power, we suspect these alpha decreases dependent upon T2 presentation timing are likely to reflect the release of inhibitory functions to allow processing of long interval T2 stimuli.

#### Section 3f - RMS Alpha Phase Synchronisation Exploratory Analyses

The TCT results for RMS APS indicated topographical consistency within all groups and conditions across the entire epoch (see Figure S13). Correlations between RMS APS averaged within the significant window (282 to 1500ms), and accuracy at detecting the second target stimuli (T2) in short interval trials indicated a significant relationship between the two variables (Pearson’s r = 0.314, p = 0.014, BF10 = 3.093), and for the correlation between APS RMS during long interval T1 trials and T2 short interval percentage correct (Pearson’s r = 0.307, p = 0.016, BF10 = 2.717) (Figure S14). Note the same pattern is present in both groups. Unfortunately, while it would be informative to perform a single trial analysis of APS to assess whether higher APS is associated with correct responses (as we have for the posterior-N2 and alpha-power), APS is calculated as the synchronisation of alpha phase to stimuli across multiple trials, so single trial analysis is not possible.

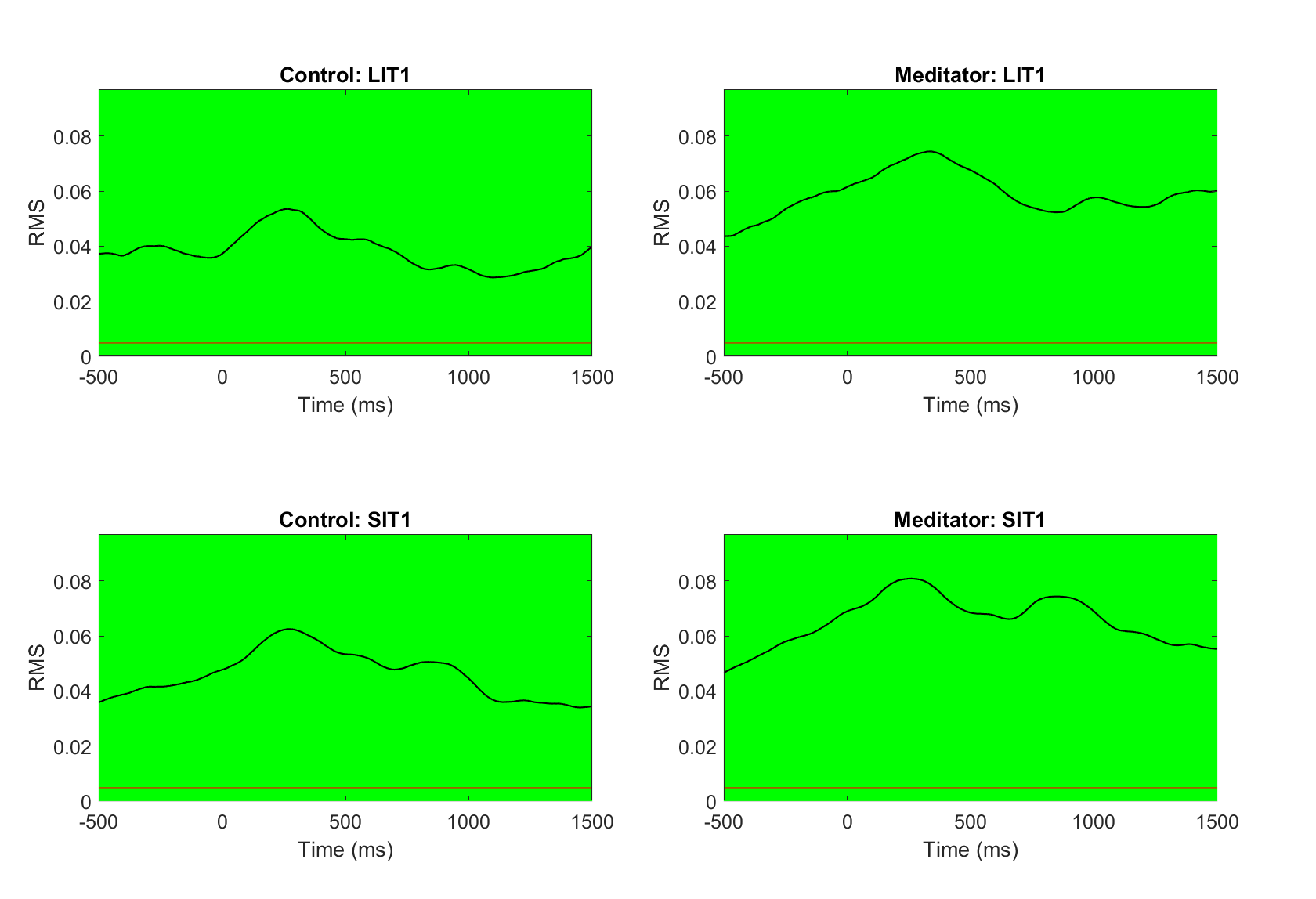

Figure S13. Topographical Consistency Test (TCT) results for Root Mean Squared (RMS) Alpha Phase Synchronisation (APS) data. The black line represents root mean squared (RMS) values across the epoch. Green periods reflect periods of significant topographical consistency within the group/condition. Grey bars reflect the p-value, with the red line at the bottom indicating the p < 0.05 level. SIT1 reflects short interval RMS TPS following the first target stimuli (T1), LIT1 reflects long interval RMS TPS following the first target stimuli (T1).

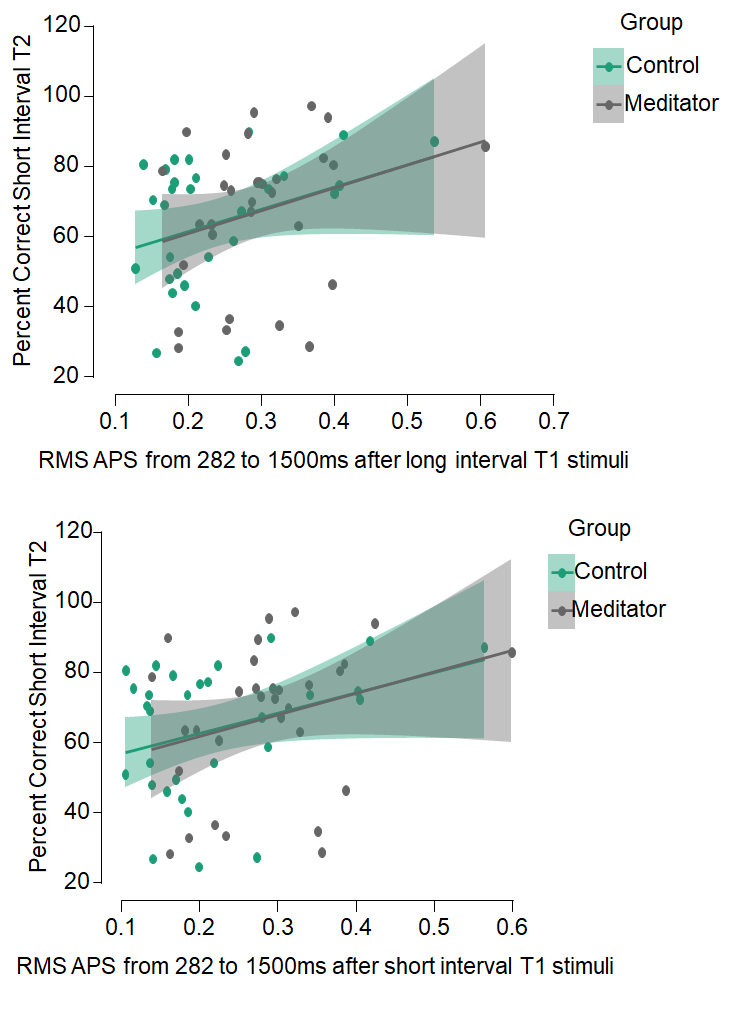

Figure S14. Scatterplots depicting the correlations between root mean squared (RMS) alpha phase synchronisation (APS) averaged within the significant window (282 to 1500ms), and accuracy at detecting the second target stimuli (T2) in short interval trials, split by group.

### Supplementary Materials Section 4 - Discussion Points

#### Potential explanations for the null result - Contextual factors

While our results suggest differences in neural activity in meditators that align with improved attention function, the groups did not differ in task performance, or in our primary analyses. The context in which our sample was studied may be a possible explanation for this null result. The current study deliberately did not ask meditators to practice meditation before or during the task to determine whether previously observed meditation-related behavioural and neural changes during the AB reflect trait changes resulting from long-term mindfulness meditation practice. Some previous studies have found a reduced AB due when mindfulness meditation was deliberately practised to induce a meditative state prior to the task (May et al., 2011). The alteration of the AB effect by meditation also seems to be specific to certain types of meditative states - only the induction of open-monitoring meditation resulted in a reduction in the AB phenomenon, not focused attention (van Vugt & Slagter, 2014). Thus, the behavioural effects of meditation in the AB task may be dependent on both a meditation-induced mindful state, and particular types of meditative practices. This interpretation is supported by studies of attention tasks other than the AB, which have shown improved attention-related performance only when a state of meditation was induced prior to the task (Cahn & Polich, 2006; Lutz et al., 2009; Sarang & Telles, 2006) or during the task (Atchley et al., 2016; Delgado-Pastor et al., 2013). However, the specificity of AB effects on meditation states is not entirely consistent, with some research having shown that meditators displayed a reduced AB without the active induction of a meditative state (Fabio & Towey, 2018; van Leeuwen et al., 2009). Alternatively, it may be that the AB task used in the current study was not sensitive enough to detect potentially enhanced attention in our meditation sample, compared to our also highly educated control sample. Our short interval trials had a 300ms separation between T1 and T2 stimuli, which replicated Slagter et al. (2007, 2009), but was longer than some research (which also tested a range of durations between T1 and T2 rather than just 300ms and 700ms). The attention blink effect is suggested to be the largest with a delay of 200ms between T1 and T2, and other modifications of the task can also make it more difficult (Nieuwenstein et al., 2009). However, given the Bayesian analysis provided strong evidence against behavioural differences, if task sensitivity was an explanation for our null result, potential differences must be similar to an on/off switch (with our current task parameters not eliciting differences at all), rather than a spectrum of increasing difference between the groups. It is also not clear why other research has shown behavioural differences using the same short interval delay. Further research is required to understand how factors such as the state of meditation, an individual’s baseline trait mindfulness, and the type of mindfulness meditation might affect attention blink performance, as well as whether alternative task parameters could be more sensitive to potential differences.

#### Characteristics of the Meditators

Another explanation for the null findings of the current study might relate to the characteristics of the meditators. The current sample had an approximate average estimated 2,376 hours of meditation lifetime experience (calculated from the average minutes meditating per session multiplied by the average frequency of meditation per week multiplied by total years of meditation reported × 52/60). While the fluctuation of meditation practice across the lifespan will likely render this estimation inaccurate for a number of participants (Hasenkamp & Barsalou, 2012), this estimate is comparable to Slagter et al. (2007), whose sample had an average of 2,967 hours of lifetime meditation experience. However, meditation research often includes considerably more experienced meditating participants; Cahn et al. (2013) estimated an average of 13,520 hours for their sample, and Delgado-Pastor et al. (2013) estimated approximately 5,850 hours for their sample. The null behavioural results of the present study may be explained by such dissimilarity between the meditation experience of current participants and the experience of participants in previous research. However, it is also worth noting that a considerable amount of research has demonstrated differences in neural activity after an 8-week Mindfulness Based Stress Reduction or Mindfulness Based Cognitive Therapy course, suggesting that the meditation experience in the current sample should have been sufficient to detect neural changes as a result of the meditation practice if they were present (Sevinc et al., 2018; Hatchard et al., 2017; van der Velden et al., 2015; Gotink et al., 2016). Further, previous research has shown that not just meditation experience, but even simply higher trait mindfulness in a population naïve to meditation practice associated with a reduction in an emotional AB (Makowski et al., 2019).

The age of participants may play a role in the null findings of the present study. The average ages of the current sample of meditators and controls used in this study were 35.77 and 31.63 years, respectively. Whilst our analysis showed this age difference was not significant, and our behavioural analysis co-varying for age showed the same results as the original analysis, it may be that the meditators in our study were too young to exhibit detectable behavioural and neurophysiological changes – perhaps meditation protects against age-related decline in AB performance, and our young meditation group had not aged enough to show this effect. However, previous research has shown that older long-term meditators have a reduced AB compared to both age-matched controls *and* a younger control group (van Leeuwen et al., 2009). As such, we think that age effects are unlikely to explain our current results.

#### Different analysis designs and techniques

While significant differences in both brain activity and behaviour were found by Slagter et al. (2009), their meditation group showed a change from pre-retreat to post-retreat, with improved attention blink performance in short interval trials from pre to post, altered P3b amplitudes to T1 and theta phase synchronisation to T2. However, direct comparisons between the groups at only the post time point (which are analogous to the analyses performed in the current study) were not performed by Slagter et al. (2009). As such, we cannot determine whether they would have been significant. Although our results do not align with theirs, the different experimental designs may suggest there is not a direct conflict between the results of the two studies. Additionally, the study design used by Slagter et al. (2009) and other studies showing differences in behaviour associated with meditation, for example, Wang et al. (2021), had an additional element not present in our study - all participants performed the attentional blink task twice. As such, their design cannot rule out the possibility that their meditation group showed altered learning of the attentional blink task rather than simply altered attentional blink performance. That is, perhaps MM is not associated with a generalised increase in the equal distribution of attention across stimuli, leading to better performance in the attentional blink task, but rather an increased ability to learn from previous experience with a task, and as a result increased performance in a previously practised task following meditation practice.

Another possibility is that brain changes associated with meditation are specific to the stage of meditation. The meditators in our study were relatively experienced, but unlikely to be as well practised as those in the study by Slagter et al. (2009). It may be that meditation elicits changes in brain activity that are qualitatively different as the meditation experience increases. Perhaps only the more superficial changes to brain activity were demonstrated in our participants, which could offer a potential explanation for why we detected some changes in brain activity associated with the attentional blink task (theta phase synchronisation), but these changes only partially overlapped with the time windows detected by Slagter et al. (2007, 2009).

Further to these points, it is worth noting that the data pre-processing and analysis techniques used in the current study differed from those used by Slagter et al. (2007, 2009). Given developments in EEG pre-processing techniques over the last decade, the data-cleaning techniques we used are more robust than those used by Slagter et al. (2007, 2009). They have been shown to reduce more of the non-neural artifacts that can confound comparisons of neural activity, while still preserving neural signals (Bailey et al. 2022a, 2022b). Our exploratory statistical comparisons were also conducted without selecting time periods of interest to average across, and all our analyses were performed without having to focus on specific electrodes of interest. Instead, we used global field potential and global dissimilarity distribution statistical tests, which include all electrodes and time points in the analyses, while applying multiple comparison controls derived from the data permutations. In our view, using approaches that require no arbitrary decisions about which time periods or electrodes to analyse is more robust against potential experimenter effects. These methods reduce both the risk of the selection of time periods to analyse after visual inspection reveals where differences are likely, which inflates the false positive rate (Kilner, 2013), and the risk of missing a significant result by selection time windows or electrodes of focus that miss potential significant effects.

Finally, in addition to the points noted in our main discussion, one further potential explanation of the lack of TPS finding when restricted to T2, but the positive result from the interaction, including all conditions, is that the TPS values might show high enough variance within groups that adding the other conditions to the comparison provides a baseline, in which context the interaction effect involving short interval T2 against long interval T2 stimuli became apparent.

#### Supplementary Strengths and limitations

Another limitation of this study was that meditators and non-meditators were recruited based on self-reported practice rather than controlled adherence to a specific practice for a course of time. This approach is limited in that it relies on the fallibility of an individual's memory and is vulnerable to social-desirability effects (Grimm, 2010). This lack of standardisation may restrict the ability to draw conclusions regarding a specific quantity of meditation practice.

It may be that our study was underpowered to detect main effect differences in TPS when the analysis was restricted to just the short interval T2 epochs. However, our sample size was approximately twice the size of Slagter et al. (2009), and after an average of 6.44 years of meditation experience, and a high current average practice time of 7.1 hours per week, even if an effect size could be detected with a larger sample, it is not clear that the effect would have a meaningful interpretation. It would be too small and require too much mindfulness practice for clinical application, and would also be too small to be used as a neurophysiological marker in future research, particularly when other tasks have produced much larger effect sizes.

One unexpected strength of the study was the ability within a partial replication attempt to be robust to minor variations in the effects from the original study. In particular, while our data replicated the finding from Slagter et al. (2009) that theta was synchronised to short interval T2 stimuli in meditators more than healthy controls, the effect was only apparent after inspecting the exploratory analysis that included all time points in the epoch following the T2 stimuli. When restricted to the exact time window Slagter et al. (2009) found to be significant, their results did not replicate, despite the fact that when examining just the early part of the same time period, there was very strong Bayesian evidence for a difference between the two groups. This highlights the usefulness of using analysis techniques that can still detect replication effects, despite variation in the effect within a different sample. This “shifted effect” in a replication sample may even go some way towards explaining the current replication crisis in psychology, and we would encourage future replication attempts to explore their results for non-exact replications, which still may support the initial finding.

#### Further Considerations

In addition to our discussion in the main manuscript, there are a number of further points that can be inferred from our results, and the combination of our study and the results reported by Slagter et al. (2007 and 2009). Firstly, Slagter et al. (2007) suggested that T1 processing, reflected by the P3b, was reduced 50ms after the presentation of T2 stimuli in the meditation group compared to the non-meditation group. However, our results showed the meditators produced smaller posterior-N2 responses in response to T1 stimuli (compared to T2 stimuli) prior to the onset of the short interval T2 stimuli. Slagter et al. (2007) suggested that a reduced P3b amplitude to T1 in meditators was not due to processing being interrupted by T2, as the effect occurred in both long interval trials and trials in which T2 was not present at all. Our results provide further support to this conjecture, demonstrating altered neural activity that precedes the presentation of T2.

Secondly, in addition to the relationship between posterior-N2 amplitude following T1 and individual correct or incorrect responses to T2, within the single trial analysis, the relationship between posterior-N2 amplitude following T2, trial number and individual trial response accuracy differed between the groups. To begin with, both meditators and controls were less likely to identify T2 stimuli if their posterior-N2 GFP was high. Controls showed the same pattern right through the task. However, by the end of the task, this pattern switched for the meditators, so that they were more likely to identify T2 targets when they showed high posterior-N2 GFP values. This suggests that the relationship between T2 posterior-N2 GFP and response accuracy is dependent on another, unidentified neural activity, such that meditators learnt to engage this unidentified neural mechanism as the task went on, allowing them to translate the higher posterior-N2 GFP values (reflecting putative attentional engagement mechanisms) into improved performance. It is also worth noting that our results indicate that differences between meditators and controls in the posterior-N2 were not due to differences in the distribution of brain activity across regions, but rather differences in the overall amplitude of neural activity (generated by the same brain regions for both groups).

Third, considering Slagter’s (2009) short interval T2 TPS results were correlated with behaviour, it could be considered strange that in our study the only TPS condition that was not correlated with percentage correct in response to short interval T2 trials was the short interval T2 TPS. We believe this might reflect a more general relationship between a participant’s ability to synchronise their theta oscillations to the onset of a stimulus and their ability to perceive and accurately respond to rapidly presented, difficult-to-perceive stimuli under scarce attentional resource situations, rather than a specific relationship between theta synchronisation to specific stimuli and responses to those specific stimuli. Indeed, it may be that other mechanisms in addition to theta synchronisation to stimuli onset are partially responsible (and perhaps also related to theta synchronisation) for a participant’s ability to perceive short interval T2 stimuli, and that the increased reliance on these putative mechanisms adds variance to the potential relationship between theta oscillation synchronisation to specific short interval T2 stimuli and accuracy of perception and response to those stimuli, reducing the direct relationship between theta synchronisation and behaviour to non-significance. It is also worth noting that the theta synchronisation result appears to be driven by an overall higher amount of theta synchronisation in meditators compared to controls, while both groups showed the same distribution of brain activity across the scalp (suggesting the same brain regions were engaged in both groups). It may be worth noting that the more equal distribution of TPS between T1 and T2 (and between short and long interval T2 stimuli) in meditators (and also the higher APS in meditators) could be considered to be aligned with predictive coding perspectives on the effects of mindfulness meditation. The increased synchrony perhaps reflects a mechanism to enable increased precision of processing of the posterior evidence (visual stimuli) in the meditation group (Manjalay et al. 2020, Verdonk and Trousselard, 2021, Lutz et al. 2019, Laukkonen and Slagter, 2021).
